# Orthogonal and Robust Analytics Enable Reproducible and Scalable Manufacturing of High Purity Extracellular Vesicles Derived from Mesenchymal Stromal Cells

**DOI:** 10.1101/2025.08.26.672476

**Authors:** Mehdi Dehghani, Meng Chai, Nazanin Talebloo, Molly Morrissey, Andrew Larey, Yuriy Kim, Camille Trinidad, Deepika Joshi, Yuxuan Wu, Yichen Zhong, Ran Cheng, Yisheng Lu, Behnaz Lahooti, Mohammad Islam, Zheng Zhao, Jordan Speidel, Paul Keselman, Prabuddha Mukherjee, Martha Mayo, Eric Black, David Splan, Namitha Haridas, Paul Marks, Thomas Gaborski, John Chon, Ethan Kishimori

## Abstract

The development of extracellular vesicle (EV) based therapeutics requires robust analytical assays and scalable downstream processing (DSP) strategies to ensure product quality and reproducibility. In this study, we established an analytical toolbox comprising scattering and fluorescence mode nanoparticle tracking analysis (NTA), fluorescence-based flow cytometry (Fl-FC), and multi-detector analytical size exclusion chromatography. Liposomes, selected for their physicochemical similarity to EVs, were used as reference materials to optimize assay parameters, fluorescence labeling conditions, and dynamic range, minimizing artifacts such as photobleaching and masking effects. Using these optimized analytical tools, we designed a scalable DSP workflow for human bone marrow mesenchymal stromal cell (hBM-MSC) derived EVs, incorporating clarification, tangential flow filtration (TFF), ion exchange chromatography (IEX), buffer exchange, and sterile filtration. IEX chromatography resulted in the elution of two cell-secreted populations, with similar scattering signals while eluate 1 showed approximately 30 times higher absorbance signal compared to eluate 2. Transmission electron microscopy revealed that eluate 1 contained non-vesicular extracellular particles (NVEPs), and eluate 2 was enriched in EVs and showed higher expression of positive markers such as CD81 and CD73 using Simple Western. Additionally, the two IEX eluates showed different proteomic and lipidomic profiles. Then the analytical toolbox was utilized to monitor the DSP process in terms of particle recovery and impurity removal throughout the process determining the high purity level of the final EV preparation. Together, these results demonstrate that orthogonal analytics coupled to a scalable DSP yield reproducible MSC-EV preparations while depleting commonly co-isolated NVEPs. This practical framework advances process analytics of MSC-EV manufacturing and supports reporting of identity, purity, and function aligned with existing guidelines in the field.

**Graphical Abstract:** 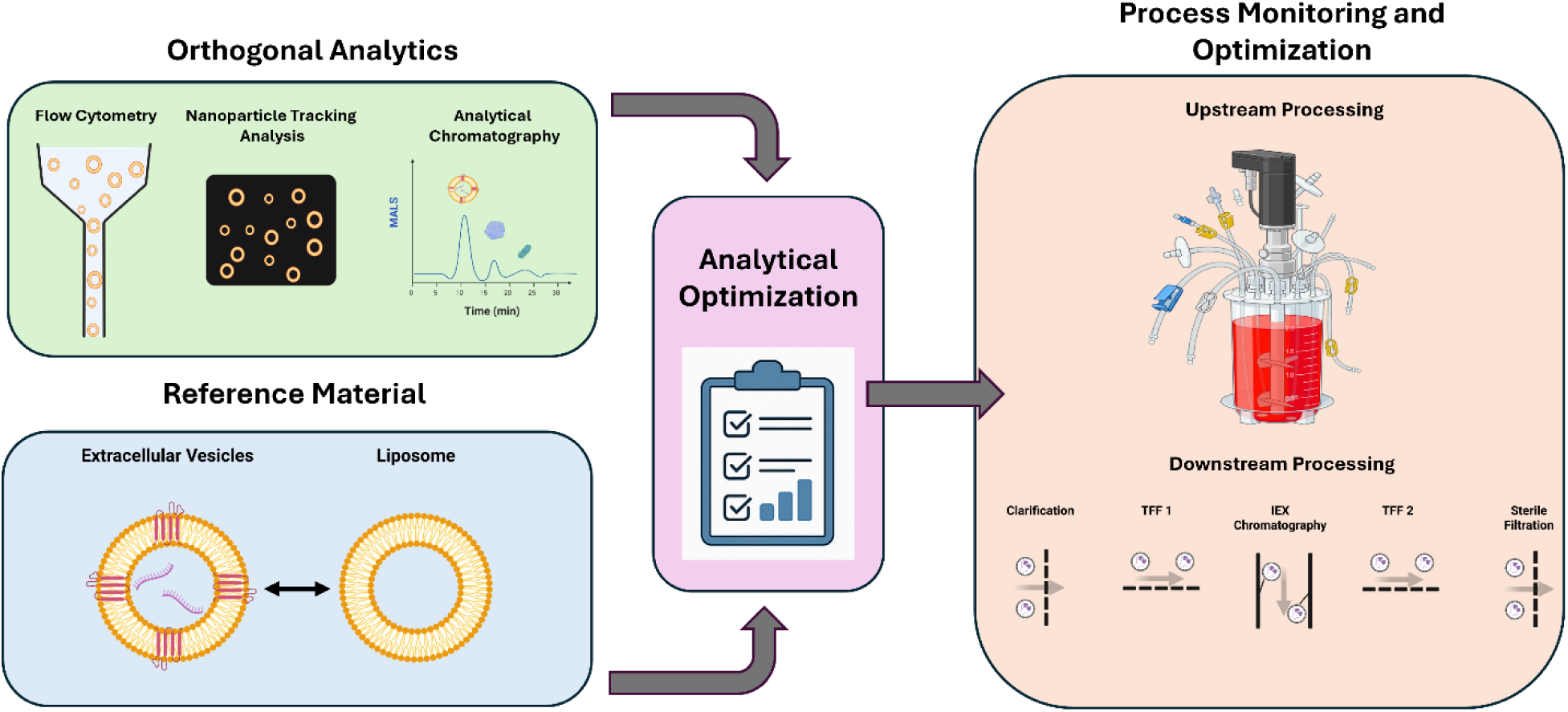

Liposomes were used as reference materials to optimize single particle analytics as well as multi detector analytical chromatography to assess particle recovery and impurity removal. This optimized analytical toolbox enabled accurate monitoring of downstream processing steps, establishing a robust workflow for reproducible and scalable mesenchymal stromal derived extracellular vesicles bioprocessing.

## Introduction

Extracellular vesicles are lipid bilayer-enclosed nanoparticles produced by nearly all cell types and can be found in most biofluids as well as cell conditioned medium (Couch *et al*., 2021; Herrmann *et al*., 2021). They play a crucial role in cell-cell communication by transporting functional cargos such as proteins, lipids and nucleic acids (Couch *et al*., 2021; Hourigan *et al*., 2025). Hence, over the past decade, EVs have emerged as diagnostic tools, therapeutic agents and drug delivery systems in preclinical and clinical settings (Escudé Martinez De Castilla *et al*., 2021; Hourigan *et al*., 2025; Tenchov *et al*., 2022). Mesenchymal Stromal Cell (MSC) derived EVs have shown promise as next generation cell-free therapeutics due to their anti-inflammatory, immunomodulatory and regenerative properties (Giebel and Lim, 2025; Kou *et al*., 2022; Muralikumar *et al*., 2021). Compared to cell-based therapies, EV-based approaches offer safety advantages, including reduced risks of tumorigenicity and unintended differentiation (Gimona *et al*., 2021; Gobin *et al*., 2021).

In MSC-EV biomanufacturing, upstream processing (USP) includes all the steps prior to harvesting or collecting the conditioned medium containing EVs such as cell isolation, cell banking, and cell expansion (Costa *et al*., 2023; Giebel and Lim, 2025; Paganini *et al*., 2019; Salvador *et al*., 2022). Whereas, the downstream processing (DSP) consists of multiple unit operations for purification, enrichment, concentration and formulation of final EV product (Adlerz *et al*., 2020; Paganini *et al*., 2019; Staubach *et al*., 2021). In EV biomanufacturing, challenges related to analytical reproducibility, process scalability, and lack of standardization impede the clinical translation and commercial development of MSC-derived EVs (Gimona *et al*., 2021; Witwer *et al*., 2019). Sensitive, optimized and reproducible analytical techniques are essential to establish and optimize a robust DSP by monitoring EV recovery and purity throughout EV production pipeline from research and development to clinical translation and scaled commercialization (Brezgin *et al*., 2023; Ng *et al*., 2019; Salvador *et al*., 2022). Despite the development of a wide range of analytical techniques, they often suffer from lack of standard references and non-optimized operating parameters and assays leading to inconsistent results across studies and laboratories (Cook *et al*., 2023; Nieuwland *et al*., 2022; Welsh *et al*., 2023).

In the first phase of this study, we established an analytical toolbox containing three techniques: nanoparticle tracking analysis, fluorescence-based flow cytometry and analytical size exclusion chromatography. NTA and Fl-FC have been widely used in the EV field for concentration measurement, size determination and protein profiling (Brealey *et al*., 2024a; Dlugolecka *et al*., 2021; Kobayashi *et al*., 2024; Lees *et al*., 2022; Midekessa *et al*., 2021). We aimed to optimize our analytical assays using liposomes as synthetic reference material for EVs. Liposomes closely mimic the size profile, lipid bilayer structure and refractive index of EVs, while remaining free from impurities and non-vesicular extracellular particles (NVEPs) (Welsh *et al*., 2020a, 2020b). We observed that post assay optimization using liposomes, orthogonal single particle analysis techniques led to comparable concentration measurements and further showed reproducibility across three instruments. However, single particle analysis techniques are primarily selective for EVs and become less reliable when analyzing crude or unpurified samples due to the presence of impurities *(Brealey et al*., 2024a; Kobayashi *et al*., 2024). To assess sample purity and enable effective monitoring and optimization of downstream processing (DSP), we incorporated analytical size exclusion chromatography (SEC) coupled with in-line multi-angle light scattering (MALS), ultraviolet (UV), and fluorescence (FL) detection (Kaddour *et al*., 2021; Normak *et al*., 2023a; Troxell *et al*., 2023). The SEC column separates different molecules based on size, while real time monitoring of MALS, absorbance and fluorescence allows semi-quantitative analysis of EVs and impurities present in the sample (Nishimura *et al*., 2024; Normak *et al*., 2023b). To confirm, we used liposomes and bovine serum albumin (BSA) as a crude EVs sample model and determined the elution profile of EV size particles and contaminants via analytical chromatography coupled with an SEC column.

In the second phase of this study, we focused on optimizing the DSP of MSC derived EVs produced in a bioreactor using the analytical toolbox. Throughout the DSP, we utilized traditional bioprocessing techniques such as clarification filters, tangential flow filtration (TFF), ion exchange chromatography (IEX) and sterile filtration (Busatto *et al*., 2018; Heath *et al*., 2018; Koch *et al*., 2023; Liu *et al*., 2025; McNamara *et al*., 2018; Staubach *et al*., 2021). These techniques have been widely deployed for different therapeutic modalities such as viral vectors and monoclonal antibodies as well as EVs (Adlerz *et al*., 2020; Matte *et al*, 2022; Nadar *et al*., 2021; Ng *et al*., 2019; Paganini *et al*., 2019; Staubach *et al*., 2021). Using IEX equipped with RT-MALS detector, we found two eluates; one enriched in vesicular structures and the other lacked vesicular components in electron microscopy (EM), while both eluates were undistinguishable in scattering characteristics using dynamic light scattering (DLS), NTA and inline MALS in analytical chromatography. We further investigated the identity of these two eluates using comprehensive and orthogonal analytics including capillary electrophoresis-based immunoassay (Simple Western), super resolution microscopy, proteomics, lipidomics, nucleic acids analysis. Finally, we demonstrated the reproducibility of the optimized DSP across independent runs and furthermore proved the scalability of the established process at larger volumes.

In conclusion, this study presents a framework for the optimization of EV analytical assays and their incorporation into developing optimized and scalable downstream processing capable of depletion of NVEPs from final EV preparations. Our findings provide novel insights into final EV product purity and recovery, allowing the researchers to improve the rigor and reproducibility of EV biomanufacturing following the recommendations of the guidelines in the field.

## Materials and Methods

### Nanoparticle tracking analysis (NTA)

500 µL of prepared samples were injected into the sample chamber of an NS300 NanoSight instrument (Malvern Panalytical Ltd.). NTA measurements were conducted using an embedded green laser (532 nm) at 45 mW along with an sCMOS camera. Unless otherwise stated, five recordings of 30 sec videos were taken per sample at camera level 16 and screen gain of 1 using NTA software version 3.4. For each measurement, videos were captured under the following conditions: cell temperature: 25°C; Syringe speed unit: 35. After capture, the videos were analyzed by the software with a detection threshold of 5, and screen gain of 10. Advanced settings: Blur, Maximum jump distance, Min track length were set at auto. At least three independent replicates of each sample were tested, and PBS was used as the dilution buffer.

### Fluorescence-based Flow Cytometry (Fl-FC)

For fluorescence-based flow cytometry (Fl-FC), a Virus Counter® 3100 was used. This instrument is designed for the enumeration of viruses and other nanoparticles of virus size. Fluorescently labelled samples (liposomes and EVs) were first tested on Fl-NTA and then diluted 10 times and ran directly on the instrument. For each test, 300 µL of the prepared liposome or EV sample was run through the flow cell of the instrument equipped with a 532 nm laser line at 150 mW laser power. Three independent replicates of each sample were tested for 60 seconds each following the manufacturer’s software manual (Sartorius).

### Fluorescence labeling Optimization

Liposome samples (Cellsome®-Exosome) were purchased from Encapsula Nano Sciences with PC: Cholesterol (70:30 molar ratio) at 50 nm (S-Lipo), 100 nm (M-lipo) and 200 nm (L-Lipo). Liposomes were serially diluted with PBS to the dynamic range of NTA instruments and tested first to determine the concentration in scattering mode.

CellMask Orange (CMO), a plasma membrane dye (Thermo Fisher) was serially diluted from stock (5 mg/mL) to: 25, 12.5, 6.25, 3.125, 1.56 and 0.78 µg/mL. Each dye concentration was then added to the sample or PBS (dye alone control) in a 1:100 ratio, i.e., 5 µL of dye in 500 µL of sample or PBS. The liposome samples (∼1× 10^9^ particles/mL) and controls were then tested on the plate reader (SpectraMax iD3 Multi-Mode Microplate Detection Platform, Molecular Devices) for spectral fluorescence detection. After incubation for 30 minutes, samples were tested on Fl-NTA and further diluted 10 times (1:10 ratio) for testing on Fl-FC.

To determine the highest liposome concentration that can be effectively labelled with optimal CMO concentration (5 µg/mL), the liposome samples were then serially diluted to ∼ 5 × 10^8^, ∼1 × 10^9^, ∼ 5 × 10^9^, ∼ 1 × 10^10^, ∼ 5 × 10^10^, ∼ 1 × 10^11^ particles/mL and then labeled with 5 µg/mL and 50 µg/mL of CMO. The labeled samples were then diluted accordingly to the dynamic range of both NTA and Fl-FC and tested.

### Size Distribution Analysis

#### Cryo-electron microscopy (Cryo-EM) Imaging of Liposome Samples

Prior to grid preparation, the samples S-Lipo and M-Lipo were diluted 10-fold with PBS (supplied by NanoImaging Services), while L-Liposome was diluted 25-fold with PBS. Each sample was preserved in vitrified ice supported by holey carbon films on 400-mesh copper grids. Each sample was prepared by applying a 3 μL drop of sample suspension to a cleaned grid, blotted away with filter paper, and immediately followed with vitrification in liquid ethane. Grids were stored under liquid nitrogen until transferred to the electron microscope for imaging. Electron microscopy was performed using a Thermo Fisher Scientific Glacios Cryo Transmission Electron Microscope (Cryo-TEM) operated at 200kV and equipped with a Falcon 4 direct electron detector. Vitreous ice grids were clipped into cartridges, transferred into a cassette and then into the Glacios autoloader, all while maintaining the grids at cryogenic temperature (below −170°C). Automated data-collection was carried out using Leginon software (Cheng *et al*., 2021; Suloway *et al*., 2005), where high magnification movies are acquired by selecting targets at a lower magnification. Images of each grid were acquired at multiple scales to assess the overall distribution of the specimen. After identifying potentially suitable target areas for imaging at lower magnifications, high magnification images were acquired at nominal magnifications of 73,000x (0.204 nm/pixel) and 28,000x (0.512 nm/pixel). The images were acquired at a nominal underfocus of −7.0 μm to −4.5 μm and electron doses of ∼2-12 e-/Å2.

#### Dynamic Light Scattering (DLS)

DLS size measurement was performed with a Zetasizer Pro-Red (Malvern Instrument, United Kingdom) in DTS0012 zetta cuvettes (Malvern Instrument, United Kingdom). 1 mL of liposome or IEX eluates (1 and 2) were added to the cuvette and measured by the instrument at 25°C with closed lid and 10 mW nominal laser power.

### Standard rEVs Preparation and Concentration Measurements

Standard recombinant EVs (Exosome standards, SAE0193-1VL) were purchased from Milipore Sigma with ∼1 × 10^9^ particles. Standard rEVs were reconstituted with 200 µL of PBS to reach ∼ 5 × 10^9^ particles/mL concentration of stock.

### Serial Dilution Experiments

Liposome and standard rEVs samples were serially diluted to ∼ 1 × 10^8^, ∼ 2 × 10^8^, ∼ 4 × 10^8^, ∼ 8 × 10^8^ particles/mL range and tested on scattering mode of NTA first. Then the samples were labeled by the optimal CMO concentration of 5 µg/mL and tested on the fluorescence mode of NTA. For Fl-FC with a wider dynamic range (1 × 10^6^ −5 × 10^8^particles/mL), and following the optimal labeling strategy, samples were diluted to ∼ 1 × 10^7^, ∼ 4 × 7, ∼ 2 × 10^8^, ∼ 5 × 10^8^ ∼ 1 × 10^9^, ∼ 4 × 10^9^ particles/mL and labeled with 5 µg/mL of CMO. Samples were incubated for 30 minutes and then diluted 10 times to the dynamic range of Fl-FC and measured.

### Analytical Chromatography

PATfix™ analytical platform (Sartorius BIA Separations) model LPG equipped with inline MALS detector, 10 mm flow path UV detector, and fluorescence detector (200-650 nm) was used in this study. The instrument was coupled with Agilent Bio SEC-3 column with 3 µm particle size, 300 Å pore size, 7.8 mm inner diameter and 300 mm length (Part number: 5190-2511, Agilent). For all the experiments in this study, 100 µL of samples were loaded into vials and 50 µL was injected at 0.5 mL/min flow rate and measurement time of 35 minutes. The column integrity was weekly assessed by running Agilent protein standard containing 5 different molecular weight proteins (Part number: 5190-9416, Agilent).

#### Crude Sample Model (BSA + Liposome) Study

Medium sized liposome samples were diluted to 1 × 10^11^ particles/mL and tested on analytical chromatography setup. The liposome samples ( 2 × 10^11^ particles/mL) were then mixed with BSA (1 mg/mL) at a 1:1 ratio. The samples were again tested on the SEC column at 0.5 mL/min flow rate to study separation of liposomes and BSA.

### Upstream Processing

#### Bioreactor Run

Human bone marrow derived MSCs were initially cultured on a 10-stack tissue culture dish (ThermoFisher) at 37°C with 5% CO_2_. The cells were plated on the 10-stack at 3000 cells/ *cm*^2^ in MSC NutriStem® XF basal medium supplemented with NutriStem® XF supplement and PLT (complete medium). Once confluent, they were trypsinized using TrypLE reagent (ThermoFisher) and added to the 10L single use Univessel ® bioreactor (Sartorius) containing 10 cm^2^/mL rhCol1 coated ACF microcarriers (10 *cm*^2^/*mL* x 10000 mL medium ÷ 360 *cm*^2^ /g 277.8 grams of microcarriers) in complete Nutristem ® XF medium at 3000 cells/cm^2^. Attachment of cells to microcarriers and cell growth were monitored daily. On day 4 the agitation was stopped and the microcarriers were allowed to settle at the bottom of the bioreactor after which 50% of the medium was removed and replaced with fresh complete medium. On day 6 the agitation was stopped and the microcarriers were allowed to settle at the bottom of the bioreactor after which the medium was removed and replaced with dPBS to both wash the MSC coated microcarriers and remove PLT. This was repeated for a total of two washes. The MSC coated microcarriers were then resuspended in 10 L of MSC NutriStem® XF basal medium supplemented with NutriStem® XF supplement (EV collection medium) and grown for an additional 2 days during which EVs were produced by the cells. Additional culture parameters are outlined supplementary Table 1. The harvest (MSCs CM) was then aliquoted into five 2L aliquots and were frozen at −80°C before transferring from Sartorius Ann Arbor site to Marlborough site. Upon arrival, The MSCs CM aliquots were thawed gradually from −80°C to −20°C (overnight), then 4°C until fully thawed, each 2L aliquot was then further aliquoted into eight 250 mL aliquots and were kept frozen at −80°C.

#### Cell attachment assay

Daily sampling to measure cell count and cell viability on microcarriers was performed by taking a 10 mL sample from the bioreactor each day. The sample was spun at 300 x g for 5 min, and the medium was removed. The pelleted material was washed with dPBS and spun at 300 x g for 5 min. The dPBS was removed and the pelleted material was trypsinized using 2 mL of TrypLE^TM^ express (ThermoFisher). After 5 min the reaction was quenched with 4 mL of dPBS and passed through a 70 µM cell strainer (Corning) to remove the microcarriers. The concentration and viability were then determined by loading the small amount of the sample onto a disposable Via1-Cassette™ and reading cassette in a NucleoCounter® NC-200™ cell counter (Chemometec).

### Downstream processing

#### Clarification

A two-step clarification strategy was used to remove residual microcarrier, dead cells, cell debris and larger particulates. MSCs conditioned medium containing EVs was pumped via a peristaltic pump (Masterflex) through a 5 µm Sartopure® PP3 (size 4, 0.018 m^2^) filter at a rate of 100 mL/min (∼334 LMH). The 5 µm filtered medium was then pumped through a 0.65 µm Sartopure® PP3 (size 4, 0.013 m^2^) filter at a flow rate of 100 mL/min (∼462 LMH). In scaled up experiments, all the operating parameters were kept the same, while for both 5 and 0.65 µm, the capsule size was increased to size 5 capsules with 0.036 m^2^ and 0.026 m^2^, respectively, leading to LMH values of 167 for 5 µm and 231 for 0.65 µm.

#### Tangential Flow Filtration 1

Prior to the TFF step, Benzonase (Millipore) was added at a concentration of 50 U/mL (0.4 µl of Benzonase per 100 mL of MSCs CM) to the clarified MSCs CM for 1 hour as the nuclease treatment step. Then the CM was processed using the Sartoflow Smart TFF system with a 300 KDa Sartocon® Slice 200 Eco PESU flat sheet cassette (3M81467901E—SW) with 0.02 m^2^ membrane area. The feed pressure was set at 1.75 bar, the permeate pressure was kept at 0.3 bar, while the retentive valve was kept open, leading to the TMP of 0.575 bar. The TFF step was conducted to perform 6X diafiltration to remove medium-associated components. The volume of the retentate was held constant by an auxiliary pump controlled by the machine that pumped dPBS into the system using the same conditions as described above.

In scaled up experiments, all the operating parameters were kept the same except 300 KDa Sartocon® Slice Eco Hydrosart was used with a 7 times larger surface area (0.14 m^2^ membrane area) proportional to the volume increase (100 mL to 700 mL MSCs CM). The TFF step was conducted to perform 6X diafiltration to remove medium-associated components. The volume of the retentate was held constant by an auxiliary pump controlled by the machine that pumped dPBS. In scaled up experiments, all the operating parameters were kept the same in Sartoflow Smart TFF system with 300 KDa Sartocon® Slice Eco Hydrosart was used with a 7 times larger surface area (0.14 m^2^ membrane area) proportional to the volume increase (100 mL to 700 mL CM).

#### Ion Exchange Chromatography (IEX)

Sartobind Convec® D nano (3 mL) membrane columns (Sartorius, Item No 97SC— 04E-C11--A) were used for IEX chromatography. The buffers composition as well as detailed information on IEX chromatography steps can be found in (**Sup Table [1-3]**). In summary, buffer A (20 mM Tris, 0 mM NaCl, pH 7.4), buffer B (20 mM Tris, 1000 mM NaCl, pH 7.4), wash buffer (20 mM Tris, 0 mM NaCl, pH 7.4) were prepared and filtered through 0.1 µm vacuum filters (Sartolab® RF 1000, Item No 180D05-E, Sartorius). The column was first flushed with 15 mL of buffer B at 15 mL/min, followed by an equilibration step with 60 mL of buffer A at 15 mL/min. The MSCs CM and medium alone control were both diluted 1:1.5 ratio to buffer A prior to loading to reduced conductivity to ∼7 mS/cm as recommended by the manual of Convec® D column. Total of 250 mL (100 mL of MSCs CM + 150 mL of buffer A) was loaded at 9 mL/min, followed by 60 mL of wash at 9 mL/min. For linear gradient strategy, the salt concentration was increased linearly from 0-1000 mM NaCl over 80 mL of elution at 3 mL/min. For stepwise elution strategy, first elution was done at 200 mM NaCl concentration for 10 mL at 3 mL/min flow rate. The first elution was then followed by a 20 mL wash step at the same salt concentration (20% of buffer B, 200 mM of NaCl) at 3 ml/min flow rate. Then, the second elution and the subsequent wash was done similarly as the first elution, 10 mL at 40% buffer B (400 mM NaCl) and 20 mL of wash at the same salt concentration at 3 ml/min flow rate. Then, the linear elution was conducted from 40-100% buffer B (400 mM – 1000 mM NaCl) for 10 mL at 3 mL/min flow rate, followed by a strip step at 100% buffer B (1000 mM NaCl) for another 10 mL at 3 mL/min flow rate.

In scalability study, Sartobind Convec® D Mini (20mL) membrane columns (Sartorius, Item No 97SC—04E4J11--A) were used for IEX chromatography. The buffers compositions were kept the same as before. The column was first flushed for 100 mL at 100 mL/min with Buffer B, followed by an equilibration step for 400 mL at 100 mL/min with buffer A. The MSCs CM and medium alone control were both diluted 1:1.5 ratio to buffer A prior to loading to reduced conductivity to ∼7 mS/cm as recommended by the manual of Convec® D column. A total of 1750 mL (700 mL of MSCs CM + 1050 mL of buffer A) was loaded at 60 mL/min, followed by 400 mL of wash at 60 mL/min. First elution was done at 20% of buffer B resulting in 200 mM NaCl concentration for 80 mL at 20 ml/min flow rate. The first elution was then followed by a 160 mL wash step at the same salt concentration (20% of buffer B, 200 mM of NaCl) at 20 ml/min flow rate. Then, the second elution and the subsequent wash were done similarly to the first elution, 80 mL at 40% buffer B (400 mM NaCl) and 160 mL of wash at the same salt concentration at 20 ml/min flow rate. Then, the linear elution was conducted from 40-100% buffer B (400 mM – 1000 mM NaCl) for 80 mL at 20 ml/min flow rate, followed by a strip step at 100% buffer B (1000 mM NaCl) for another 80 mL at 20 ml/min flow rate.

#### Tangential Flow Filtration 2

We included a 2nd TFF step in the scaled-up experiments. The eluate 2 (80 mL) was processed using the Sartoflow Smart TFF system with a 300 KDa Sartocon® Slice 200 Eco PESU flat sheet cassette (3M81467901E—SW) with 0.02 *m*^2^ membrane area. The feed pressure was set at 1.75 bar, the permeate pressure was kept at 0.3 bar, and the retentate valve was kept open, leading to the TMP of 0.575 bar. The TFF step was conducted to perform 6X diafiltration to reduce the salt concentration with buffer exchange from 400 mM salt to dPBS.

#### Sterile Filtration

The post TFF 2 retentate containing purified MSC derived EVs was filtered using a syringe pump at a constant flow rate of 5 mL/min using Sartoscale 25, Sartopore® 2 XLG 0.2 (Sartorius, Item No 5445307GV—LX—C). They were then aliquoted into 1mL aliquots and stored at −80 ^0°^C.

### Transmission Electron Microscopy

200 mesh formvar carbon coated copper grids were glow discharged 30s at 30 mA using room air. 3 µl of sample were applied to the carbon face of glow discharged grids for 30 seconds with the excess wicked away with Whatman paper. Grids were then incubated sequentially two times with molecular grade water and 0.75% uranyl formate by touching the carbon face of the grids to droplets of each solution and then immediately wicking the excess away. Grids were allowed to air dry prior to imaging. Samples were imaged using a Talos 120C transmission electron microscope equipped with a CETA 16 camera using Thermo Fisher’s TIA software. Electron microscopy was performed in the Electron Microscopy Resource (RRID: SCR_012366) in the Center for Advanced Research Technologies at the University of Rochester Medical Center.

### Capillary Electrophoresis-based Immunoassay (Simple Western)

Immunoassays were performed with the use of Jess™ Simple Western™ instrument (ProteinSimple, Cat# 004-650) according to manufacturer’s guidelines. Eluate 1 and 2 protein concentration was assessed using Pierce™ BCA Protein Assay Kit (Thermo Scientific, cat#23227). Next, eluate 1 and 2 were diluted in 0.1x sample buffer to the total protein concentration of 0.4 μg/μL. Then, the samples were prepared by mixing four parts of diluted eluate 1 and 2 were mixed with one part of 5x fluorescent master mix, containing 5x sample buffer, fluorescent standard, and 200 mM DTT from EZ standard pack (ProteinSimple, cat#PS-ST01EZ) and subsequent denaturation at 95 °C for 5 min. MSC and T-cell lysates were used as positive controls.

The following primary antibodies were diluted to the final concentration of 0.1 μg/μL (1:10) in antibody diluent #2 (ProteinSimple, cat#042-203) and used to probe for the proteins of interest: α-CD9 (Cell Signaling, cat#50–173-3388), α-CD81 (Proteintech, cat#MAB46152,), α-Flotillin-1 (SCBT, cat#sc-74566), α-Syntenin-1 (Abcam, cat#ab133267), α-Albumin (R&D systems, cat#MAB1455), α-CD73 (R&D systems, cat#MAB57951), α-CD105 (R&D systems, cat#AF1097) and α-GAPDH (R&D systems, cat#MAB1455).

Jess separation module with 12-230 kDa range (ProteinSimple, cat#SM-W001) was used to run the assay, according to manufacturer’s guidelines. Briefly, prepared samples, primary antibodies, secondary antibodies from anti-rabbit (ProteinSimple, cat#DM-001), anti-mouse (ProteinSimple, cat#DM-002), anti-goat (ProteinSimple, cat#DM-006) and anti-human (ProteinSimple, cat#DM-005) detection modules, biotinylated ladder, streptavidin-HRP (ProteinSimple, cat#042414) as well as Luminol-S (ProteinSimple, cat#043-311) and peroxide (ProteinSimple, cat#043-379) mixture were loaded on a microplate and chemiluminescence assay was set as follows: separation time – 30 min, separation voltage – 375 V, antibody diluent time – 30 min, primary antibody time – 30 min, secondary antibody time – 30 min. Chemiluminescent detection profile was set at High Dynamic Range 4.0 and contrast was manually adjusted for each sample. All data were analyzed using Compass for SW software, version 6.3.0 (ProteinSimple).

### Proteomic Analysis

#### Sample Preparation

Buffer composition was adjusted to 8M urea, pH 8.0, 1X Roche Complete Protease Inhibitor. Samples were sonicated using an amplitude of 35-40% with 1s pulses on and off for 20s. Lysates were clarified by centrifugation at 14,000 x g and supernatants transferred to a new tube. Protein quantitation was performed using Qubit. 50 μg of sample was digested overnight with trypsin. Briefly, samples were reduced for 1 hour at RT in 12 mM DTT followed by alkylation for 1 hour at RT at 15 mM iodoacetamide. Trypsin was added to an enzyme: substrate ratio of 1:20 for 18 h. Each sample was acidified to 0.3% TFA and subjected to SPE using Waters μHLB.

#### Mass Spectroscopy

500 ng per sample was analyzed on an LC/MS with a Thermo Fisher Vanquish Neo UPLC system interfaced to a Thermo Fisher Orbitrap Astral. Peptides were loaded on a trapping column and eluted over a 75 μm analytical column at 350 nL/min; the column was heated to 55° C. A 30 min gradient was employed. The mass spectrometer was operated in data-independent mode. Sequentially, full scan MS data (240,000 FWHM resolution) from m/z 380-980 was followed by 300 x 2 m/z precursor isolation windows; products were acquired in the Astral at 40,000 FWHM resolution. The maximum ion inject time (IIT) was set to 3.5 ms for data-independent acquisition (DIA); the NCE was set to 25.

### Lipidomic Analysis

#### Lipid extraction

Samples were thawed on ice and 100 µL of ultrapure water extract (containing protease inhibitors, phenylmethylsulfonyl fluoride, and EDTA) were added to resuspend the pellet. 50 µL of sample was combined with 500 µL of methanol (ThermoFisher), methyl tert-butyl ether (Fisher), and internal standard (Avanti/zzstandard). The mixture was vortexed for 15 mins. Then, 100 µL of water was added and the mixture vortexed for 1 min prior to centrifugation at 12,000 RPM and 4° C for 10 mins. 300 µL of supernatant was extracted and dried by vacuum concentration. Next, the powder was reconstituted with 200 µL of reconstitution solution and stored at −80° C. The remaining 50 µL of resuspended pellet was freeze-thawed 3 times, centrifuged at 12,000 RPM for 10 mins, and the supernatant taken to determine total protein concentration using the BCA Protein Assay Kit.

#### Chromatography-mass spectrometry

Ultra performance liquid chromatography (UPLC) (Nexera LC-40) and tandem mass spectrometry (MS/MS) (Triple Quad 6500+) were performed on the samples with a ThermoFisher Accucore C30 (2.6 μm, 2.1 mm x 100 mm) chromatographic column and the following mobile phases: A) acetonitrile (ThermoFisher)/water (60/40, V/V) (with 0.1% formic acid (Sigma Aldrich) and 10 mmol/L ammonium formate (Sigma Aldrich)); B) acetonitrile/isopropyl alcohol (Fisher) (10/90, V/V) (with 0.1% formic acid and 10 mmol/L ammonium formate). The gradient program was performed as follows: 80:20 (V/V) at 0 mins, 70:30 (V/V) at 2 mins, 40:60 (V/V) at 4 mins, 15:85 (V/V) at 9 mins, 10:90 (V/V) at 14 mins, 5:95 (V/V) at 15.5 mins, 5:95 (V/V) at 17.3 mins, 80:20 (V/V) at 17.5 mins, and 80:20 (V/V) at 20 mins. The flow rate was 0.35 mL/min, column temperature 45° C, and the injection volume 2 µL. The effluent was alternatively connected to an ESI-triple quadrupole-linear ion trap (QTRAP)-MS.

MS/MS was performed with the following conditions: LIT and triple quadrupole (QQQ) scans were acquired on a triple quadrupole-linear ion trap mass spectrometer (QTRAP) (QTRAP 6500+ LC-MS/MS System), equipped with an ESI Turbo Ion-Spray interface, operating in positive and negative ionization mode and controlled by Analyst 1.6.3 software (Sciex). The ESI source operation parameters were as follows: ion source, turbo spray; source temperature 500°C; ion spray voltage (IS) 5500 V (positive), −4500 V (negative); and ion source gas 1 (GS1), gas 2 (GS2), curtain gas (CUR) set at 45, 55, and 35 PSI, respectively. Instrument tuning and mass calibration were performed with 10 and 100 μmol/L polypropylene glycol solutions in QQQ and LIT modes, respectively. QQQ scans were acquired as MRM experiments with collision gas (nitrogen) set to 5 PSI. DP and CE for individual MRM transitions was done with further DP and CE optimization. A specific set of MRM transitions were monitored for each period according to the lipids eluted within the period.

### PicoGreen™ Assay for DNA Quantification

The quantification of the residual DNA was performed using Quant-iT PicoGreen dsDNA Assay Kits (ThermoFisher Scientific) following the manufacture’s protocol. Samples, Blank, and DNA standards (0.5 ng/mL – 1000 ng/mL) were prepared as needed. Subsequently, 100 μL of each blank, sample, and standard was added to a black 96-well plate, followed by the addition of 100 μL of working PicoGreen™ assay reagent. The plate was incubated in the dark at room temperature for 3 mins and then read at excitation of 480 nm and emission of 520 nm using a SpectroMax iD3 plate reader (Molecular Devices). Standards and samples absorbance were corrected by subtracting the absorbance of the blank samples. The optical density of the samples was interpolated from the standard curve using linear fitting.

### BCA Assay for Protein Concentration

The quantification of the residual protein was performed using Pierce™ BCA Protein Assay Kits (ThermoFisher Scientific, Waltham, MA) following the manufacture’s protocol. Samples, Blank, and bovine serum albumin (BSA) standards (25 μg/mL – 2000 μg/mL) were prepared as needed and 25 μL of each blank, samples and standards were added to a clear 96 well plate. 200 μL of BCA working reagent was then added and mixed thoroughly on a plate shaker for 30 seconds. The plate was then incubated at 37°C for 30 mins and the absorbance was measured at 562 nm using a SpectroMax iD3 plate reader (Molecular Devices, San Jose, CA). Standards and samples absorbance were corrected by subtracting the absorbance of the blank samples. The optical density of the samples was interpolated from the standard curve using 4PL curve fitting.

### Scratch Wound Healing Assay

Adult human dermal fibroblasts (hDFA, ATCC) were expanded according to the supplier’s instructions. Briefly, cells were seeded at 3333 cells/cm^2^ in a T150 flask (Corning) in hDFA growth medium (hDFA-GM) consisting of DMEM (Gibco) with 10% FBS (VWR) and 1% L-glutamine (Gibco) and penicillin/streptomycin (Gibco). Medium was exchanged 24 hours after seeding and cells were expanded for a total of 5 days (80% confluence) before passaging and banking. For the scratch assay, P6 hDFA were thawed and expanded as described for 3 days (80% confluence) before seeding 15000 cells/well in an Imagelock 96 well plate (Sartorius) coated with 100 µg/mL rat tail collagen (Corning). Cells were allowed to adhere overnight. 6 hours prior to scratching, the medium was exchanged with 0.1% FBS hDFA-GM. Cells were scratched using the Woundmaker tool (Sartorius) and washed with DMEM. The +CTL was complete hDFA-GM (10% FBS), the -CTL was 0% FBS hDFA-GM, and the treatment was 50 µL of EV preparation (8.46E4 particles/seeded cell) + 200 µL of -CTL medium. The vehicle control used 50 µL of PBS (Sartorius) instead of EV preparation.

Live imaging was performed using the Incucyte (Sartorius) with imaging at 10X in phase and orange channels every 2 hours for 48 hours. Images were analyzed using a custom Python script that identifies and tracks nuclei in the phase image and counts the number of cells that migrate into the scratch area. Scratch closure is quantified using the number of cells that migrate into the scratch normalized to the number of cells in the unscratched area of the same field of view in each image.

### Additional Materials and Methods

Additional methods and materials can be found in supplementary material. All statistical analyses were done using GraphPad prism v10.

## Results

### Analytical Optimization

#### Optimization of NTA parameters

Single particle analysis techniques are used widely to measure the concentration in EV samples and further to assess and optimize EV production and purification processes (Arab *et al*., 2021; Dong *et al*., 2020). Among these techniques, NTA and FC are notable due to their ability to analyze EV samples in both scattering and fluorescence mode *(Dlugolecka et al., 2021; Kobayashi et al., 2024; Mladenović et al., 2025;* *Tirelli et al*., 2025). NTA is a technique to measure size and concentration of EVs in both scattering and fluorescence mode by tracking their Brownian motion. FC relies on hydrodynamic focusing principle and has been widely used in the EV field to measure size and concentration. The FC used in this study, Virus Counter® 3100, is a fluorescence-based technique that has been shown to quantify EVs (Dehghani *et al*., 2021; Kruse *et al*., 2022). In addition to the quantification, we also further investigated the characteristics of individual peaks detected. This enabled detailed sample analysis to ensure single event detection and avoid coincidence or swarm-based detection **(Sup Fig 1)**.

However, recent studies have shown inconsistencies and lack of reproducibility between different analysis modes and techniques (Arab *et al*., 2021; Nguyen *et al*., 2024; Vestad *et al*., 2017). The most recent “Minimal Information for Studies of Extracellular Vesicles (MISEV)” guidelines further highlighted the extracellular particles family containing both vesicular and non-vesicular particles which can complicate accurate single EV analysis *(Welsh et al*., 2024). Additionally, the lack of standard reference materials has been emphasized as a major obstacle for reliable and reproducible EV concentration measurements using single particle analysis techniques (Welsh *et al*., 2023, 2020b).

In this study, we selected liposomes as reference materials to optimize the analytical assays for EV quantification using NTA and Fl-FC since; A) liposomes are vesicular and do not contain non-vesicular particles; B) they can be synthesized with a similar size distribution as EVs; C) they exhibit a refractive index similar to EVs which is an important consideration in scattering-based detections; D) their lipid bilayer structure makes them suitable for fluorescence-based detection using lipophilic dyes. Given these characteristics, liposomes can be considered as artificial vesicles for analytical assay development. We hypothesized that, under optimal assay parameters, liposome concentrations measured by scattering and fluorescence mode of NTA, as well as by Fl-FC, would result in comparable values.

The CellMask™ dye family have been used successfully to label and quantify EVs with NTA and FC (Brealey *et al*., 2024b; Midekessa *et al*., 2021; Nikiforova *et al*., 2021). We previously demonstrated that CellMask™ Orange (CMO) dye, a lipophilic dye that integrates into lipid bilayers, enables reliable labeling and quantification of EVs from various sources using FC (Dehghani *et al*., 2021). Building on this approach, we applied the same labeling strategy to quantify liposomes using the fluorescence mode of NTA, with a 60 second acquisition time in static mode.

Initial results revealed significant discrepancies in particle concentrations measured by NTA in scattering versus fluorescence mode (**Sup Fig 2-a**). Further inspection of the Fl-NTA videos indicated a pronounced photobleaching effect, as evidenced by a visible decline in fluorescence intensity over time (**Sup Fig 2-b**), comparing the first and last video frames. To mitigate photobleaching, we reduced the acquisition times to 30 and 15 seconds; however, concentration measurements from the fluorescence mode continued to underestimate the values relative to the scattering mode, suggesting that photobleaching remained a confounding factor. To address the observed photobleaching issue, we transitioned from static mode to flow mode, wherein particles are driven through the laser beam using a syringe pump at a controlled flow rate. We evaluated three flow rate settings (unit values: 20, 35, and 50). Although the absolute flow rate values for these units are not specified, higher unit numbers correspond to faster flow rates. At unit 20, discrepancies between fluorescence and scattering measurements persisted, likely due to prolonged laser exposure and ongoing photobleaching. In contrast, the flow rate unit 50 resulted in abnormally high concentration values across both detection modes (**Sup Fig 2-a and c**). After reviewing the recorded videos, we concluded that this flow rate exceeded the tracking capabilities of the software, resulting in noisy and unreliable measurements in both scattering and fluorescence mode.

Among the tested settings, flow rate unit 35 provided the most consistent and accurate results. At this setting, measured concentrations of liposomes in fluorescence and scattering modes were not significantly different, indicating minimal photobleaching and stable particle detection. Consequently, we selected flow mode at unit 35 as the optimal operating condition for all subsequent NTA fluorescence analyses in this study.

#### Optimization of Fluorescence Labeling

Fluorescence-based enumeration of single EVs using fluorescence-based flow cytometry (Fl-FC) and nanoparticle tracking analysis (Fl-NTA) typically requires a washing step to remove unbound dye after labeling. Commonly used methods for this purpose include size exclusion chromatography, ultrafiltration, and ultracentrifugation. While effective, these washing procedures can result in EV loss and can make optimization of downstream processing more challenging (Dehghani and Gaborski, 2020; Fortunato *et al*., 2021; Rautaniemi *et al*., 2022).

To overcome these limitations, we aimed to develop an optimized labeling protocol that eliminates the need for a washing step. To achieve this, we systematically investigated the influence of the dye-to-particle ratio on concentration measurements using orthogonal techniques (Andronico *et al*., 2021). Our objective was to identify a dye concentration that provides a high fluorescence signal in the presence of liposomes, while maintaining minimal background signal in dye alone controls (i.e., in the absence of liposomes), thereby maximizing the signal-to-noise ratio.

To determine the optimal dye concentration for labeling without a washing step, we first monitored fluorescence intensity using a plate reader while maintaining a constant liposome concentration while systematically increasing the CMO concentration from 0.39 µg/mL to 25 µg/mL (**Fig 1-a**). Within the CMO concentration range of 0.76 µg/mL to 6.25 µg/mL, labeled liposome samples exhibited significantly higher fluorescence signals compared to dye alone controls, indicating effective labeling with minimal background. However, at concentrations above 6.25 µg/mL, the fluorescence intensity of the dye alone controls exceeded that of the labeled liposome samples, suggesting an increase in background signal due to excess unbound dye and reduced signal-to-noise ratio. These findings indicate that CMO concentrations of 12 µg/mL and 25 µg/mL are too high for effective wash-free labeling. Furthermore, CMO-labeled liposome samples displayed a stable fluorescence signal for up to 2 hours, with peak emission centered around 575 nm, consistent with the excitation/emission detection settings of both Fl-NTA and Fl-FC (**Fig 1-a**).

**Figure 1.**
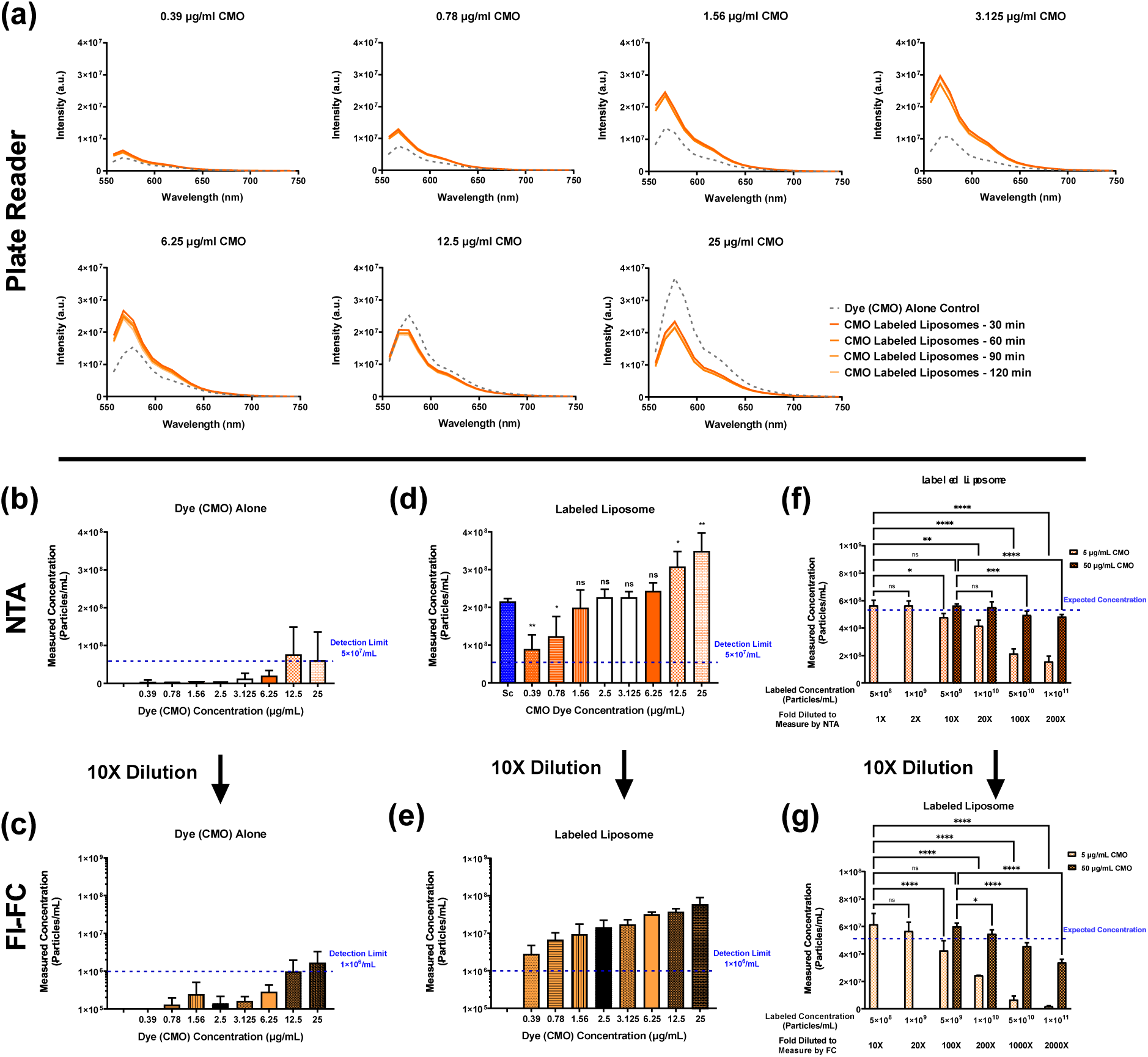
**Analytical Optimization;** Establishing a no-wash assay for fluorescence labeling of liposomes using CellMask™ Orange (CMO) dye. **(a)** Fluorescence emission spectra of CMO labeled liposomes and dye alone control over time (30 to 120 min) by systematically increasing dye concentrations (0.39 to 25 µg/mL). **(b, c)** Particle concentrations detected in dye only controls (no liposomes) in **(b)** Fl-NTA and **(c)** Fl-FC. **(d, e)** CMO labeled liposomes concentrations measured by **(d)** Fl-NTA and **(e)** Fl-FC. **(f, g)** Evaluation of dynamic range and labeling efficiency while systematically increasing liposome concentrations labeled with CMO measured by **(f)** Fl-NTA and **(g)** Fl-FC. Using 5 µg/mL CMO, reliable detection was maintained up to 1 × 10^9^particles/mL; a higher dye concentration (50 µg/mL) was required for accurate quantification at liposome concentrations from 1 × 10^9^till 1 × 10^10^particles/mL. Multicomparison statistical analysis was performed using ordinary one-way ANOVA **(c)** or two-way ANOVA **(f and g)** with Tukey’s multiple comparison test (*p<0.05, **p<0.01, ***p<0.001, ****p < 0.0001; ns not significant). All data are presented as mean ± standard deviation with n 3 independent measurements. Sc: Scattering mode, Fl: Fluorescence mode of NTA, Fl-FC: Fluorescence-based flow cytometry. CMO: CellMask™ Orange dye.

As recommended by the MIFlowCyt-EV guidelines (a framework for standardized reporting of extracellular vesicles flow cytometry experiments), dye alone control is essential in single particle analysis to distinguish true EV signals from dye-associated artifacts (Welsh *et al*., 2020a). Recent studies have reported that certain lipophilic dyes, such as PKH dyes, can form micelles or aggregates that are indistinguishable from labeled EVs in fluorescence-based assays (Dehghani *et al*., 2020; Pužar Dominkuš *et al*., 2018; Takov *et al*., 2017). Therefore, it is critical to confirm that the number of detected events in dye alone controls remain below the assay’s lower LOD. To address this, we extended our initial plate reader fluorescence measurements by systematically increasing the CMO concentration and evaluating the labeling efficiency and background signal using both Fl-NTA and Fl-FC.

While the dynamic range for NTA is reported to be approximately 5 × 10^7^ to 1 × 10^9^ particles/mL, the dynamic range for Fl-FC is wider, spanning from 1 × 10^6^ to 1 × 10^8^ particles/mL. We first evaluated the CMO dye alone control using Fl-NTA and then subsequently diluted 10-fold for Fl-FC based on the dynamic range differences between two instruments. In the CMO concentration range lower than 6.25 µg/mL, the number of detected events remained below the lower LOD for both Fl-NTA ( 5 × 10^7^ particles/mL) and Fl-FC ( 1 × 10^6^particles/mL), indicating minimal background interference (**Fig 1-b and c**). However, at higher dye concentrations (12.5 and 25 µg/mL), both techniques detected particle concentrations above their respective lower LOD, suggesting the presence of dye-associated artifacts that could lead to false-positive particle counts in labeled samples (**Fig 1-b and c**). Further analysis of the Fl-NTA videos revealed pronounced background fluorescence signal and visual cloudiness at higher concentrations (**Sup Fig 3-a**). Similarly, baseline fluorescence analysis in Fl-FC showed elevated values for 12.5 and 25 µg/mL dye alone controls compared to the buffer control, further indicating increased background noise (**Sup Fig 3-b**).

Next, we labeled liposome samples using the same CMO concentration range tested in the dye alone control experiments and analyzed them using both Fl-NTA and Fl-FC. At the lowest CMO concentrations tested (0.39 and 0.78 µg/mL), the measured liposome concentrations were below those obtained via the scattering mode of NTA, likely due to insufficient labeling (**Fig 1-d**). As the CMO concentration increased from 1.56 to 6.25 µg/mL, the measured concentration increased and eventually plateaued, aligning closely with the scattering mode NTA values—indicating effective labeling with minimal background interference (**Fig 1-d**). As expected, CMO concentrations above 6.25 µg/mL resulted in apparent liposome concentrations that exceeded those measured in scattering mode, consistent with the elevated background signal previously observed in dye alone controls (**Sup Fig 3-a**). Again, labeled samples from Fl-NTA measurements were then diluted 10-fold to measure the concentration on Fl-FC. A similar trend as Fl-NTA was observed for Fl-FC showing insufficient labeling at 0.39 and 0.78 µg/mL CMO, plateau region in particle concentrations up to 6.25 µg/mL of CMO and then an increase in the particle concentration for 12.5 and 25 µg/mL of CMO (**Fig 1-e**). These findings were further supported by baseline fluorescence values, which were noticeably elevated in the high CMO concentration groups (**Sup Fig 3-[b and c]**). Additional analyses of Fl-FC data, including one-second fluorescence histograms and two-dimensional peak plots, confirmed the presence of background noise at higher dye concentrations and validated the trends observed across both dye alone and labeled samples (**Sup Fig 3-[d and e]**).

Based on the results from plate reader fluorescence measurements, Fl-NTA, and Fl-FC analyses, we identified 5 µg/mL as the optimal CMO concentration for EV labeling. This concentration provided high labeling efficiency while maintaining event counts in dye alone controls below the lower LOD, thereby eliminating the need for a dye washing step. This approach enabled us to define an optimal labeling protocol that maximizes fluorescence signal from labeled particles while minimizing false-positive events in the dye only control.

To determine the upper limit of liposome concentration that can be effectively labeled using 5 µg/mL CMO without washing, we systematically increased the liposome concentration. For liposome concentrations exceeding the upper detection limit of either instrument, samples were appropriately diluted to fall within the respective dynamic ranges of Fl-NTA and Fl-FC. Our results showed that 1 × 10^9^particles/mL is the highest liposome concentration that can be reliably labeled with 5 µg/mL CMO, as determined by both Fl-NTA and Fl-FC. Notably, at liposome concentrations above this threshold, a sharper decline in detected particle counts was observed in Fl-FC compared to Fl-NTA, suggesting that Fl-FC is more sensitive to suboptimal labeling and requires control over dye-to-particle ratios for accurate quantification (**Fig 1-[f and g]**). Then, to evaluate whether higher dye concentrations could extend the usable range, we increased the CMO concentration 10-fold to 50 µg/mL and labeled liposome samples at concentrations above 1 × 10^9^ particles/mL. As expected, this adjustment enabled efficient labeling and quantification of liposome concentrations up to 1 × 10^10^particles/mL using both Fl-NTA and FC (**Fig 1-[f and g]**). Similar decay was observed for concentrations above 1 × 10^10^particles/mL for both Fl-NTA and Fl-FC, suggesting higher dye concentration is required to properly label liposomes at concentrations above 1 × 10^10^ particles/mL.

Under these optimized labeling conditions, both techniques yielded consistent concentration measurements, indicating robust and accurate enumeration. Taken together, we established a no-wash fluorescence labeling protocol optimized for both CMO and liposome concentrations, as summarized in (**Sup Fig 4-a**). Unless otherwise stated, all subsequent fluorescence-based measurements were performed in accordance with the schematic outlined in (**Sup Fig 4-a**).

Next, we evaluated the size distribution of three different size liposome samples using orthogonal analytical techniques and compared the results with cryo-preservation electron microscopy (cryo-EM) as the reference standard. To obtain statistically robust cryo-EM measurements, over 1,000 individual particles were analyzed per sample, and particle size distributions were plotted as frequency versus diameter (**Sup Fig 5-[a-c]).** We then compared these cryo-EM distributions with those obtained from scattering mode NTA in both static and flow modes (Sc-NTA), fluorescence mode NTA in flow mode (Fl-NTA), and dynamic light scattering (DLS). Among these techniques, Fl-NTA in flow mode produced size distribution profiles most closely aligned with the cryo-EM data, accurately capturing the overall size distribution width and the frequency (**Sup Fig 5-[d-f]**). In contrast, the scattering-based techniques (Sc-NTA and DLS) consistently exhibited a shift toward larger particle sizes, likely due to their higher sensitivity to larger particles with higher scattering intensity within polydisperse samples. These findings highlight the advantages of fluorescence-based single particle tracking, particularly when used under optimal labeling conditions, for more accurate size profiling (**Sup Fig 5-[d-f]**).

#### Assessment of Concentration Measurement Accuracy

Following the optimization of both operating parameters and the no-wash labeling protocol, we assessed the accuracy and linearity of concentration measurements across the dynamic range of each instrument. In accordance with the MIFlowCyt guidelines (Welsh *et al*., 2020a), which recommend verifying single particle detection capability, we conducted serial dilution experiments using liposome samples of varying sizes (**Fig 2-a**). The measurements began at the lowest concentration above the detection threshold of the buffer control (<5 × 10^7^particles/mL for NTA and <1 × 10^6^particles/mL for Fl-FC) and proceeded with approximately 2-fold incremental increases up to the respective upper detection limits of SC-NTA and Fl-NTA. The *expected concentration* for each dilution was calculated theoretically based on the first measurable data point above the buffer control and was assumed to increase proportionally with the fold-change in particle number (represented by black dashed lines in **Fig 2-[b and c]**). This approach allowed us to evaluate the dynamic range and quantitative accuracy of each technique under optimized assay conditions.

**Figure 2.**
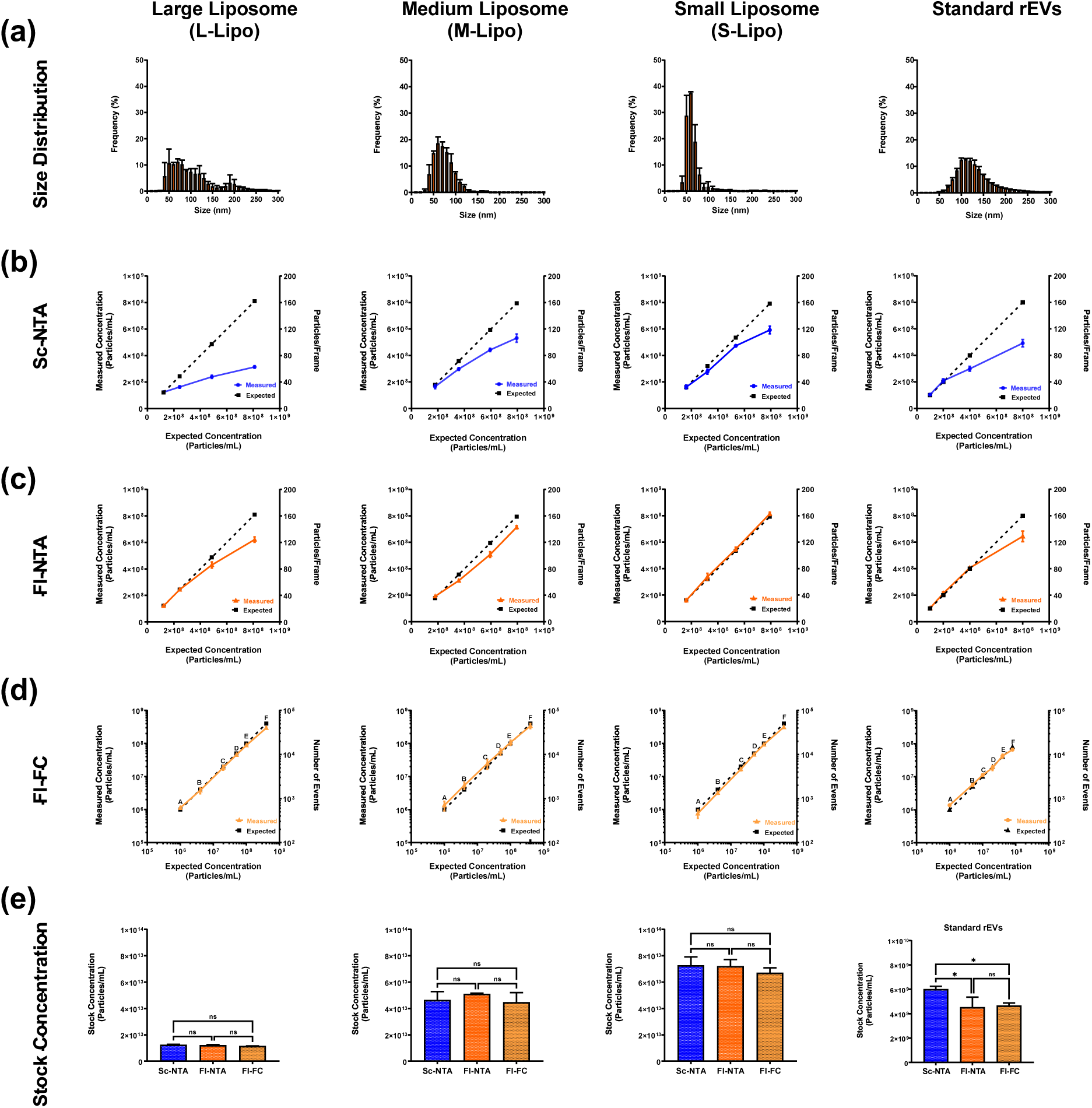
**Analytical Optimization;** Evaluating concentration measurements accuracy across three single particle analysis techniques. **(a)** Size distribution of large (L-Lipo), medium (M-Lipo), and small (S-Lipo) liposomes, as well as standard recombinant extracellular vesicles (rEVs), measured by Fl-NTA. **(b–d)** Measured (solid lines) versus expected concentration (black dash lines) plots across dilution series using **(b)** Sc-NTA, **(c)** Fl-NTA, and **(d)** Fl-FC. **(e)** Stock concentration measurements across all samples under optimal labeling and measurement conditions leading to similar particle concentrations across all three single particle analysis technologies. Multicomparison statistical analysis was performed using ordinary one-way ANOVA with Tukey’s multiple comparison test (*p<0.05, **p<0.01, ***p<0.001, ****p < 0.0001; ns not significant). All data are presented as mean ± standard deviation with 3 independent measurements. Sc-NTA: Scattering mode of NTA, Fl-NTA: Fluorescence mode of NTA, Fl-FC: Fluorescence-based flow cytometry.

We initially evaluated the concentration accuracy of unlabeled liposome samples of varying sizes using Sc-NTA. For large liposome samples characterized by a broad and heterogeneous size distribution (L-Lipo), the measured concentrations deviated from the expected values and did not exhibit linearity across the serial dilutions (**Fig 2-b**). This deviation became more pronounced at higher concentrations and is attributed to the *masking effect*, a known limitation of Sc-NTA in polydisperse samples (Bachurski *et al*., 2019; Dehghani *et al*., 2021). The masking effect arises from the strong size dependency of the scattering signal, which scales with the sixth power of the particle radius. As a result, larger particles dominate the scattering signal and obscure the detection of smaller particles, leading to underestimation of total particle concentration in heterogeneous samples. As expected, the medium-sized (M-Lipo) and small-sized (S-Lipo) liposome samples which were less heterogenous in size, demonstrated improved linearity in the dilution series compared to L-Lipo. Nevertheless, even for M-Lipo and S-Lipo, deviations from linearity were observed at the upper end of the dynamic range, indicating reduced accuracy in concentration measurements under large particle numbers (**Fig 2-b**).

We hypothesized that Fl-NTA would be less susceptible to the masking effect compared to Sc-NTA, given that the fluorescence intensity of lipophilic dye labeled particles scales with the square of the particle radius (surface area), as opposed to the sixth power in light scattering. To test this hypothesis, we conducted the same serial dilution experiments using CMO-labeled liposome samples and assessed the linearity of concentration measurements in Fl-NTA (**Fig 2-c**). As anticipated, Fl-NTA showed improved linearity across the dilution series compared to Sc-NTA, particularly for medium-sized (M-Lipo) and small-sized (S-Lipo) liposomes (**Fig 2-c**). However, for the large and polydisperse liposome samples (L-Lipo), concentration measurements at the higher concentration still deviated from expected values, indicating that the masking effect persists, albeit to a lesser extent, in fluorescence-based detection (**Fig 2-c**).

To determine whether this masking phenomenon is specific to liposomes, we repeated the linearity experiments using standardized recombinant extracellular vesicles (rEVs) (Geeurickx *et al*., 2019) (**Fig 2-[a-c]**). Similar trends were observed: both Sc-NTA and Fl-NTA showed deviations at higher concentrations, with the masking effect more pronounced in Sc-NTA. This finding supports the interpretation that the observed discrepancies are related to signal intensity scaling and are not unique to liposome composition. Based on these observations, we conclude that NTA-based quantification is most accurate within the lower half of the instrument’s dynamic range and is influenced by particle size heterogeneity. We found that Sc-NTA measurements are most reliable when fewer than ∼30 particles are present per frame, while Fl-NTA remains accurate up to approximately ∼90 particles per frame, offering a broader linear detection window under optimal labeling conditions (**Sup Figure 4-b**).

To confirm that the observed masking effect was not operator-dependent, we repeated the serial dilution experiments using M-Lipo samples with two additional operators on Sc-NTA. The results consistently showed deviations between the measured and expected concentrations, mirroring our initial findings suggesting that the inaccuracy is not influenced by the operator (**Sup Fig 6-a**). To further verify that the masking effect was not instrument-specific, the same linearity experiments were conducted on two additional NTA instruments, located in separate laboratories. In all cases, the masking phenomenon was reproducible, demonstrating measurement deviations across systems (**Sup Fig 6-b**). These findings collectively indicate that the masking effect is an intrinsic limitation of NTA technology under conditions of high particle concentration and/or heterogeneity, rather than a consequence of operator variability or instrument-specific artifacts.

We next evaluated the performance of Fl-FC for concentration measurements using the same set of liposome samples. Unlike NTA, Fl-FC measurements were not affected by the masking effect, owing to the flow-based design of the instrument, which ensures that particles pass through the interrogation point individually (**Fig 2-d**). Across all liposome sizes, the measured concentrations followed the expected linear trend in the serial dilution experiments. However, at concentrations approaching the upper detection limit, we observed slight deviations caused by coincidence events—instances where two or more particles pass through the laser simultaneously and are mistakenly recorded as a single event (Dehghani *et al*., 2021). The number of events detected in controls such as dye alone and buffer alone control were below lower LOD in Fl-FC (**Sup Fig 7-a**). Detailed analysis of the peak characteristics revealed the emergence of a distinct population at high concentrations, characterized by increased peak widths in 2D plots and higher baseline values (**Sup Fig 7-[b-d]**). This profile is indicative of a broadened Gaussian curve resulting from simultaneous detection of adjacent particles. Correspondingly, we also observed an elevation in baseline fluorescence values at these concentrations, further supporting the presence of coincidence artifacts. These findings demonstrate that while Fl-FC is not subject to the masking effect observed in NTA, accuracy at high particle concentrations may still be compromised by coincidence. However, these artifacts can be identified and mitigated through peak characteristics and baseline signal analysis. Similar results were obtained with recombinant extracellular vesicles (rEVs), that also exhibited linear concentration measurements across the dilution series, further validating the robustness of Fl-FC for quantitative single particle analysis in this context (**Sup Fig 7-e**).

Using optimized NTA operating parameters, including controlled particle number per frame and the validated no-wash fluorescence labeling protocol, we obtained consistent and reproducible stock concentration measurements for liposomes of varying sizes (**Fig 2-e**). In the case of recombinant extracellular vesicles (rEVs) across three orthogonal analytical platforms, we detected lower concentrations on Fl-NTA and Fl-FC compared to Sc-NTA, possibly due to presence of impurities like non-vesicular particles which contributes to the higher scattering values (**Fig 2-e**).

To further assess the reproducibility of our optimized assays, we measured the stock concentrations of rEV samples across three independent Fl-FC and NTA instruments. The results demonstrated high consistency in concentration measurements, confirming the robustness and inter-instrument reproducibility of the optimized protocols (**Sup Fig 6-c**). These findings collectively support the reliability of established analytical toolbox for accurate and reproducible quantification of single particles using both reference materials (liposomes) and recombinant EV (rEVs) preparations.

Single particle analysis techniques are known to be less reliable in crude or unpurified samples due to the presence of impurities that can interfere with particle detection. To assess the impact of such impurities on measurement accuracy, we evaluated liposome quantification in the presence of BSA as a model contaminant. We first performed a systematic BSA dilution series and analyzed samples using both Sc-NTA and Fl-NTA. For Sc-NTA, BSA concentrations ranging from 1 to 50 µg/mL remained below the instrument’s detection limit (**Sup Fig 8-[a and b]**). However, at 100 µg/mL, BSA alone produced detectable particle counts above the threshold, accompanied by visual cloudiness and increased background signal (**Sup Fig 8-b**).

In contrast, Fl-NTA detected events at BSA concentrations as low as 10 µg/mL, likely due to nonspecific interactions between the CMO dye and BSA (**Sup Fig 8-[a and c]**). Similar results were observed with Fl-FC, where concentrations of 10 µg/mL BSA and larger (followed by 10 times dilution) produced counts above the lower LOD and elevated baseline values (**Sup Fig 8-[d and e]**). Based on these findings, we established maximum allowable impurity concentrations of 50 µg/mL for Sc-NTA and 10 µg/mL for both Fl-NTA and Fl-FC to ensure reliable particle enumeration in crude samples. To validate the robustness of liposome quantification in the presence of BSA, we compared measurements of liposomes alone, BSA alone, and liposome-BSA mixtures (1:1 ratio). When BSA concentrations were maintained below the defined thresholds, liposome concentrations remained consistent across all three single particle analysis platforms (**Sup Fig 8-f**). Furthermore, BSA did not alter the size distribution profiles of liposomes in either Sc-NTA or Fl-NTA (**Sup Fig 8-[g-i]**), nor did it affect signal characteristics in Fl-FC, as evidenced by stable 2D peak plots and one-second histogram profiles (**Sup Fig 8-[m-r]**).

#### Analytical Chromatography Evaluation

To further expand our analytical toolbox, we integrated analytical chromatography into our workflow. The setup consisted of a size exclusion chromatography (SEC) column for particle separation, followed by in-line UV, MALS, and fluorescence detectors (**Fig 3-a**). SEC operates on the principle of size-based separation: particles larger than the pore size of the column matrix (>300 Å for the column used in this study) are excluded from the resin and elute first, forming the so-called void peak. Smaller molecules are retained longer within the column pores and elute later, enabling separation by size. In addition to in-line MALS and UV, we leveraged the intrinsic fluorescence properties of proteins— specifically their aromatic amino acids like tryptophan—by setting the excitation wavelength at 280 nm and the emission at 350 nm, that allows for sensitive detection of protein-based impurities (Nishimura *et al*., 2024; Normak *et al*., 2023b).

**Figure 3.**
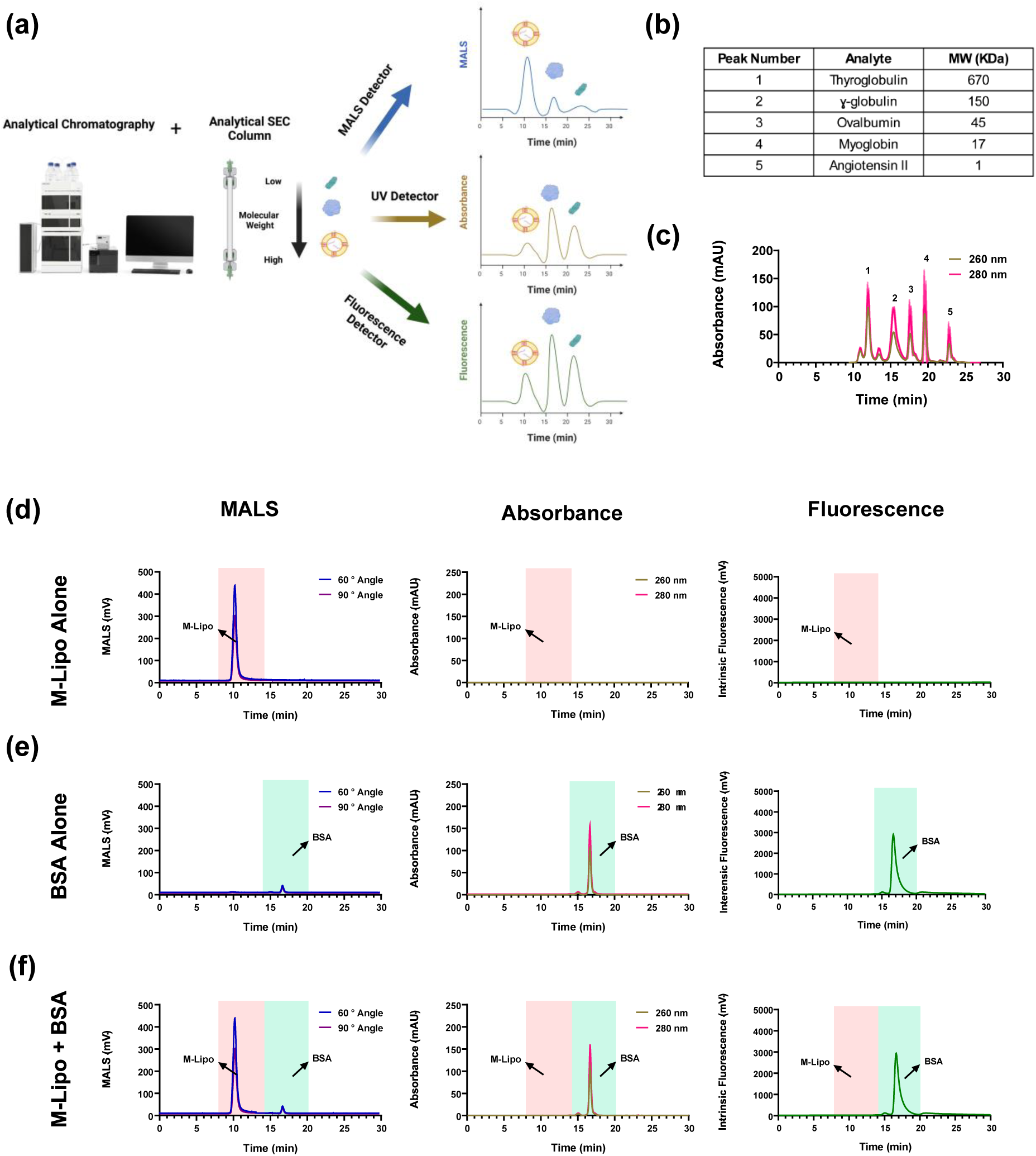
**Analytical Optimization;** Establishing analytical assays using multi-detector analytical chromatography. **(a)** Schematic of the analytical chromatography setup integrating size exclusion chromatography (SEC) with multi-angle light scattering (MALS), ultraviolet (UV), and intrinsic fluorescence detectors for comprehensive sample characterization. **(b)** Molecular weight (MW) table for the protein standard mixture used to validate size-based separation. **(c)** UV absorbance chromatogram of the protein standard mix at 260 nm and 280 nm, confirming effective resolution of proteins with varying MW. **(d–f)** Chromatograms of BSA alone **(d)**, M-Lipo alone **(e)**, and a mixture of M-Lipo + BSA **(f)** measured by UV absorbance, intrinsic fluorescence (Ex 280 nm / Em 350 nm), and MALS (60° and 90° angles). Liposomes eluted within the void volume (9–14 min, highlighted in red), while BSA eluted later (14–20 min, highlighted in green). Schematics were created in Biorender.com.

This multi-detector configuration enables comprehensive profiling of both crude and purified samples, facilitating the distinction between target particles (e.g., liposomes and EVs) and contaminating biomolecules (e.g., free proteins, RNA, and DNA). In principle, liposomes and EVs are expected to exhibit strong MALS signals due to their larger size, while showing minimal UV absorbance and intrinsic fluorescence (**Fig 3-a**). In contrast, free biomolecules such as proteins and nucleic acids typically yield high UV absorbance and fluorescence signals, but low MALS intensity—unless present at very high concentrations. Together, this orthogonal detection approach enables detailed compositional analysis and assessment of sample purity. To establish the elution profiles of molecules of varying sizes, we first analyzed a protein standard mixture (Agilent) comprising five proteins of known molecular weight (**Fig 3-b**). The largest protein, thyroglobulin (670 kDa), eluted between 11–14 minutes, corresponding to the void volume, while the smaller proteins—within the pore size of the SEC resin—eluted later and were well resolved, confirming effective size-based separation (**Fig 3-c**).

Then, like single particle analysis techniques, we first utilized liposomes as artificial vesicles (EV reference material) to better understand the performance of analytical chromatography. Liposomes produced a strong signal on the in-line MALS detector, with negligible UV absorbance and intrinsic fluorescence signal (minutes 9-14, highlighted as red color). To further study the detection range of analytical chromatography setup, we performed a serial dilution of medium-sized liposomes (M-Lipo), testing concentrations from 1 × 10^9^ to 5 × 10^11^ particles/mL. Due to their large size, liposomes eluted within the void volume (9–14 minutes; highlighted in red (**Fig 3-d and Sup Fig 9-a**). The in-line MALS detector used in this study features eight scattering angles, with 60° and 90° yielding the highest signal intensity. From the dilution series, we determined that the lower detection limit for M-Lipo was approximately 1 × 10^9^particles/mL. Furthermore, when comparing liposome samples of different sizes at the same particle concentration, the MALS signal followed a size dependent trend, with large liposomes (L-Lipo) producing the highest intensity, followed by M-Lipo and small liposomes (S-Lipo), respectively, which are consistent with light scattering principles (**Sup Fig 9-e**).

To evaluate the dynamic range of impurity detection in our analytical chromatography setup, we conducted a serial dilution experiment using BSA as a model impurity. BSA eluted between 14–20 minutes, consistent with its smaller hydrodynamic size relative to the SEC column pore size (highlighted in green, **Fig 3-e and Sup Fig 9-b**). A strong linear correlation was observed between BSA concentration and absorbance at both 260 nm and 280 nm, with a lower LOD of 20 µg/mL. Intrinsic fluorescence detection—using excitation at 280 nm and emission at 350 nm—demonstrated even greater sensitivity, with a lower LOD of 0.4 µg/mL (highlighted in green, **Fig 3-e and Sup Fig 9-c**). Calibration curves were generated for both absorbance and fluorescence modes by plotting peak intensity and area under the curve (AUC) against BSA concentration, confirming the semi-quantitative capability of the system for protein-based impurities (**Sup Fig 9-d**). When BSA and liposomes were mixed and analyzed using the analytical chromatography setup, we observed complete separation between the two components, as evidenced by distinct elution profiles (**Fig 3-f**). These data demonstrate that the SEC– MALS–UV–fluorescence setup enables orthogonal profiling of EV samples and co-isolated impurities with high sensitivity, resolution, and quantification accuracy, supporting its utility for EV process monitoring and purity assessment.

### Upstream Processing

We aimed to develop a scalable manufacturing process to produce EVs EVs from human bone marrow-derived mesenchymal stromal cells (hBM-MSCs). Cells were cultured in a bioreactor with microcarriers. hBM-MSCs were cultured for 6 days in MSC basal medium supplemented with supplement and human platelet lysate (PLT) **(Fig 4-a)**. Cell growth and viability on microcarriers were assessed through daily sampling from the bioreactor, with cell counts obtained from representative samples from the bioreactor. Cell growth and proliferation was also monitored by brightfield microscopy and nucleic acid staining using DAPI **(Fig 4-b)**. By day 3, microcarrier aggregation due to cell bridging was observed, prompting an increase in agitation speed to maintain microcarriers in suspension and ensure homogeneous distribution. Metabolites levels were monitored throughout the culture, and on day 4, a 50% (5L) batch medium exchange was performed to replenish nutrients in the consumed culture medium **(Sup Fig 10-a)**. Following medium exchange, a marked increase in cell density was observed, reaching a plateau at approximately 7.5 × 10^5^cells/mL on day 6 onwards **(Fig 4-c)**.

**Figure 4.**
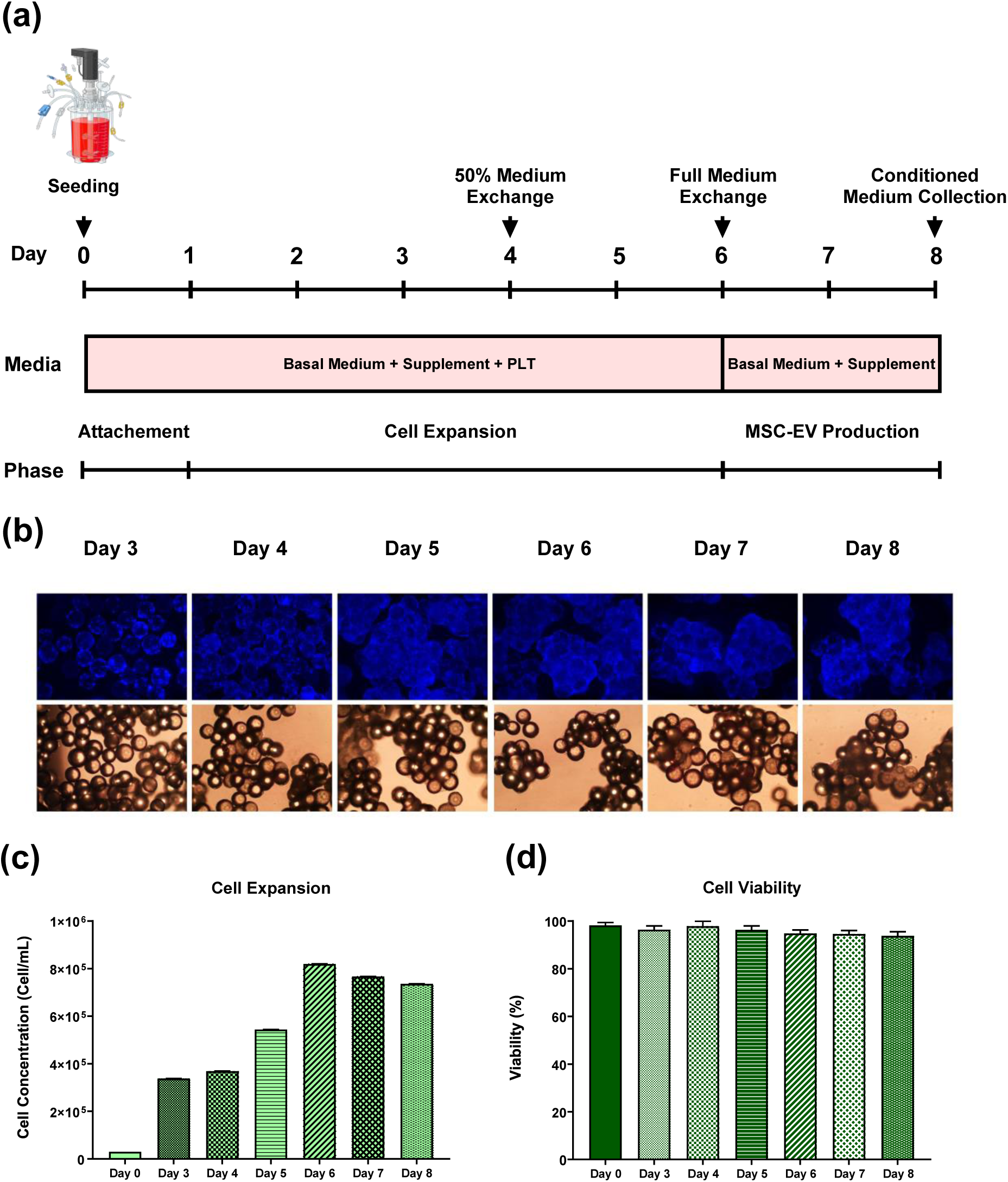
**Upstream Processing (UPS);** Scalable bioreactor expansion of human bone marrow mesenchymal stromal cells (hBM-MSCs) for EV production. **(a)** Schematic of the upstream culture process showing process steps, medium composition, and culture phases. hBM-MSCs were seeded at 3,000 cells/cm² onto microcarriers in a 10 L single-use bioreactor. Cultures were maintained in NutriStem® XF MSC basal medium supplemented with NutriStem® XF supplement and platelet lysate (PLT) for 6 days, with a 50% medium exchange on day 4. On day 6, the PLT-containing medium was fully replaced with PLT-free medium for EV production, and MSCs conditioned medium (MSCs CM) was harvested on day 8. **(b)** Representative fluorescence microscopy (DAPI nuclear staining, top row) and brightfield images (bottom row) from days 3–8 showing cell attachment, proliferation, and microcarrier aggregation. **(c)** Cell concentration increased steadily until plateauing at ∼7.5 × 10^5^ cells/mL by day 6. **(d)** Cell viability remained consistently above 90% for the entire culture period. All data are presented as mean ± SD with n 3 independent measurements. Schematics were created in Biorender.com.

On day 6, cellular aggregation was observed, and aggregates were allowed to settle at the bottom of the bioreactor by cessation of the agitation. The culture medium was then removed, and the cells were washed twice with Dulbecco’s phosphate-buffered saline (dPBS) to eliminate residual components from full medium. The bioreactor was then replenished with fresh, PLT-free medium (basal medium + supplement), to ensure that any EVs subsequently collected were exclusively secreted from the MSCs and not from exogenous PLT supplement. The culture was continued for an additional 2 days to allow for EV production and secretion into the medium. Cell viability remained consistently above 90% throughout the entire bioreactor run **(Fig 4-d)**. Additionally, over 97% of the cell populations were positive for CD73, CD90 and CD105 while 2.5% were positive for CD34 **(Sup Fig 10-b)**. On the other hand, the cells did not express the negative markers such as CD45 (0.01%), CD3 (0.07%) and HLA DR (0.087%) (Dominici *et al*., 2006).

### Downstream Processing

Downstream processing (DSP) typically consists of multiple unit operations which normally start off with a clarification step. Clarification is a normal flow filtration step in which the feed solution passes perpendicularly through a membrane and feed components larger than the pore size are retained. The clarification step is implemented in the DSP to remove larger particulates such as live cells, dead cells, cell debris and larger vesicles serving the same function as low-speed centrifugation steps used in ultracentrifugation. In this study, we utilized a two-step clarification strategy using sequential filtration through 5 µm and 0.65 µm pore size filters. Both steps were conducted at a constant flow rate of 100 mL/min, corresponding to a flux rate of 462 and 334 Liters per square meter per hour (LMH). Flux rate is an important parameter to assess the efficiency and performance of filtration operations. A total of 100 mL of MSCs CM was first clarified, while the pressure was monitored. Throughout the filtration, pressure remained stable with no noticeable increase, indicating the absence of membrane fouling or clogging (Data not shown). Unless otherwise specified, all samples in this study underwent two-step clarification as the first unit operation in DSP.

Ion Exchange Chromatography (IEX) is a common unit operation in DSP that isolates molecules based on their charge. Since EVs have a negatively charged surface, they are first captured on the cationic platforms like membranes, monoliths or resins, by electrostatic interaction and then eluted by increasing the ionic strength (salt concentration) of the elution buffer (Heath *et al*., 2018; Koch *et al*., 2023; Malvicini *et al*., 2023). We evaluated the performance of a membrane chromatography device, Sartobind Convec® D, which has been successfully shown for purification of lentiviruses for isolating MSC derived EVs (Pamenter *et al*., 2024). A total of 100 mL of clarified MSCs CM was loaded on the column first using a chromatography instrument equipped with RT-MALS detector to enhance process analytical monitoring. Upon sample loading, an increase in UV absorbance signal (∼1500 mAU) was observed with almost no RT-MALS signal. This signal is also referred to as flow through signal (FT) which are molecules that do not bind or interact with the column **(Fig 5-a)**. The loading phase was then followed by a wash step to remove unbound molecules, during which both UV and RT-MALS signals returned to baseline, confirming effective clearance. We then proceeded to the elution step by integrating a linear gradient strategy ranging from 0 to 1 M NaCl. During the elution phase, the RT-MALS detector revealed two partially overlapping (non-resolved) peaks, while the UV detector showed a high absorbance signal corresponding to the first peak, and a small signal for the second. Additionally, both elution peaks also happened below 400 mM salt concentration **(Fig 5-a)**.

**Figure 5.**
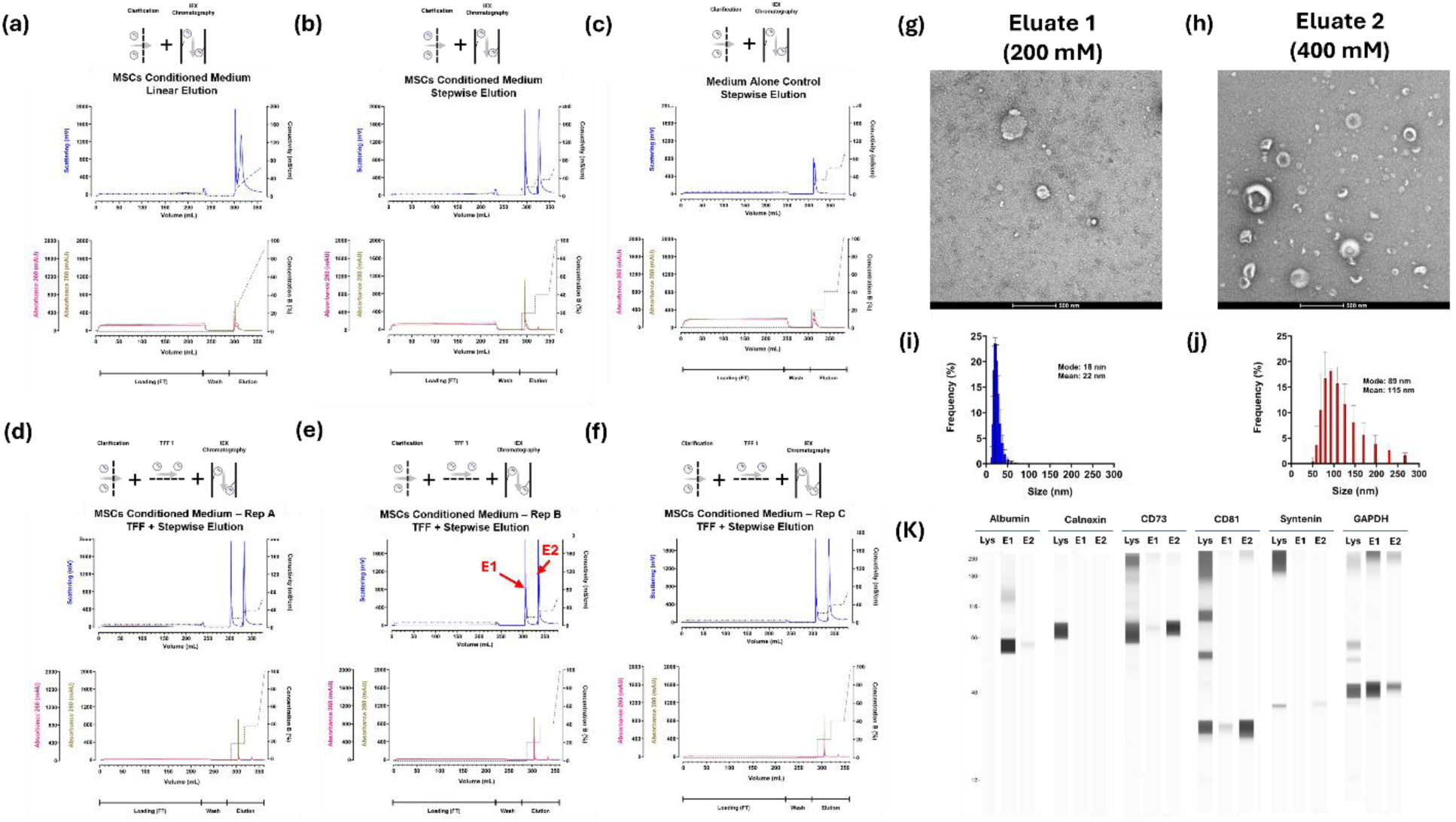
**Downstream Processing (DSP);** Development and validation of a robust DSP workflow for the purification of MSC derived extracellular vesicles (MSC-EVs) using ion exchange chromatography (IEX). **(a)** Real time monitoring of linear salt elution strategy using clarified MSCs conditioned medium (MSCs CM) on a membrane-based IEX column. Linear gradient elution revealed two overlapping peaks on RT-MALS detector with high UV absorbance for the first peak and weaker absorbance for the second. **(b)** Stepwise elution strategy (200 mM followed by 400 mM NaCl) improved resolution of the two eluate peaks, yielding well-separated populations with comparable RT-MALS signal intensity. The first peak exhibited strong UV absorbance, while the second had minimal UV signal, suggesting differences in composition. **(c)** Medium alone control subjected to the same stepwise elution protocol revealed signal only in the 200 mM eluate, indicating the presence of medium-associated components co-eluting in the first eluate, while no signal was observed at 400 mM step. **(d-e)** Reproducibility of the full DSP workflow was assessed across three independent DSP runs. All runs showed consistent and resolved dual elution profiles with strong RT-MALS signals and reproducible UV intensities, confirming process robustness. **(f–g)** TEM images of the first (200 mM) and second (400 mM) eluates. The 400 mM eluate 1 showed vesicular structures with characteristic cup-shaped morphology typical of EVs, while the 200 mM eluate 2 lacked cup-shaped morphology and displayed contaminant-enriched background and non-vesicular features. Scale bars: 500 nm. **(i–j)** DLS analysis showing size distribution profile of eluate 1 and 2. All data are presented as mean ± standard deviation with n 3 of independent DSP runs. **(k)** Simple Western analysis of both eluates alongside cell lysate control using positive and negative markers. EV-associated protein, showed faint bands in the second eluate and cell lysate, showing low expression levels. Schematics were created in Biorender.com.

To improve the resolution between the two eluate peaks observed during the linear gradient, we modified the elution strategy to a stepwise salt elution, in which salt concentration increases step by step. Specifically, we implemented a 200 mM salt concentration step for 10 mL, followed by a second 10 mL step at 400 mM. To further minimize cross-contamination between the two eluting populations, a 20 mL wash was incorporated after each elution step. A total of 100 mL clarified MSCs CM was then loaded under the new step by step elution strategy **(Fig 5-b)**. The RT-MALS detector revealed two fully resolved peaks with similar intensity (∼2 V) at 200 and 400 mM salt concentrations, indicating effective separation of the two populations. UV detectors showed a strong absorbance signal for the first peak (approximately 1100 mAU at 280 nm and 800 mAU at 260 nm), while the second peak exhibited minimal absorbance (< 40 mAU) **(Fig 5-b)**.

To determine whether the two populations are extracellular or cell-secreted components, we performed the same chromatographic procedure for unconditioned medium (medium alone control) as a process control. We loaded 100 mL of clarified medium alone control and proceeded with stepwise elution protocol. Notably, both RT-MALS and UV signals were detected during 200 mM salt elution step, while no detectable signal was observed at 400 mM salt eluate **(Fig 5-c)**. These findings indicate that the first eluate observed in MSCs CM contains a mixture of cell-secreted biomolecules and medium-associated components **(Fig 5-b)**. This result shows the importance of including medium alone controls as procedural control.

Tangential flow filtration (TFF) is a widely used unit operation in DSP, in which the feed stream flows tangentially across the membrane surface. This flow configuration helps to minimize membrane fouling by sweeping away retained molecules on the membrane and preventing cake formation. TFF can be applied to various process applications including concentration, ultrafiltration, microfiltration, diafiltration, buffer exchange and dialysis and have been widely used in EV processing (Busatto *et al*., 2018; Kim *et al*., 2021; Lei *et al*., 2025; Liu *et al*., 2025; McNamara *et al*., 2018).

Next, we incorporated a TFF step post clarification and prior to chromatography to remove medium-associated components. Additionally, we incorporated a nuclease treatment step with 1 hour incubation with Benzonase prior to TFF to digest free-floating host cell DNA, RNA and chromatin aggregates (Von Elling-Tammen *et al*., 2025). We performed six consecutive cycles of diafiltration in which, PBS buffer was added to the retentate to restore the original volume (100 mL) after each cycle. The TFF process was conducted under controlled pressure conditions as outlined in materials and methods. Transmembrane pressure (TMP) is defined as the pressure difference between the feed side and the permeate side of the membrane and is considered a critical parameter for optimizing TFF performance. This TMP value was selected after performing a TFF trial (Data not shown). We first performed this TFF step on clarified medium alone control. The resulting 6X diafiltered retentate was then loaded onto the IEX column. We observed no detectable RT-MALS and UV signals during elution, suggesting effective removal of medium-associated components by the incorporated TFF step prior to IEX chromatography **(Sup Fig 11-[a-c])**.

We next applied the full downstream processing workflow including clarification, TFF and IEX chromatography to 100 mL of MSCs CM. The 6x diafiltered MSCs CM was loaded onto the IEX column using the previously optimized stepwise elution strategy. As observed in the earlier experiments, we found two distinct and well-resolved peaks on RT-MALS detector with comparable intensity, while UV detector again showed a higher absorbance signal for the first eluate peak relative to the second **(Fig 5-d)**. To further assess the efficiency of the capturing of the extracellular components with IEX chromatography, we introduced an additional control by reloading the collected flow-through (FT), representing unbound components from the initial load, onto the column **(Sup Fig 11 – e and f)**. This FT rerun showed no detectable signal in either UV or RT-MALS detectors, suggesting high binding efficiency during the initial column loading step of MSCs CM **(Sup Fig 11 – e and f)**. To further evaluate process reproducibility, the entire workflow was repeated two additional times (n 3 independent DSP runs) on 100 mL of MSCs CM **(Fig 5-[d-f])**. Across all runs, consistent elution profiles were obtained on both RT-MALS and UV detectors. All control experiments including buffer alone, medium alone and FT rerun, consistently showed no detectable signal **(Sup Fig 11)**. These results confirm the robustness and reproducibility of the process and further support the two observed populations representing cell-secreted components efficiently captured and eluted by the IEX column **(Sup Fig 11-i)**.

### Identity Investigation

#### Biophysical Analysis

To further investigate the identity and composition of the two eluates, we performed transmission electron microscopy (TEM), dynamic light scattering (DLS) and Simple Western analysis. TEM imaging revealed that the second eluate (400 mM NaCl) was enriched in vesicular structures with the characteristic cup-shaped morphology commonly associated with EVs **(Fig 5-h)**. In contrast, the first eluate (200 mM NaCl) lacked cup-shaped morphology and displayed a more contaminant-rich background and non-vesicular features **(Fig 5-g)**. More TEM images can be found in **(Sup Fig 12-a and b)** at two different magnifications. For further validation, the equivalent eluates from the medium alone control were also imaged by TEM and showed no detectable structures, confirming the absence of cell-secreted components. **(Sup Fig 13-a)**. Dynamic Light Scattering (DLS) analysis showed that the second eluate exhibited a typical EV size distribution with mean size of ∼115 nm and mode size of ∼90 nm, while the first eluate contained predominantly smaller particles with mean size of ∼25 nm and mode of ∼20 nm **(Fig 5-i and j)**.

#### Marker Analysis

Simple Western provided further insight into presence and abundance of specific markers in the two eluates **(Fig 5-k)**. Albumin, a well-known contaminant and abundant serum protein, was enriched in the first eluate. Calnexin, an endoplasmic reticulum marker used as a negative control for EV purity, was detected in cell lysate control but was absent in both IEX eluates, suggesting minimal cellular contamination. CD73, a known marker of MSCs, was highly enriched in the second eluate and was also detected in cell lysate, supporting the MSC origin of this eluate. CD81, a common marker of EVs, was detected in the second eluate, with a faint band also observed in the first eluate. Syntenin, another EV-associated protein, showed weak expression, with a faint band observed only in the second eluate and cell lysate **(Fig 5-k)**. Consistent with these findings, Simple Western of the second eluate from the medium alone control showed no detectable signal for any of the markers tested, including EV-associated and contaminant markers **(Sup Fig 13-b)**. These results further confirm that the vesicular content observed in the MSCs CM is of cell-secreted origin and not derived from the medium formulation.

Additionally, we investigated RNA cargo of the samples to better understand compositions of eluates 1 and 2. EVs have been demonstrated to encapsulate a wide range of coding and non-coding RNAs, contributing to the diverse bystander effects they exert on recipient cells (Kim *et al*., 2017; Fabbiano *et al*., 2020; Dellar *et al*., 2022; Miceli *et al*., 2024). Messenger RNAs (mRNAs) have been consistently identified within EVs and these mRNAs were found to retain their functionality, suggesting their potential role in modulating cellular response in recipient cells. Recent studies have shown that RNA packaging into EVs is a selective process, influenced by the repertoire of RNA-binding proteins, the recruitment of endosomal release pathways and secretory autophagy (Fabbiano *et al*., 2020; Leidal and Debnath, 2021; O’Grady *et al*., 2022). To characterize the molecular identity of MSC-derived EVs, we selected mRNAs encoding well-established MSC markers - *THY1* (Thy1/CD90), *ENG* (Endoglin/CD105), and *NT5E* (ecto-5′-nucleotidase/CD73) - and quantified their expression in extracellular environment (Billing *et al*., 2016; L. Ramos *et al*., 2016; Nombela-Arrieta *et al*., 2011).

Our rationale was that most RNA species, including mRNAs, due to selective packaging into EVs with the involvement of secretory autophagy, ESCRT (endosomal sorting complex required for transport) pathway, utilization of packaging signals, would be detected in higher abundance within vesicular particles (eluate 2) in comparison to their non-vesicular particles counterparts (eluate 1) (Leidal *et al*., 2020; Fabbiano *et al*., 2020; Leidal *et al*., 2021; Oka *et al*., 2023). Reverse Transcription quantitative PCR (RT-qPCR) analysis revealed that eluate 2 was significantly enriched in all three mRNA species compared to eluate 1 **(Sup Fig 13-c)**. Additionally, *GAPDH* mRNA, a commonly used housekeeping gene, also showed increased presence in the eluate 2. As shown in **(Sup Fig 13-c)**, the eluate 2 showed a significantly higher quantity of mRNA content, suggesting increased mRNA encapsulation within these vesicles when compared to eluate 1. In summary, our data suggested that eluate 2 had increased the quantity of MSC specific mRNAs and it was likely enriched in vesicular particles that allowed encapsulation of RNAs.

#### Tetraspanins Profiling

In addition to proteomic and lipidomic analysis, we studied the expression level of common Tetraspanins (CD9, CD63 and CD81) between MSCs CM, eluate 1 and 2 using direct stochastic optical reconstruction microscopy (dSTORM). Super resolution microscopy has been recently employed for single particle level characterization of EVs derived from body fluids such as amniotic fluids (Gebara *et al*., 2022), and MSCs CM (Gebara *et al*., 2022; Salem *et al*., 2024). A larger number of clusters were observed in MSCs CM and the two eluates compared to the applied control samples, such as eluate 2 of medium alone, dye alone control and unlabeled eluate 2 **(Sup Fig 14-a and b)**. The number of detected clusters was significantly higher in eluate 2 compared to eluate 1 and MSCs CM **(Sup Fig 14-a and b)**. In terms of size, eluate 2 was larger followed by MSCs CM and eluate 1 **(Sup Fig 14-c)**. The expression profile of CD9, CD63 and CD81 was different between samples **(Sup Fig 14-[d-f])**. Eluate 2 showed 59% triple positivity, while eluate 1 had 43% triple positivity. On the other hand, eluate 1 showed 34% double positivity for CD63 and CD9, while only 11% were double positive in eluate 2. In total, around 85% of the clusters detected in eluate 2 were positive for CD81. Representative images of individual clusters detected in MSCs CM, eluate 1 and 2 can be found in **(Sup Fig 14-[g-f])**.

#### Protein Composition Analysis

To evaluate protein composition of the eluates 1 and 2, we executed proteomics analysis of the eluates, MSC CM and medium alone. First, we measured the total protein content of the two eluates and eluate 1 showed almost 10 times higher total protein concentration compared to eluate 2 **(Fig 6-a)**. Next, we sought to characterize the proteomic content of the samples. We performed liquid chromatography-mass spectrometry (LC-MS) on both eluates, as well as medium alone and MSCs CM. As a quality control measure, we compared both eluates with the medium alone and we detected fewer than 400 unique protein signatures in the medium alone (data not shown). Next, we compared the protein content of both eluates and detected over a total of 1767 protein signatures **(Fig 6-b)**. Among these, 300 proteins were unique for eluate 1, while 483 were overrepresented in eluate 2, indicating differential protein signatures subsets between the two eluates **(Fig 6-b)**. The remaining 984 signatures were found in both eluates, suggesting the protein population that persists across the two populations. To evaluate the reproducibility of our findings, we analyzed three replicates of both eluates. Venn diagrams **(Sup Fig 15-a)** showed that three replicates of eluate 1 shared 1284 proteins between each other, while 47-131 proteins were replicate-specific. Three replicates from eluate 2 had 1467 common protein signatures, while 60-91 proteins were replicate-specific. Overall, experiments showed the presence of an essential set of proteins in each eluate, along with a relatively small number of replicate-specific protein signatures, that can be explained by inherent variability of LC-MS (Piehowski *et al*., 2013).

**Figure 6.**
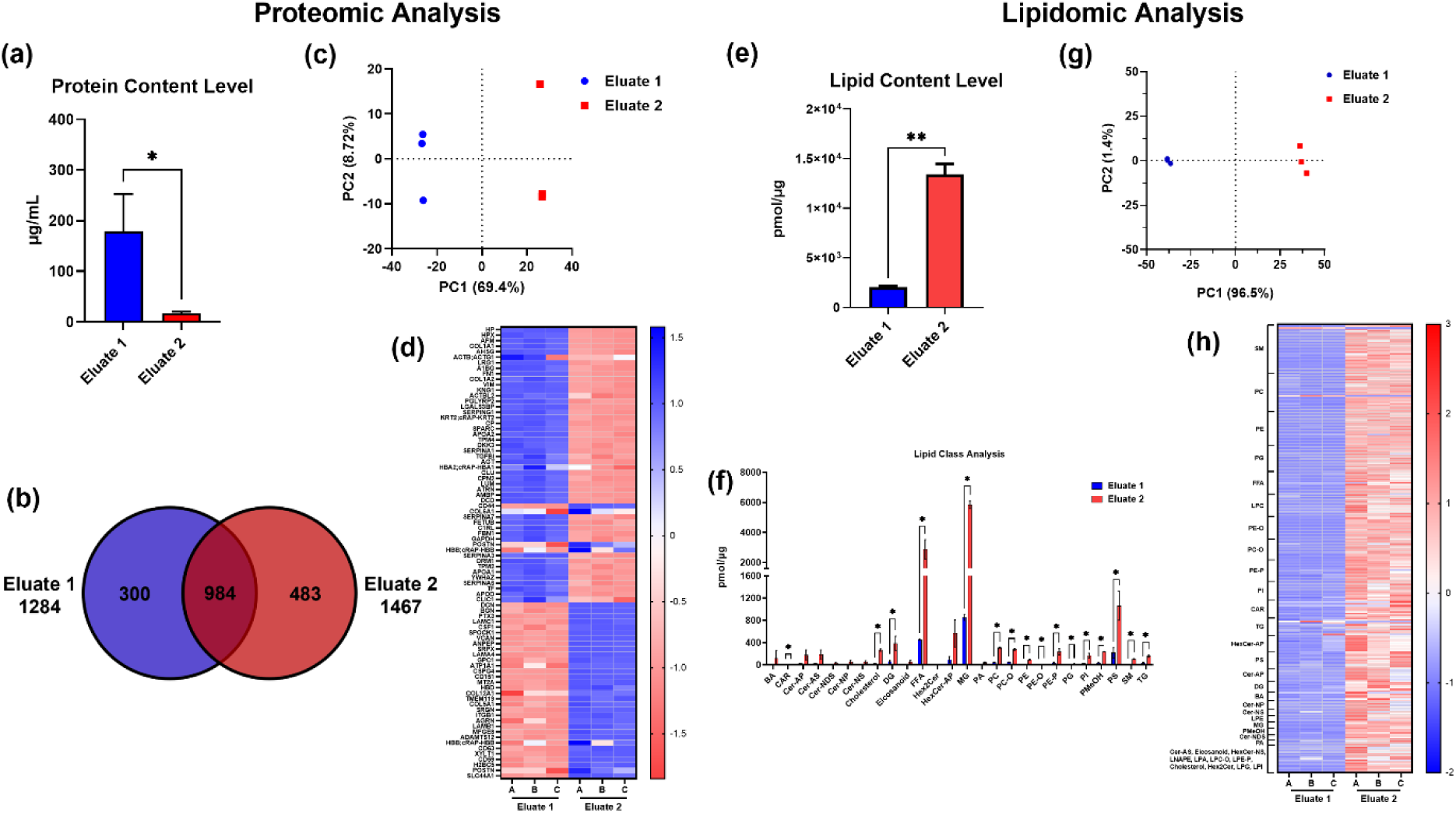
**Identity Investigation;** Comparative proteomic and lipidomic profiling of the two eluates from MSCs conditioned medium (MSCs CM). **(a-d)** Proteomic analysis; **(a)** Total protein concentration measured by BCA assay showed that eluate 1 contained ∼10-fold higher protein content compared to eluate 2. **(b)** Venn diagram depicting the overlap and unique protein signatures between the two eluates. **(c)** Principal Component Analysis (PCA) of LC-MS data demonstrated clear separation of the two eluates along PC1 (69.4% variance) and PC2 (8.72% variance). **(d)** Heatmap of the top 50 most abundant proteins across the two eluates, highlighting distinct expression patterns. **(e-h)** Lipidomic Analysis; **(e)** Total lipid content showed that eluate 2 contained ∼6.5-fold higher lipid content compared to eluate 1. **(f)** Lipid class composition in each eluate. **(g)** Lipidomics PCA indicated a clear separation between eluates along PC1 (96.5% variance) **(h)** Heatmap of lipid species abundance across the two eluates, highlighting distinct expression patterns.

To characterize the variance between eluate 1 and 2, we performed Principal Component Analysis (PCA) using log_2_ intensity values. As a result, the plot **(Fig 6-c)** showed a clear separation between the two eluates along PC1, which accounts for 69.4 % of the total variance and PC2 corresponded to 8.72% of the variance. The separation indicates distinct protein composition between the two eluates, consistent with a Venn diagram in **(Fig 6-b)**. Next, we characterized the differential expression of the proteins with the use of *limma* package (Gentleman *et al*., 2004). We visualized the expression profile of the top 50 most abundant proteins found across two eluates via a heatmap **(Fig 6-d)**. The heatmap shows a relative abundance of the protein signatures across two eluates, with color intensity (blue for low, red for high) indicating relative intensity of the protein signatures. The differential protein expression profiles observed in the two eluates provide additional evidence of the involvement of separate secretion pathways underlying the release of these particle populations. Additionally, we performed a comparative proteomics analysis of the top 50 differentially expressed proteins between the two eluates. The heatmaps **(Sup Fig 15-c)** illustrate the top 50 differentially expressed proteins, upregulated in eluate 1 relative to eluate 2, and vice versa, underscoring the specific protein signature profiles of both eluates.

To visualize the distribution of intensities of signatures of the proteins of interest we generated protein rank plots for each of the three samples including MSCs CM, eluate 1 and 2 **(Sup Fig 15-b)**. As a result, the intensity distribution showed a broad dynamic range varying from 10^3^ to 10^11^, indicating a complex mixture of secreted proteins. To visualize the changes of proteomic profiles after chromatographic separation we decided to plot the proteins that have been used as exosome markers or previously been detected in extracellular vesicles (EVs) and non-vesicular particles (NVEPs) by other groups such as CD63, CD81, CD82, CD9, PDCD6IP/Alix, TSG101, LGALS3BP, FGA and PGK1 (Welsh *et al*., 2023). Endoplasmic reticulum marker calreticulin (CALR) and albumin (ALB) were used as the control for cell debris and free-floating proteins respectively **(Sup Fig 15-b)** (Shin *et al*., 2017; Jepessen *et al*., 2014; Ter-Ovanesyan *et al*., 2021; Ekstrom *et al*., 2022; Welsh *et al*., 2023).

Eluate 1 displayed a narrower intensity range from 10^3^ to 10^9^, with albumin and LGALS3BP still dominating the top ranks **(Sup Fig 15-b)**. However, despite being detected previously, CD9 and CD82 appeared at the lower ranks in comparison to MSCs CM, while albumin intensity showed significant drop, suggesting successful enrichment of specific protein populations more abundant in eluate 1. Lastly, eluate 2 showed an even narrower intensity range between 10^3^ to 10^9^, with CD63 appearing closer to the top of the rank and CD81 and CD9 moving closer to the top and TSG101 becoming detectable at the bottom of the rank **(Sup Fig 15-b)**. These proteins are routinely used as exosome markers and their increased intensity in the ranks indicates a different enrichment profile corresponding to vesicular particles (Welsh *et al*., 2023). Interestingly, LGALS3BP, that has been previously shown to be enriched in non-vesicular extracellular particles such as exomeres, showed increased intensity in eluate 1 in comparison to MSC CM (Zhang *et al*., 2018; Zhang *et al*., 2019; Zhang *et al*., 2021, Tosar *et al*., 2022). Overall, the rank plots show differential protein signatures intensities across the samples, reflecting the successful chromatographic elution. Additionally, we compared the relative abundance of the common EV markers in eluates and as a result, TSG101, PDCD6IP/Alix, CD81, CD82, CD63 and CD9 showed increased expression in E2 in comparison to E1 (**Sup Fig 15-e**). Interestingly, LGALS3BP showed increased expression in E1, suggesting a possible enrichment in non-vesicular particles (Tosar *et al*., 2022; Zhang *et al*., 2018).

To elucidate the biological origin of eluates, we performed Gene Set Enrichment Analysis (GSEA) **(Sup Fig 15-d)**. eluate 1 showed enrichment in components of protein-lipid complexes and their receptors, including LDLR, LCAT, APOA4, and APOH, all of which are well-established constituents of lipid-protein assemblies (Mehta *et al*., 2022; Ma *et al*., 2024). Additionally, proteins associated with high-density lipoprotein (HDL) particles - such as APOE, PLTP, PON1, and LCAT - were also enriched in eluate 1. Furthermore, eluate 1 showed enrichment in proteasome complex components (PSMB7, PSMA1, PSMA4, PSMB5, PSMB4), consistent with previous findings by other research groups (Ben-Nissan *et al*., 2022; Bonhoure *et al*., 2022; Kim *et al*., 2025). We also detected components of endopeptidase complexes, including CFH and CAPN2, which have been previously reported in EVs of diverse origins (Bushey *et al*., 2021; Dong *et al*., 2025). Interestingly, cellular components associated with blood microparticles - such as CPN2, SERPINF2, APOA4, ORM1, and SERPINA3 - were also present in eluate 1 **(Sup Fig 15-d)** (Zhang *et al*., 2016; Xu *et al*., 2022; Meng *et al*., 2022).

In contrast, eluate 2 samples were predominantly enriched in the proteins involved in vesicles membrane formation such as APLP2, CD46 and RAB35 (Lee *et al*., 2023) **(Sup Fig 15-d)**. Basal components of the sarcolemma, such as BGN and various solute carriers (SLCs), were also identified in eluate 2, consistent with their routine detection in EVs reported in previous studies (Deng *et al*., 2023; Zhou *et al*., 2018; Console *et al*., 2018; Hirpara *et al*., 2024). Additionally, eluate 2 were enriched in membrane transport proteins involved in endosomal membrane recycling - such as RAB35, SCAMP3 and VAMP3 - further supporting the notion that the second eluate may originate from ESCRT-dependent secretion pathways (Sheehan *et al*., 2016; Kumar *et al*., 2023; Thomas *et al*., 2016; Fader *et al*., 2009). Basolateral membrane components such as ENPP1 and ATP2B4 were also detected in eluate 2 **(Sup Fig 15-d)**.

#### Lipid Composition Analysis

Next, we investigated the lipid profile and composition of the two eluates using quantitative lipidomics. Ultra performance liquid chromatography-tandem mass spectrometry was performed on both eluates. The analysis of the number of identified lipids in each subclass showed similar distribution of lipid species in the two eluates with Sphingomyelin (SM) and Phosphatidylcholine (PC), Free Fatty Acids (FFA), Phosphatidylethanolamine (PE) and Phosphatidylglycerol (PG) as the dominant classes **(Sup Fig 15-f)**. Principle component analysis (PCA) was performed on all the samples to examine the overall lipid differences between the two eluates and the variation within each eluate. PCA revealed clear separation between the two eluates along the first principal component (PC1, 96.5% variance), with minimal variance observed along PC2 (1.4%) **(Fig 6-g)**.

Analysis of the total lipid abundance showed that eluate 2 (∼13387 pmol/µg on average) had over 6.5-fold total lipids compared to eluate 1 (∼2039 pmol/µg on average). The lipid class analysis and quantification revealed more details on lipid composition of the two eluates. Amongst the lipid classes, MG was the predominant subclass in both eluates, followed by FFA, and Phosphatidylserines (PS). The heatmap for both eluates can be found in (**Sup Fig 15-i)** indicating the relative quantification of each lipid class. To further evaluate compositional differences between eluates, we performed a lipid class ratio analysis by calculating the abundance of each lipid class in eluate 2 (E2) relative to eluate 1 (E1). The resulting bar plot **(Sup Fig 15-g)** illustrates the lipid abundance ratio (E2/E1) in detected lipid species across all identified lipid classes. A blue dashed line indicates the total lipid abundance ratio between E2 and E1, calculated to be 6.5, serving as a reference threshold for enrichment. Several lipid classes exhibited enrichment above this threshold, indicating a relative accumulation in the eluate 2. Bile acid (BA) with ∼20-fold, Cholesterol with ∼12-fold higher abundance was found in eluate 2. Within glycerophospholipids lipid class, 14-fold enrichment for N-acyl-Lysophosphatidylethanolamine (LNAPE), 18-fold for PE, 21-fold for ether-linked Phosphatidylethanolamine (PE-O), 26-fold for Lysophosphatidylethanolamine (LPE) and 13-fold for Lysoalkenylphosphatidylethanolamine (LPE-P) was observed. Additionally, the top 20 differentially expressed lipids between eluate 1 and 2 are shown in (**Sup Fig 15-h)** with fold-change value shown as log_2_ values.

Eluate 2 was enriched by a factor of 6.5 in total lipid content normalized to total protein content, which reinforces that eluate 2 is enriched in EVs, consistent with analytical chromatography, particle count, and proteomic data. eluate 1 and 2 had similar lipid compositions, further indicating lipid signal in eluate 1 may be from EVs that co-eluted in eluate 1. Concerning lipid composition, FFA and MG accounted for a high mass percentage of total lipid content. In agreement with this, quantitative lipidomics have shown FFA to be approximately 35% of lipids detected in bone marrow MSC exosomes (Haraszti *et al*., 2016). In another study, quantitative lipidomics of human induced pluripotent stem cell derived blood vessel organoid EVs identified MG and FFA as the most abundant lipid subclasses (Ene *et al*., 2025). Stearic acid and palmitic acid (the 2 most abundant FFAs in the data) alone or as fatty acid residues have been shown to be packaged into or co-isolated with bone marrow and adipose tissue MSC-EVs *(Casati et al*., 2022), including at high relative peak intensity (Showalter *et al*., 2019). Additionally, we used a serum free medium during our EV production phase, which has been shown to result in FFA enrichment compared to serum containing medium in umbilical cord, bone marrow, and adipose MSC microvesicles (Haraszti *et al*., 2019).

We observed numerous lipid species in the subclasses SM, PC, PE, and PG in line with broad evidence that these are major lipid components of biological membranes and thus EVs (Skotland *et al*., 2020). Lastly, we observed the subclass TG in both eluates; although TG is not a membrane component (Skotland *et al*., 2025), it is contained in lipoproteins and lipid droplets which can be packaged inside or co-isolated with EVs, potentially explaining our results (Nikitidou *et al*., 2017). Overall, our findings agree with quantitative lipidomics in the literature and show that eluate 2 lipid composition supports enrichment of EVs.

### Downstream Process Monitoring

To comprehensively evaluate the performance of our downstream processing (DSP) workflow, we utilized the analytical toolbox established in this study. Particle concentrations were measured at each intermediate step using Sc-NTA, Fl-NTA, and Fl-FC. In parallel, total protein and DNA concentrations were quantified across all DSP steps for both MSCs CM and medium alone control.

Additionally, analytical chromatography—combining size exclusion separation with in-line MALS, UV, and fluorescence detection—was employed to assess the relative abundance of EVs versus soluble impurities. This orthogonal approach enabled a detailed evaluation of each unit operation’s performance to EV enrichment and impurity clearance throughout the process.

#### Clarification

All three single particle analysis techniques detected a modest reduction in particle concentration following clarification, likely due to the removal of larger particulates present in the MSCs CM. Specifically, particle retention rates post-clarification were 86% for Sc-NTA, 83% for Fl-NTA, and 83% for Fl-FC **(Fig 7-[a-c])**. Protein quantification using BCA assay showed a 14% reduction (86% recovery) in total protein content in MSCs CM and a 7% reduction (93% recovery) in the medium alone control **(Fig 7-[d and g])**. Analytical chromatography revealed MALS signals corresponding to EV size analytes (minutes 9-14, highlighted as red color), while UV absorbance and intrinsic fluorescence signals (minutes 14-30, highlighted as green and yellow) were predominantly associated with impurity fractions, consistent with the unpurified nature of the pre-processed samples **(Fig 8-[a-c])**. Quantitative analysis of the area under the curve (AUC) of intrinsic fluorescence for impurity-associated peaks (minutes 14-30) showed a 17% reduction in MSCs CM and a 13% reduction in medium alone control, corresponding to impurity recoveries of 83% and 87%, respectively **(Fig 7-[e and h] and Sup Fig 17-[a-c])**. Furthermore, DNA quantification using the PicoGreen™ assay showed a 16% reduction in DNA mass in MSCs CM following clarification, compared to a 3% reduction in medium alone control **(Fig 7-[f and i])**.

**Figure 7.**
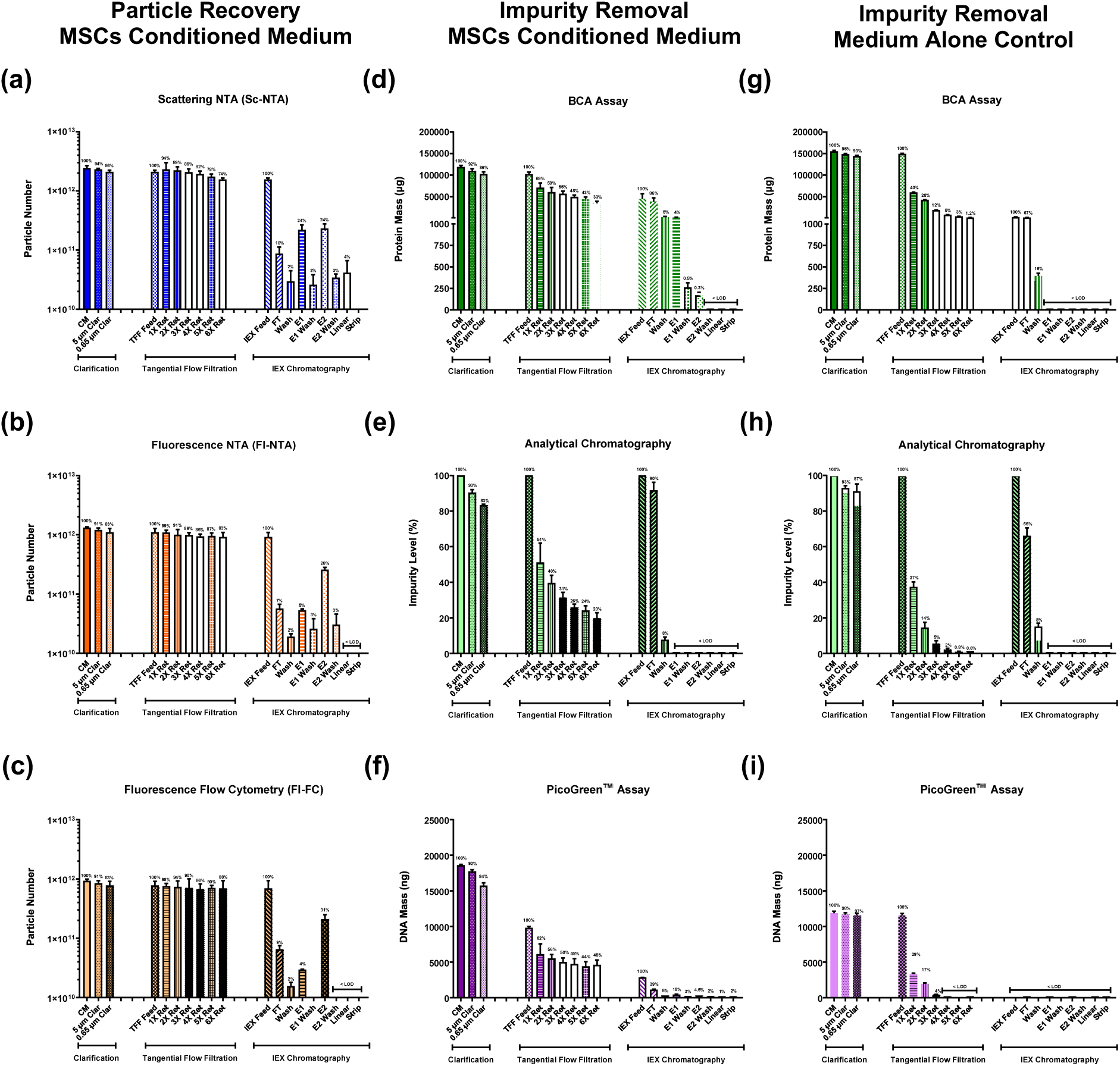
**Downstream Process (DSP) monitoring;** Step by step DSP evaluation using orthogonal analytics to assess particle recovery and impurity removal. **(a–c)** Particle recovery at each DSP step quantified by scattering mode nanoparticle tracking analysis (Sc-NTA), fluorescence mode NTA (Fl-NTA), and fluorescence-based flow cytometry (Fl-FC). **(d–f)** Impurity removal from MSCs CM assessed by total protein quantification (BCA assay), analytical chromatography (size exclusion separation with in-line MALS, UV absorbance, and intrinsic fluorescence detection), and DNA quantification (PicoGreen™ assay). **(g–i)** Impurity removal from medium alone control assessed using the same analytical toolbox. All data is presented as mean ± SD with n 3 independent DSP runs. The percentage values are based on the feed of each step of DSP (step recovery).

**Figure 8.**
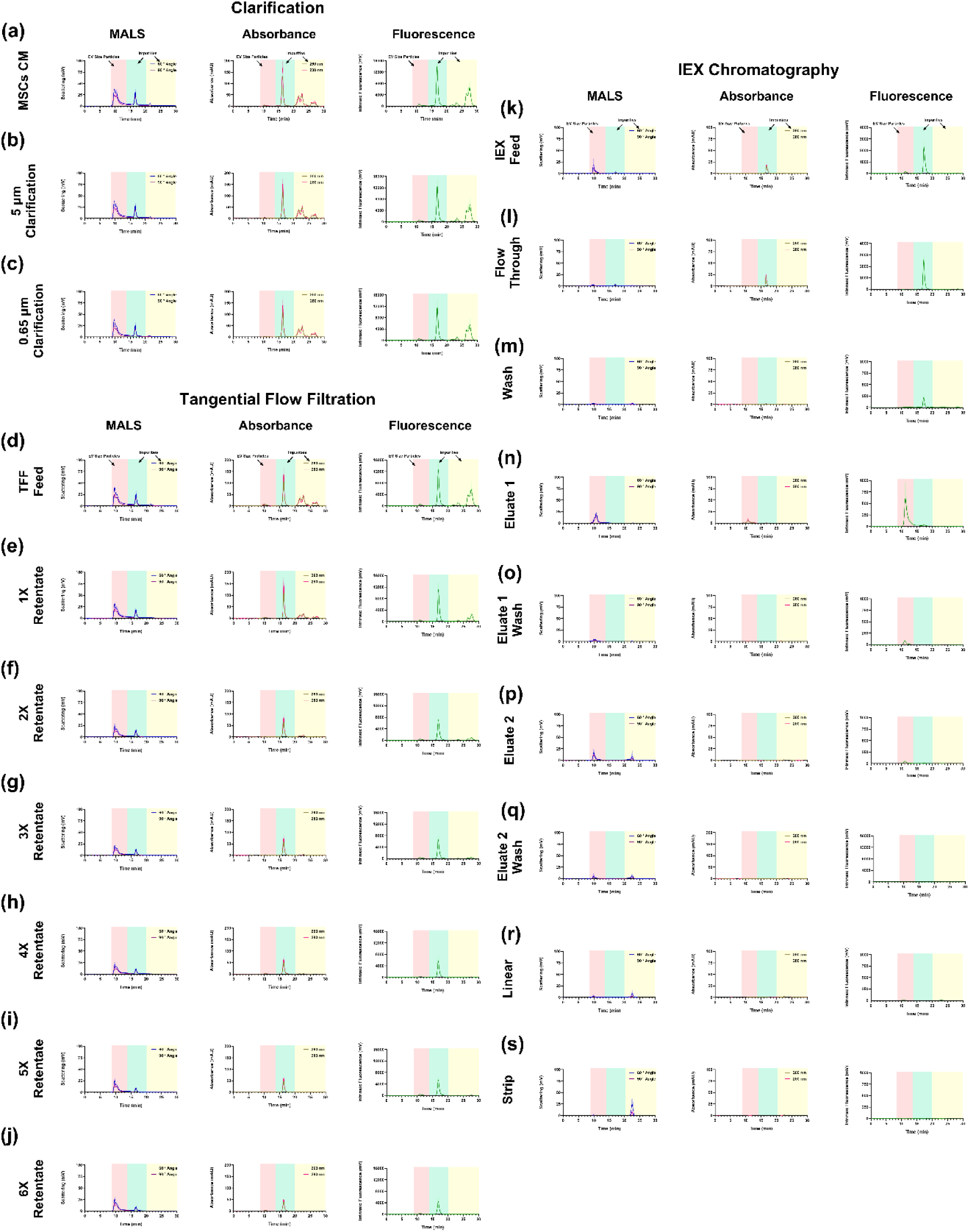
**Downstream Process (DSP) Monitoring;** Step by step DSP evaluation using multi-detector analytical chromatography. Size exclusion chromatography with in-line MALS, UV absorbance, and intrinsic fluorescence detection was used to evaluate particles and impurities distribution at each DSP stage. **(a–c)** Clarification step showing EV size particles peaks (9–14 min, highlighted in red) and impurity-associated peaks (14–20 min, highlighted in green) along with late-eluting smaller soluble impurities (>20 min, highlighted in yellow). **(d–j)** Six-cycle tangential flow filtration (TFF) removed ∼80% of impurities in clarified MSCs CM with complete elimination of late-eluting peaks (>20 min, highlighted in yellow) and major reduction of 14–20 min (highlighted in green) impurities. **(k–m)** Flow through (FT) and wash samples from IEX chromatography showed impurity peaks without EV-size particles signal, confirming efficient EV capture. **(n and p)** Comparison of eluates: both showed similar MALS signal intensity, but eluate 1 exhibited peaks in UV and fluorescence channels, while eluate 2 had minimal signals. Minimal signals were detected in all the other tested samples, such as the wash steps after eluate 1 **(o)** and eluate 2 **(q)** as well as linear (in which salt concentration was increased from 400 to 1000 mM) **(r)** and strip (in which salt concentration was kept as 1000 mM) **(s)**. All data are presented as mean ± SD with n 3 independent DSP runs.

#### Tangential Flow Filtration

Single particle analysis techniques demonstrated consistent trends throughout the 6× diafiltration process. Particle recovery rates were 74% by Sc-NTA, 83% by Fl-NTA, and 88% by Fl-FC, indicating efficient retention of particles across the TFF step **(Fig 7-[a-c])**. Protein quantification via BCA assay revealed that the diafiltration step significantly improved sample purity, with a 67% reduction in total protein content for MSCs CM and a 99% reduction in the medium alone control **(Fig 7-[d and g])**.

These findings were supported by analytical chromatography. Chromatographic profiles showed complete elimination of late-eluting peaks (after 20 minutes, highlighted as yellow) on both UV and intrinsic fluorescence detectors, while impurity-associated peaks eluting between 14–20 minutes were also successfully reduced (highlighted in green) **(Fig 8-[d-j])**. Quantitative analysis of the area under the curve for impurity fractions confirmed 80% impurity removal in MSCs CM and 99% in the medium alone control **(Fig 7-[e and h])**. As expected, chromatographic profiles for the medium alone control showed complete removal of all impurity peaks, validating the effectiveness of the TFF process in clearing medium-associated components **(Sup Fig 17-[d-j])**.

In addition, DNA quantification using the PicoGreen™ assay revealed a 55% reduction in DNA content for MSCs CM **(Fig 7-[f])**. DNA in the medium alone control was reduced below the lower LOD after diafiltration, further confirming the TFF step’s efficacy in removing DNA related contaminants **(Fig 7-[g])**. To evaluate the impact of benzonase treatment on host cell DNA content, we incorporated a nuclease digestion step into the downstream processing workflow. Across all three MSCs CM runs, benzonase treatment did not affect total protein concentration, as confirmed by BCA analysis **(Sup Fig 16-[g])**. Similarly, no change in protein content was observed in the medium alone control, indicating that measured concentration by PicoGreen™ assay was associated with DNA **(Sup Fig 16-[g])**. In contrast, DNA quantification revealed a significant reduction in DNA content following a 1-hour benzonase digestion in MSCs CM samples **(Sup Fig 16-[h])**. No measurable change in DNA concentration was observed in medium alone control, suggesting that the DNA targeted by benzonase was derived from cell-secreted components rather than medium-associated ones **(Sup Fig 16-[h])**. These results support the use of benzonase as an effective strategy for selectively removing host cell DNA without compromising protein or vesicle content.

#### Ion Exchange Chromatography

We evaluated the performance of the IEX chromatography step using the Convec® D membrane column by analyzing all intermediate samples with our orthogonal analytical toolbox. Single particle analysis revealed that 10%, 7%, and 9% of the feed particles were detected by Sc-NTA, Fl-NTA, and Fl-FC, respectively in the flow through (FT) **(Fig 7-[a-c])**. Protein quantification by BCA assay showed that 86% of the feed protein was present in the FT, consistent with the analytical chromatography area under the curve (AUC) analysis, which indicated 90% impurity remained in the FT **(Fig 7-[d and e])**. Chromatographic profiles of the FT confirmed the presence of the impurity peak (14–20 min, highlighted in green) and no signal in EV size fraction (9-14 min, highlighted in red) suggesting efficient capturing of larger particles such as EVs on the column **(Fig 8-[k and l])**. PicoGreen™ assay further showed that 40% of the DNA from the feed was detected in the FT **(Fig 7-[f])**.

The wash step following sample loading was similarly evaluated. All three single particle techniques detected only ∼2% of input particles in the wash, while BCA and AUC analysis showed 5% and 9% of the feed protein were found in the wash, respectively **(Fig 7-[a-e])**. A small UV absorbance and intrinsic fluorescence signal was observed in the impurity elution window (14–20 min, highlighted in green), indicating residual impurity clearance during the wash **(Fig 8-[m])**. PicoGreen™ assay showed 5% DNA content in the wash **(Fig 7-[f])**. Notably, all post-diafiltration medium alone control samples showed protein and DNA concentrations below the lower LOD, confirming effective impurity removal **(Fig 7-[g])**.

The wash step following sample loading was similarly evaluated. All three single particle techniques detected only ∼2% of input particles in the wash, while BCA and AUC analysis showed 5% and 9% of the feed protein were found in the wash, respectively **(Fig 7-[a-e])**. A small UV absorbance and intrinsic fluorescence signal was observed in the impurity elution window (14–20 min, highlighted in green), indicating residual impurity clearance during the wash **(Fig 8-[m])**. PicoGreen™ assay showed 5% DNA content in the wash **(Fig 7-[f])**. Notably, all post-diafiltration medium alone control samples showed protein and DNA concentrations below the lower LOD, confirming effective impurity removal **(Fig 7-[g])**.

Sc-NTA analysis revealed comparable particle concentrations in eluate 1 and 2, each containing approximately 24% of the feed particles loaded **(Fig 7-[a])**. These results aligned well with RT-MALS data obtained during IEX chromatography, in which both eluates exhibited a 2V MALS signal **(Fig 5-[d-f])**. Analytical chromatography also showed equivalent MALS signal intensity in the EV-size fraction window (9–13 minutes, highlighted in red) for both eluates, indicating that from a scattering-based perspective, the two eluates were similar **(Fig 8-[n and p])**.

In contrast, fluorescence-based methods (Fl-NTA and Fl-FC) revealed significant differences between the two eluates. Fl-NTA and Fl-FC showed higher particle recovery in eluate 2 (29% and 31%, respectively) compared to eluate 1 (5% and 4%) **(Fig 7-[b and c])**. This suggests that fluorescence-based quantification is more specific to EVs, although some non-specific labeling of non-vesicular particles and impurities as well as co-eluted EVs may contribute to residual signal in eluate 1. Notably, all three orthogonal techniques showed good concordance in particle concentrations for eluate 2, supporting its high purity of extracellular vesicles and the absence of similar size non-vesicular impurities **(Sup Fig 16-[e])**.

BCA analysis showed that eluate 1 contained 4% of the feed protein, whereas eluate 2 contained only 0.3% **(Fig 7-[d])**. These results were in strong agreement with analytical chromatography data, which showed a prominent UV and intrinsic fluorescence signal for eluate 1 and a very low signal for eluate 2 **(Fig 8-[n and p])**. This was consistent with real time UV detection during the IEX run, which recorded a much higher absorbance for eluate 1 compared to eluate 2 **(Fig 5-[d-f])**.

A comprehensive summary of particle concentrations across all DSP unit operations is provided in **(Sup Fig 16-[a-e])**. As expected, Sc-NTA consistently reported higher particle counts compared to Fl-NTA and Fl-FC, particularly in early process steps where non-vesicular particles are prevalent. However, as purification progressed, particle concentrations across the three techniques converged, indicating improved sample purity. BCA results confirmed higher protein content in eluate 1 (180 µg/mL) compared to eluate 2 (20 µg/mL), while the original MSCs CM protein concentration was 1.18 mg/mL. DNA content was measured as ∼40 ng/mL in eluate 1 versus 12 ng/mL in eluate 2, while MSCs CM showed 185 ng/mL **(Sup Fig 16-[f-i])**. Although DLS analysis indicated differences in size between the two eluates, Sc-NTA and Fl-NTA showed overlapping size distributions, likely due to limitations in resolving subtle structural differences between vesicular and non-vesicular particles **(Sup Fig 16-[j-m])**.

### Downstream Processing Scalability

To evaluate the scalability of the downstream processing (DSP) workflow for MSC-derived EVs, we increased the feed volume from 100 mL to 700 mL and adjusted each unit operation accordingly. Clarification was scaled using larger filter capsules: from size 4 to size 5, increasing the effective membrane area from 0.018 m² to 0.036 m² for the 5 µm filter, and from 0.013 m² to 0.026 m² for the 0.65 µm filter, respectively. A larger tangential flow filtration (TFF) cassette was used for diafiltration, increasing the membrane area from 0.02 m² to 0.14 m². The IEX chromatography column was also scaled from 3 mL to 20 mL membrane volume. To complete the workflow, a second TFF step (2× diafiltration) was added post-IEX to exchange the buffer and reduce NaCl concentration from 400 mM to physiological levels (PBS). A final sterile filtration step was incorporated to establish a complete and scalable DSP.

This scaled-up process consisted of five-unit operations: clarification, first tangential flow filtration, IEX chromatography, second tangential flow filtration and sterile filtration **(Fig 9-a)**. Orthogonal single particle analysis techniques, Sc-NTA, Fl-NTA, and Fl-FC showed particle recoveries of 81%, 83%, and 84%, respectively, after clarification step **(Fig 9-[b-d] and Sup Fig 18-[a and b])**. Additionally, clarification removed ∼4% of protein content (96% recovery), while no DNA reduction was observed by PicoGreen™ assay **(Fig 9-[e and g] and Sup Fig 18-[g and h])**. Chromatographic analysis showed the presence of impurity peaks (14–20 min, highlighted in green and 20–30 min, highlighted in yellow) on UV and intrinsic fluorescence channels **(Fig 9-j)**. AUC analysis confirmed a 9% impurity reduction during clarification **(Fig 9-f)**.

**Figure 9.**
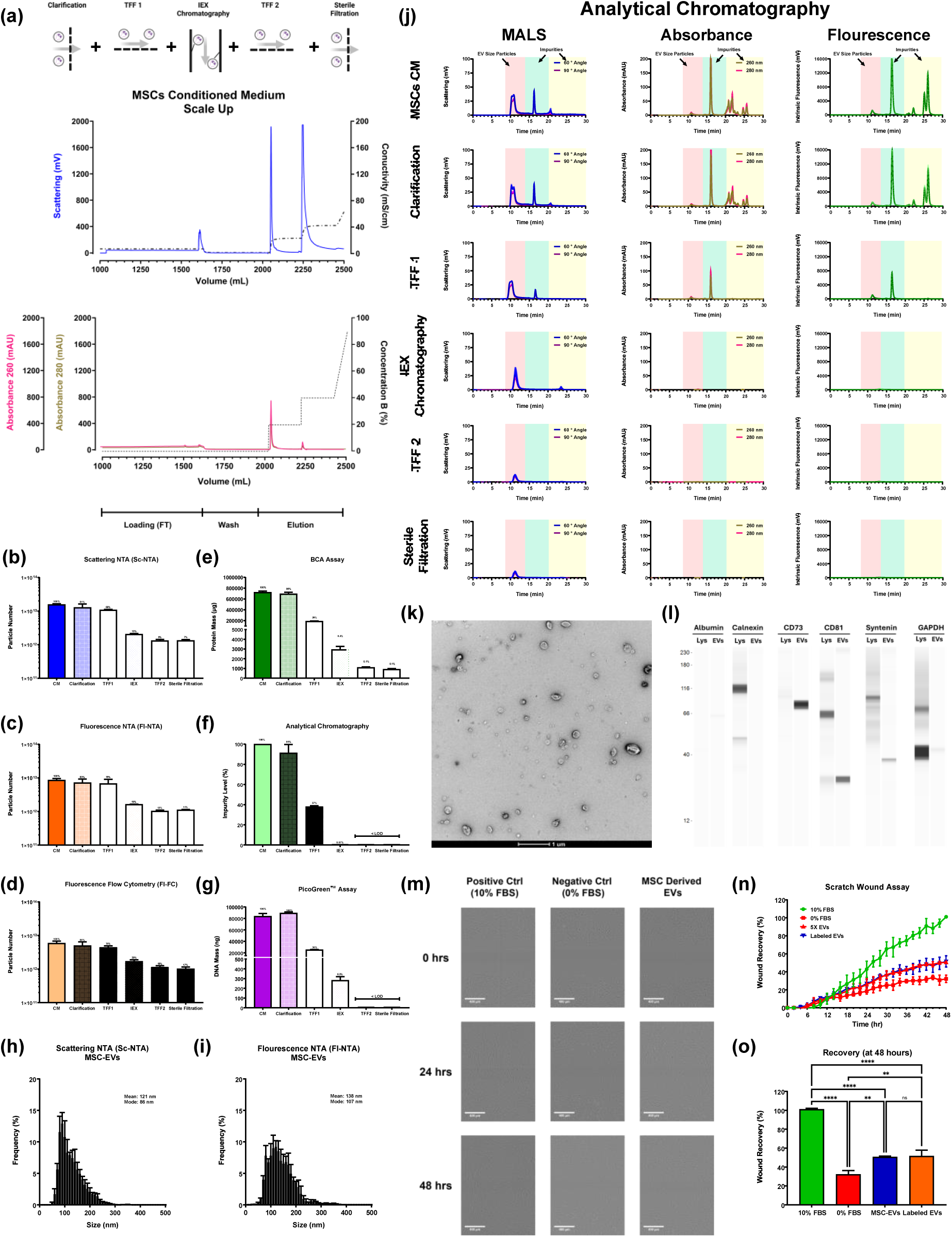
**Downstream Processing (DSP) Scalability;** Scaled DSP for MSC derived EVs and orthogonal analytical assessment of particle recovery and impurity removal. **(a)** Schematic representation of the scaled DSP workflow comprising clarification, tangential flow filtration (TFF), ion exchange (IEX) chromatography, post-IEX buffer exchange via TFF, and sterile filtration. Particle recoveries for each unit operation were determined by **(b)** -NTA, **(c)** Fl-NTA, and **(d)** Fl-FC. Protein removal throughout the entire DSP was quantified using **(e)** BCA assay, while impurity reduction was evaluated by **(f)** analytical chromatography through analysis of impurity-associated peak area under the curve (AUC). DNA removal was assessed via **(g)** PicoGreen™ assay. Representative chromatograms of analytical chromatography for DSP steps are shown in **(j)**, with EV-size peaks (9–14 min) highlighted in red, impurity peaks (14–20 min) in green, and late-eluting impurities (>20 min) in yellow. Final MSC-EVs preparation characterization included **(h)** scattering and **(i)** fluorescence NTA size distribution profiles, **(k)** transmission electron microscopy (TEM) imaging showing characteristic EV cup-shaped morphology, scale bar: 1 µm, and **(l)** Simple Western confirming EV marker expression (CD81, CD73, Syntenin) and absence of negative markers (calnexin, albumin). **(m–o)** Fibroblast scratch migration assay demonstrating retained bioactivity of purified MSC-EVs, quantified by cell density recovery within the wound area over 48 hour, supported by qualitative representative images, scale bar: Unless otherwise specified, Multicomparison statistical analysis was performed using ordinary one-way ANOVA with Tukey’s multiple comparison test (*p<0.05, **p<0.01, ***p<0.001, ****p < 0.0001; ns not significant).

Following 6X diafiltration, protein content was reduced to 26% of the MSC-CM feed, supported by analytical chromatography and AUC analysis, which indicated a reduction in impurity levels to 37% **(Fig 9-[e, f and j] and Sup Fig 18-g)**. Impurity peaks after 20 minutes (highlighted in yellow) were completely removed, and the 14–20 minute (highlighted in green) impurity region was substantially reduced **(Fig 9-j)**. Particle recoveries were 68% (Sc-NTA), 78% (Fl-NTA), and 74% (Fl-FC) **(Fig 9-[b-d] and Sup Fig 18-c)**. DNA content was reduced to 30% of the feed mass following benzonase digestion and diafiltration **(Fig 9-g and Sup Fig 18-h)**.

For IEX chromatography, only the EV-containing eluate (eluate 2) was analyzed **(Fig 9-a)**. Particle recoveries were 13% (Sc-NTA), 19% (Fl-NTA), and ∼30% (Fl-FC) **(Fig 9-[b-d] and Sup Fig 18-d)**. Analytical chromatography showed complete removal of impurity peak (14–20 min, highlighted in green), while BCA and AUC analyses indicated 99% protein removal **(Fig 9-[e, f and j] and Sup Fig 18-g)**. DNA concentration was reduced to 0.3% of the original feed, as determined by PicoGreen™ assay **(Fig 9-g and Sup Fig 18-h)**.

Subsequent processing steps included buffer exchange (second TFF) and sterile filtration. Final total particle recoveries of the second TFF and sterile filtration were 9% and 7% (Sc-NTA), 12% and 11% (Fl-NTA), and 18% and 17% (Fl-FC), respectively **(Fig 9-[b-d] and Sup Fig 18-{e and f})**. BCA and AUC analyses are presented, and both DNA and impurity signals were below the lower LOD at this stage in **(Fig 9-[e-g] and Sup Fig 18-[g and h])**.

Final EV characterization confirmed the presence of typical EV biophysical and biochemical characteristics. The final sterile MSC-EV samples showed expected size distribution on SC-NTA and Fl-NTA and showing mean size of 121 nm and mode of 86 nm in scattering while fluorescence showed 138 nm for mean and 107 nm for mode **(Fig 9-[h and i]**. The size distribution of the samples across the entire DSP process can be found in **(Sup Fig 18-i).** TEM imaging revealed characteristic cup-shaped morphology **(Fig 9-k)**. Simple Western analysis further validated EV identity, showing strong expression of CD81 and CD73, a faint band for Syntenin, and absence of calnexin and albumin, supporting EV purity **(Fig 9-l)**. A summary of particle, protein, and DNA concentrations across all DSP steps is provided in **(Sup Fig 18-[a-h])**. Final EV preparations yielded 2.55 × 10^10^ particles/mL (Sc-NTA), 2.23 × 10^10^particles/mL (Fl-NTA), and 2.03 × 10^10^particles/mL (Fl-FC), with a total protein concentration of 16 µg/mL (BCA) and DNA content below the lower LOD of PicoGreen™ assay **(Sup Fig 18-[a-h])**.

MSC-EVs have been shown to increase fibroblast and endothelial cell migration (Costa *et al*., 2023; Ferreira *et al*., 2017; Kim *et al*., 2023; Manzoor *et al*., 2024). Therefore, we verified that our MSC-EVs preparation retained bioactivity in a fibroblast scratch assay. Scratch wound assays are normally analyzed based on wound area closure. However, we developed a new analysis method to report recovery based on the number of cells that migrated in the wound area compared to the original number of cells before scratch was introduced. We developed a custom Python script to quantify cell density in the scratched area compared to the unscratched area in the same field of view. Fibroblast scratches treated with the MSC-EV preparation showed significantly higher migration into the scratched area compared to the vehicle control over the course of 48 hours **(Fig 9-[m-o])**. Qualitative evaluation of images confirmed treatment effectiveness and our quantitative results **(Fig 9-[n and o])**. Furthermore, EV preparation labeling qualitatively verified EV uptake by fibroblasts whereas the dye only control showed no signal **(Sup Fig 18-[k and l])**. These results suggest our robust DSP and high purity MSC-EV preparations retain bioactivity.

## Discussion

The complex and heterogeneous nature of EVs are the major players in the historical analytical challenges in the field. Therefore, well-studied, well-characterized, reproducible, and standardized analytical assays are critical for the development of end to end, reproducible and scalable EV biomanufacturing workflows, from research and development to clinical translation and commercialization. Analytical assays optimization drives: a) accurate quantification and characterization, b) reproducibility and consistency, c) standardization across the field, d) process optimization, e) acceleration to therapeutic development, f) support of regulatory and quality requirements. Schematics were created in Biorender.com.

In this study, we established three orthogonal analytical techniques; NTA (Scattering and fluorescence), FC (fluorescence), and analytical chromatography. We utilized liposomes as a suitable model or reference for assay development and optimization. Liposomes possess unique properties that make them strong surroges for EVs, such as a similar refractive index, similar size distribution, vesicular nature and the absence of non-vesicular particles. However, it is important to acknowledge the caveat that liposomes tend to spontaneously fuse to form multi-lamellar structures. Multi-lamellar liposomes have a different refractive index compared to single phospholipid bilayer EVs and this refractive index mismatch can impact their usability as a reference material in scattering-based detection (Welsh *et al*., 2020b).

After optimization of the operating parameters of NTA as well as fluorescence labeling optimization using CellMask Orange™ (CMO) dye, three single particle analysis techniques delivered comparable particle concentration measurements. When standard rEVs were tested, similar results were obtained among Sc-NTA, Fl-NTA and Fl-FC. Although Sc-NTA returned a higher concentration potentially due to presence of impurities and lower specificity in scattering mode. It is also important to note that the measurements from each device represent the concentrations of particles above the lower LOD of each respective instrument and technique.

Analytical chromatography coupled with analytical SEC and equipped with in-line MALS, UV and Fluorescence detectors served as a critical component in process optimization. As single particle analysis techniques tend to become less reliable in crude samples, analytical chromatography provides comprehensive results regarding the presence of different size molecules including EVs and impurities.

ISEV Rigor and Standardization (R&S) subcommittee, aiming to improve reproducibility of EV research, recently established a task force on CCM-EV focusing on cell culture parameters and their impact on EVs (Shekari *et al*., 2023). From the proposed cell culture parameter checklist, we have reported the details to the best of our capabilities in this study, including details of producing cells, EV production medium, culturing conditions, and collection of MSCs CM. However, we did not perform any microbial contaminant evaluation of CM. In summary, our study was performed in 3D using a 10L single use bioreactor and microcarriers. It has been suggested that 3D cultures produce higher EVyields compared to 2D systems due to higher cell density and higher EV production rate per cell per volume *(Costa et al., 2023;* *De Almeida Fuzeta et al*., 2020; Shekari *et al*., 2023). We also used PLT as a complex biological supplement in the growth and expansion phase of the cell culture as opposed to FBS, to avoid animal components and develop the upstream process in a Xeno-free medium (Shekari *et al*., 2023; Staubach *et al*., 2021). The medium was switched to PLT free medium in EV production and collection phase, to ensure that the collected MSCs CM did not contain EVs derived from PLT.

A new type of IEX chromatography column, called Sartobind Convec® D was the main component in the DSP process. Sartobind Convec® D is a membrane absorber specifically designed for purification of lentiviruses with mild salt elution strategy. The RT-MALS detector connected to the chromatography instrument played a crucial role in this study. Without the RT-MALS detector, only one peak would have been detected using UV detector (IEX eluate 1 – 200 mM NaCl) which we identified to be non-vesicular particles. Additionally, adjusting the elution strategy from linear elution to stepwise elution led to the elution of two distinct populations. Both IEX eluates exhibited a comparable and strong RT-MALS signal while IEX eluate 1 demonstrated a pronounced absorbance signal and IEX eluate 2 demonstrated little absorbance.

To ensure that the two populations are indeed cell-secreted, we included a critical control experiment, medium alone control. Loading clarified medium alone control on the IEX column demonstrated RT-MALS and UV signals from IEX eluate 1 with no signal on eluate 2. Therefore, we included a TFF step with 6X diafiltration prior to the IEX chromatography to remove medium-associated components. Analytical chromatography revealed that the implementation of 6X diafiltration unit operation prior to IEX chromatography reduced the signal on all 3 detectors below the lower LOD, which was further confirmed by BCA analysis showing below the lower LOD protein content in 6X diafiltered medium alone. When the 6X diafiltered sample was loaded on IEX chromatography, no signal was detected on RT-MALS, with intrinsic fluorescence and UV detectors further validating successful removal of medium-associated components.

The sequence of clarification, TFF and IEX chromatography was then performed in triplicate demonstrating the reproducibility of the process, in which two IEX eluates were collected, one with strong RT-MALS and absorbance signal and the other with comparable RT-MALS but minimum absorbance signal, also ensuring both IEX eluates to be cell-secreted. These results were in agreement with off-line analysis via Sc-NTA and analytical chromatographic plots from RT-MALS signal with both detecting comparably. A high absorbance and intrinsic signal on analytical chromatography also validated the observed real time UV detector while IEX chromatography was conducted. The analytical chromatography results on all the samples across unit operations showed complete removal of all the impurity fractions detected in MSCs CM harvest suggesting successful purification and enrichment of EVs.

In addition to validating reproducibility, we evaluated the scalability of the established process, while expanding the process by adding a second TFF step for buffer exchange and sterile filtration at the end of the process. While the scalability process was conducted, replicating the small-scale process, it is important to note that, if the target final product is EVs, the first TFF step is not necessary. The exclusion of the first / step can increase the total recovery of the process as it was measured by Sc-NTA, Fl-NTA and Fl-FC. The observed concentration measurements again showed the importance of well-optimized and reliable analytical tools. Sc-NTA failed to distinguish between NVEPs and EVs, while fluorescence-based detection demonstrated increased specificity towards EVs. However, it still suffered from non-specific interactions between the dye and impurities and leftover EVs that co-eluted in eluate 1 as well as in the crude samples, suggesting concentration measurements are not completely reliable in crude samples.

Other groups have also utilized IEX chromatography for purification and enrichment of EVs from different cell lines such as HEK293 *(Heath et al., 2018; Koch et al*., 2023) and MSCs (Malvicini *et al*., 2023). The most comparable study was done by Koch *et al*., using IEX chromatography also equipped with RT-MALS detector (Koch *et al*., 2023). However, the study was done using linear gradient strategy and not stepwise. Similarly, the authors used a wide range of analytical methods to study the collected fractions such as NTA, ELISA, BCA, Western Blot, AF4, TEM and Exoview. Interestingly, the authors also observed two partially resolved MALS peaks while only one expressed an absorbance signal.

New MISEV Guidelines have highlighted the use of accurate terminologies and emphasized on the separation and analytical challenges of non-vesicular extracellular particles and vesicular particles (Welsh *et al*., 2024). Zhang *et al*.,in 2018 identified a new subset of non-vesicular extracellular particles termed exomeres which were separated by asymmetric-flow field flow fractionation (AF4) (Zhang *et al*., 2018). The online QELS monitor measured the hydrodynamic diameter of the exomeres fraction to be around 47nm while NTA failed to resolve exomere fractions from small and large exosome (term used in the original article) based on size. TEM images however, identified a population smaller than 50 nm (∼35 nm) which lacked an external membrane structure and was found in 20 different cell lines (Zhang *et al*., 2018). AF4 was further used in multiple studies to separate EVs from different sources such as human serum (Kim *et al*., 2020) and human plasma (Wu *et al*., 2020). AF4 has been also coupled with other techniques like capillary electrophoresis for separation of EVs (Gao *et al*., 2022) and density cushion ultracentrifugation for EVs in human blood (Hu *et al*., 2024).

Recently, ultracentrifugation also has been used to enrich exomeres. Zhang *et al*., enriched small EVs (sEVs) by centrifugation at 167K two times, and then subjected the supernatant to the same g force for 16 hours to enrich exomeres (Zhang *et al*., 2019). While NTA data showed an overlap in the size distribution between sEVs and exomeres, sEVs exhibited a larger size distribution. The group proceeded even further in a different study by subjecting the supernatant of the exomere step to ultracentrifugation at 367,000g for 16 hours and the pellet was termed supermere (Jeppesen *et al*., 2023; Zhang *et al*., 2021). In 2022, Tosar *et al*., in a commentary article, raised interesting questions on whether exomeres and supermeres are homogenous extracellular nanoparticles or a complex mixture consisting of compositionally distinct extracellular nanoparticles that tend to co-isolate due to similar sedimentation coefficients and size by ultracentrifugation (Tosar *et al*., 2022). Authors also addressed the fact that Galectin-3-binding protein (LGALS3BP) which was originally described as a potential universal marker of exomeres, is in fact a protein of the extracellular matrix that self-assembles into particles (ring-like decamers) of 30-40 nm. The authors concluded a wide range of extracellular complexes such as spliceosome, ribosomes and other abundant cellular complexes co-isolate with exomeres and supermeres (Tosar *et al*., 2022). Similarly, our eluate 1 (NVEPs) is a mixture of extracellular complexes and not one single distinct but heterogeneous subpopulation.

Additionally, Ghosal *et al*., recently showed enrichment of 3 different populations of extracellular particles and termed them as 14K-lEV, 100K-sEV and 167K-NVEP using sequential centrifugation steps (Ghosal *et al*., 2025). They found that NVEPs display a distinct biochemical and molecular signature, with elevated protein-to-lipid ratio, using biochemical assays and Raman spectroscopy. It is critical to note that in order to study the impact of the techniques used for EVs vs NVEPs separation, the same feed needs to be tested side by side and the cell culturing parameters should remain the same in CM production phase.

It is important to acknowledge that the level of impurity removal achieved in this study may not be necessary for all EV-based therapeutic products. The degree of purification required depends largely on the intended therapeutic indication and how the final EV product is defined. For example, in the first-in-human clinical study by Warnecke *et al*., investigating human stromal cell derived extracellular vesicles, the final product was described as an *EV-enriched secretome fraction*, a preparation that contains EVs alongside other soluble, cell-secreted factors (Warnecke *et al*., 2020). This terminology reflects a broader compositional profile that may retain biological activity despite the presence of non-vesicular components.

For such less purified EV formulations, the critical consideration is not necessarily complete impurity removal, but rather the clear definition of product-specific quality attributes. Establishing these attributes is essential for ensuring batch-to-batch consistency, product reproducibility, and alignment with regulatory expectations. Therefore, while high-purity EV preparations may be required in certain contexts, other applications may prioritize functional performance and compositional consistency over excessive purification.

Finally, we verified our EV preparations retained bioactivity in a fibroblast scratch wound assay. Adipose MSC-EVs produced in a stirred tank bioreactor have been shown to improve HUVEC migration in a scratch wound assay after 20 hours (Costa *et al*., 2023). Our results agree with the findings that adipose MSC-EVs significantly improve fibroblast migration after 24 hours (Ferreira *et al*., 2017), and Wharton’s jelly MSC-EVs improve fibroblast migration in vitro after 24 hours as well as in a rat full thickness skin wound (Kim *et al*., 2023). Our EV preparation dose (8.46 × 10^4^particles/seeded cell) was relatively high compared to doses seen in the literature. Additionally, we ensured consistent scratches with a qualified tool and performed live imaging to quantify wound closure over 48 hours. Our custom Python script enabled label free quantification of cell density in the scratched area normalized to the unscratched area in the same field of view, potentially increasing migration quantification accuracy and dynamic range compared to software calculating percent closure based on the leading edge of migrating cells. This robust implementation of the fibroblast scratch wound assay serves as a starting point for evaluating EV bioactivity in response to USP and DSP changes.

The primary objective of this study was to establish reliable analytical workflows and DSP strategies for extracellular vesicles (EVs) derived from MSCs (**Fig 10**). While the upstream process was not optimized in the current work, the robust analytical and DSP framework developed here provides a strong foundation for future upstream process development. Upcoming studies will focus on optimizing critical upstream parameters, including medium composition (e.g., comparison of chemically defined versus serum- or platelet lysate-supplemented medium), cell seeding density, passage number, and the timeline of cell expansion and EV production.

**Figure 10.**
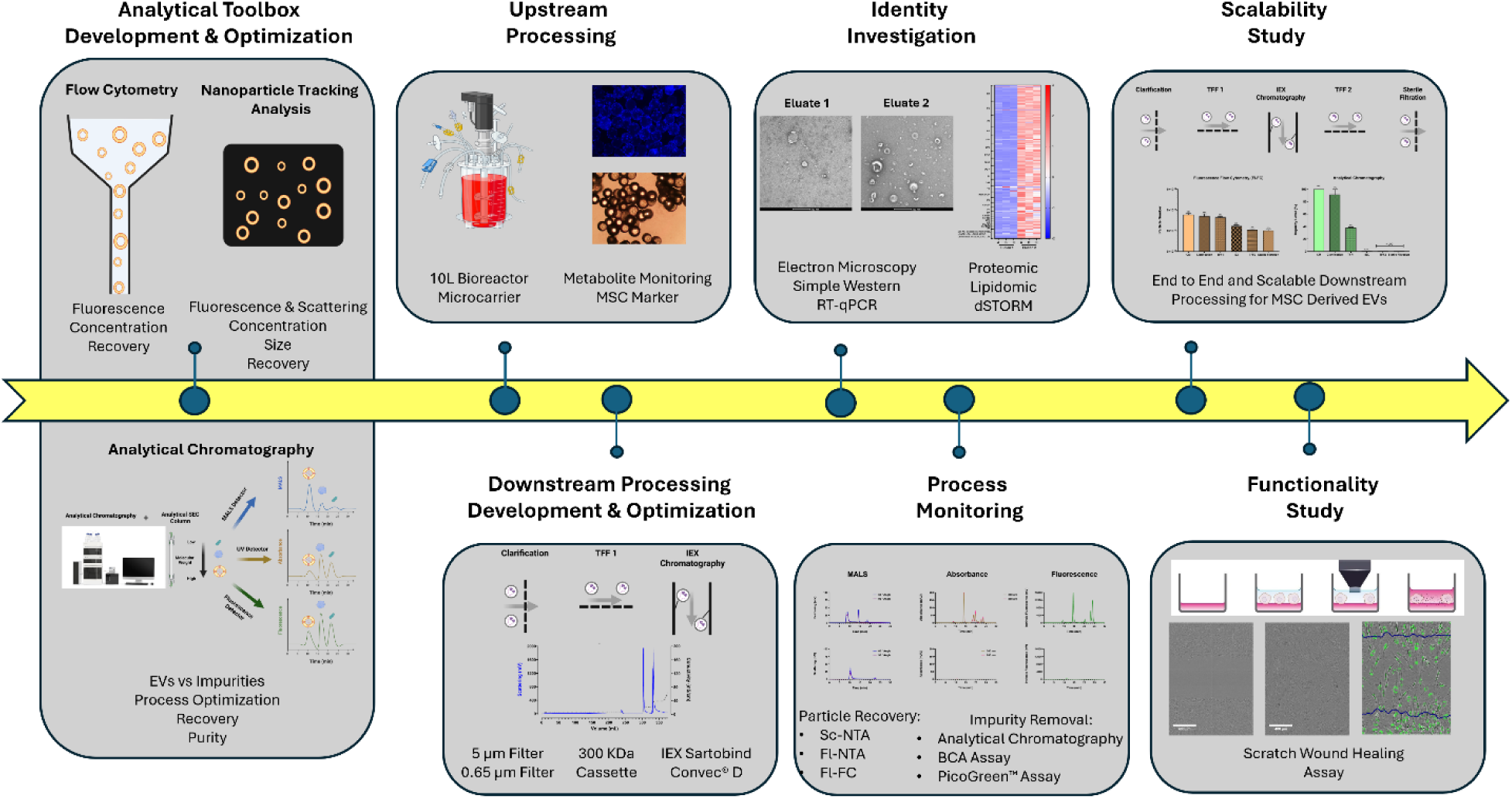
**Study Workflow;** Study workflow showing the establishment and optimization of the analytical toolbox containing Nanoparticle Tracking Analysis, Fluorescence-based Flow Cytometry and Analytical Chromatography. The Analytical toolbox was then utilized to establish a robust downstream process for Mesenchymal Stromal Cell derived Extracellular Vesicles in a single use bioreactor and microcarriers. The Downstream process using Ion Exchange Chromatography resulted in enrichment of Extracellular vesicles versus Non-Vesicular Extracellular Particles in two different eluates. The Downstream process was then shown to be reproducible. The identity of these two eluates was then comprehensively studied using orthogonal techniques, such as Electron Microscopy, Simple Western, Proteomics and Lipidomics. The downstream process was then excessively monitored for particle recovery and purity level using the established analytical toolbox. The established downstream process was then successfully shown to be scalable. Lastly, the final Extracellular Vesicles preparation, maintained bioactivity in a scratch wound assay. Schematics were created in Biorender.com.

In addition to upstream optimization, future efforts will aim to evaluate the scalability and applicability of the established DSP across different MSC sources such as umbilical cord and adipose derived; but also, other cell types commonly used in biomanufacturing, such as HEK293, CHO, and immortalized MSC lines. These studies will be essential for broadening the utility of the platform and supporting the development of scalable, standardized workflows for diverse EV-based therapeutic applications.

While the Sartobind Convec® D membrane column represents a mild IEX chromatography platform, enabling the elution of target vesicles at relatively low salt concentrations, other chromatographic approaches may offer additional resolution. For instance, stronger IEX platforms such as monolithic columns provide a stronger bond between target molecules and the column, which could be employed to further fractionate purified EV populations. This enhanced separation capability could enable the identification and characterization of EV subpopulations with distinct biophysical or biochemical properties, thereby offering new insights into EV heterogeneity and its potential relationship to therapeutic function.

## Author Contributions

M.D. contributed to analytical optimization, upstream processing and downstream processing with designing and executing experiments as well as analyzing the data and writing the manuscript. M.C. contributed to the analytical optimization with designing and executing experiments and analyzed the data. N.T. M.M and D.J. performed the downstream processing optimization with designing and executing experiments and analyzed the data. A.L. contributed to the functionality assay and wrote the manuscript. M.M. contributed to the design of experiments and optimization of the functionality assay. Y.K. contributed to scalability study and the characterization using Simple Western and PCR and wrote the manuscript. Y.W. and Y.Z. conducted off-line analytics in the scalability study. R.C, Y.L., B.L., and C.T. contributed to the analytical optimization. J.S., P.K., and P.M contributed to the development of custom design algorithms for fluorescence-based flow cytometry and scratch wound healing assay. N.H., D.S, and E.B., contributed to upstream processing development. Z.Z., M.I. and P.M. contributed to the development of downstream processing. Z.Z. also contributed to data analysis of proteomics. T.G., J.C. and E.K. contributed to the analytical optimization and multi-laboratory/instrument studies.

## Acknowledgements

The authors thank Dr. Chad Calloway for electron microscopy, which was performed in the Electron Microscopy Resource (RRID: SCR_012366) in the Center for Advanced Research Technologies at the University of Rochester Medical Center. The authors thank Ricardo Bastos and Chris Gabriel who performed ONI super-resolution nanoparticle imaging experiments. The authors thank Oscar Reif, Michael Olszowy, David Pollard, Sunandan Saha and Mark Szczypka for their support and consulting throughout this whole study. The authors thank Richard Jones who performed proteomic analysis. The authors thank Dr. Jeff Chu who performed lipidomic analysis. The authors thank Eric Bushong who performed the Cryo-EM and size distribution analysis of liposomes.

