## Supplementary Material for "Orthogonal and Robust Analytics Enable Reproducible and Scalable Manufacturing of High Purity Extracellular Vesicles Derived from Mesenchymal Stromal Cells"

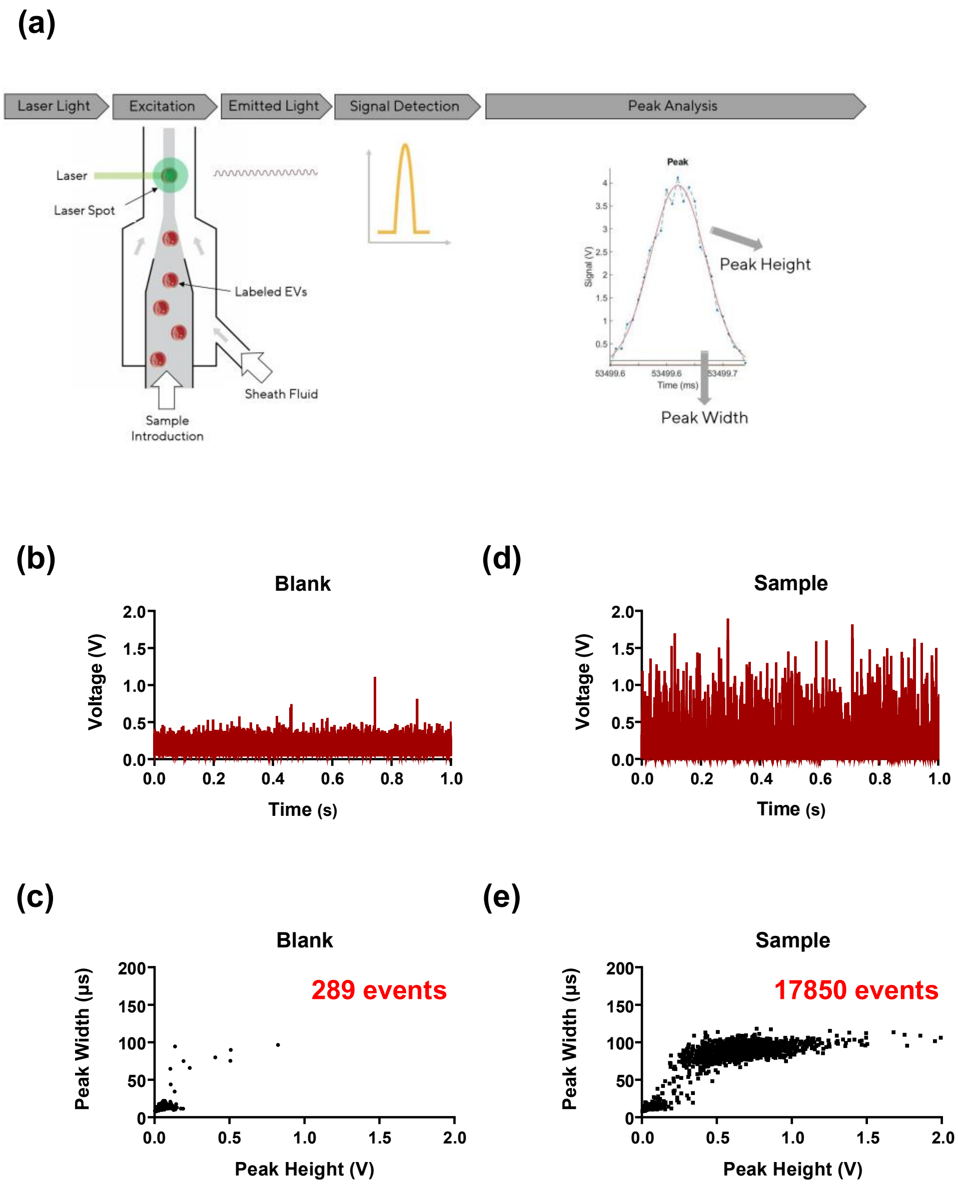


**Supplementary Figure 1. Analytical Optimization;** Schematic and representative data outputs of the detailed peak analysis method for characterization of fluorescently labeled extracellular vesicles (EVs) using fluorescence-based flow cytometry (FI-FC). **(a)** Schematic overview of the detection workflow. Labeled EVs are hydrodynamically focused and pass through laser illuminated spot where excitation and emission occur. The emitted signal is detected as raw voltage values, which is then fit by a gaussian curve and subsequently analyzed to extract peak height and peak width parameters using an in-house algorithm.**(b, c)** Representative 1 second raw signal histogram **(b)** and corresponding 2D peak height vs. peak plot **(c)** for a blank (buffer alone) control, showing 289 peaks.**(d, e)** Representative 1 second raw signal histogram **(d)** 2D peak height vs. peak width plot **(e)** for a sample, showing 17850 events.

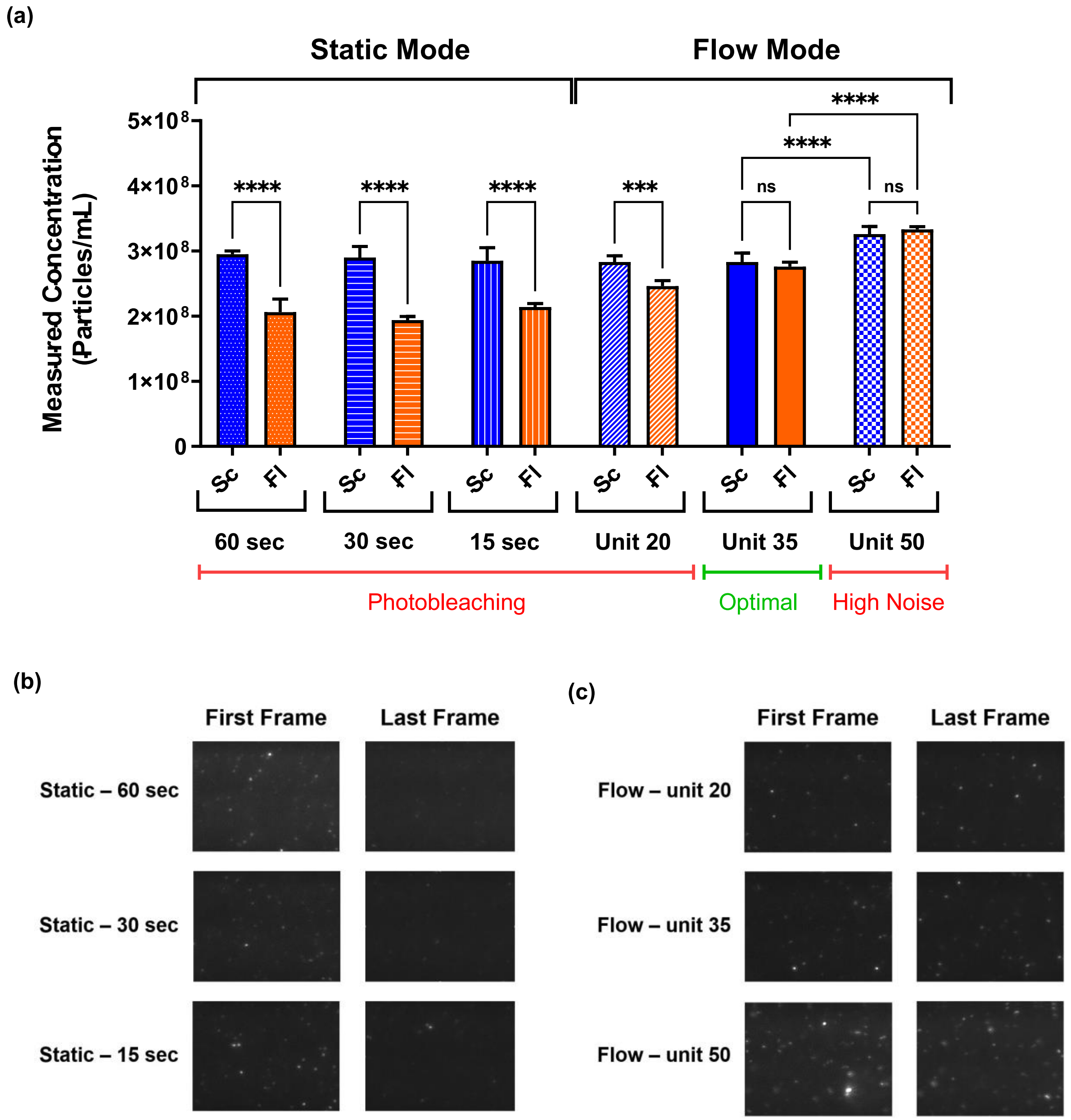


**Supplementary Figure 2. Analytical Optimization;** Evaluation and optimization of NTA acquisition settings to minimize photobleaching and ensure accurate particle quantification in fluorescence mode.

**(a)** Comparison of liposome concentrations measured in scattering (Sc) and fluorescence (Fl) modes under static and flow acquisition settings. In static mode, fluorescence-based measurements significantly underestimated the concentrations relative to scattering mode across all acquisition times (60 s, 30 s, 15 s) due to the photobleaching effect. Transitioning to flow mode reduced photobleaching and improved agreement between Sc and Fl measurements. At flow unit 20, discrepancies between modes persisted, while flow unit 50 yielded artificially high concentrations due to excessive noise and tracking artifacts. Flow unit 35 produced the most consistent results, with no significant difference between Sc and Fl modes, indicating no photobleaching and reliable enumeration. Multicomparison statistical analysis was performed using ordinary one-way ANOVA with Tukey's multiple comparison test (*$p<0.05$, **$p<0.01$, ***$p<0.001$, ****$p < 0.0001$; ns = not significant). All data are presented as mean ± standard deviation with n= 5 independent recorded videos. **(b, c)** Representative NTA video frames from static **(b)** and flow **(c)** modes. In static mode, fluorescence intensity visibly diminished between the first and last frame at all acquisition durations, confirming photobleaching. In contrast, flow mode preserved fluorescence intensity across frames.

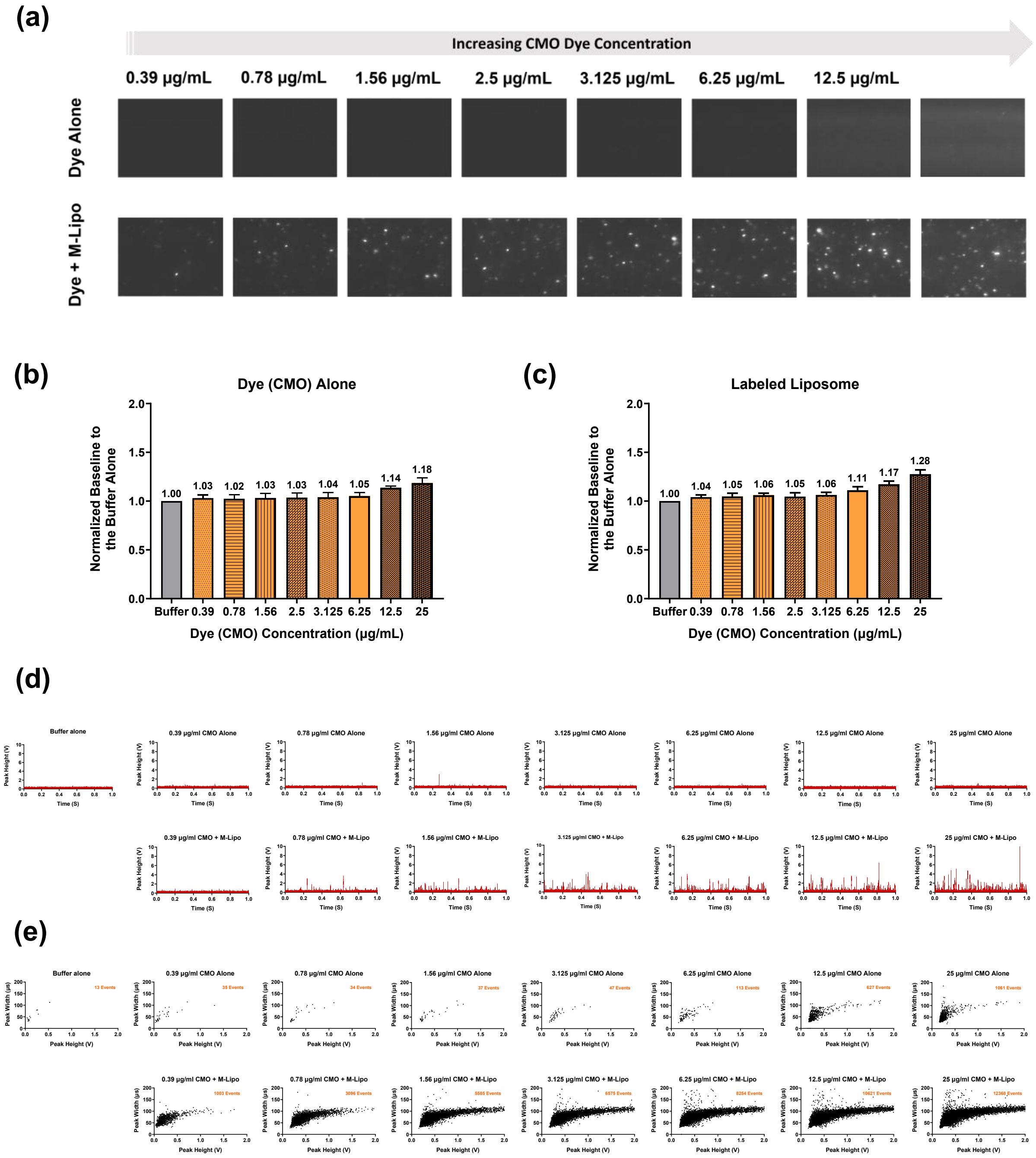


**Supplementary Figure 3. Analytical Optimization;** Optimization of a no-wash assay for fluorescence labeling of liposomes using CellMask Orange (CMO).

**(a)** Representative NTA video frames of CMO dye alone (top row) and labeled medium-sized liposomes (M-Lipo; bottom row) while systematically increasing dye concentrations (0.39–25 µg/mL). At concentrations between 0.78–6.25 µg/mL, bright fluorescence signals were observed in the presence of liposomes, with minimal background in dye alone controls. Higher concentrations (12.5 and 25 µg/mL) resulted in increased background fluorescence in dye alone samples. **(b–c)** Flow cytometry baseline analysis for **(b)** dye alone control and **(c)** labeled liposomes. At high dye concentrations (12.5 and 25 μg/mL), both dye alone and labeled samples showed elevated baseline fluorescence, indicating increased background noise. **(d)** Representative one second raw signal histogram from FI-FC for dye alone controls (top row) and labeled liposomes (bottom row). **(e)** 2D plots of peak height versus peak width of the detected events. Dye alone controls at high concentrations show number of events above the lower LOD, consistent with dye aggregation or micelle formation, while labeled samples demonstrate effective part cle labeling at intermediate concentrations.

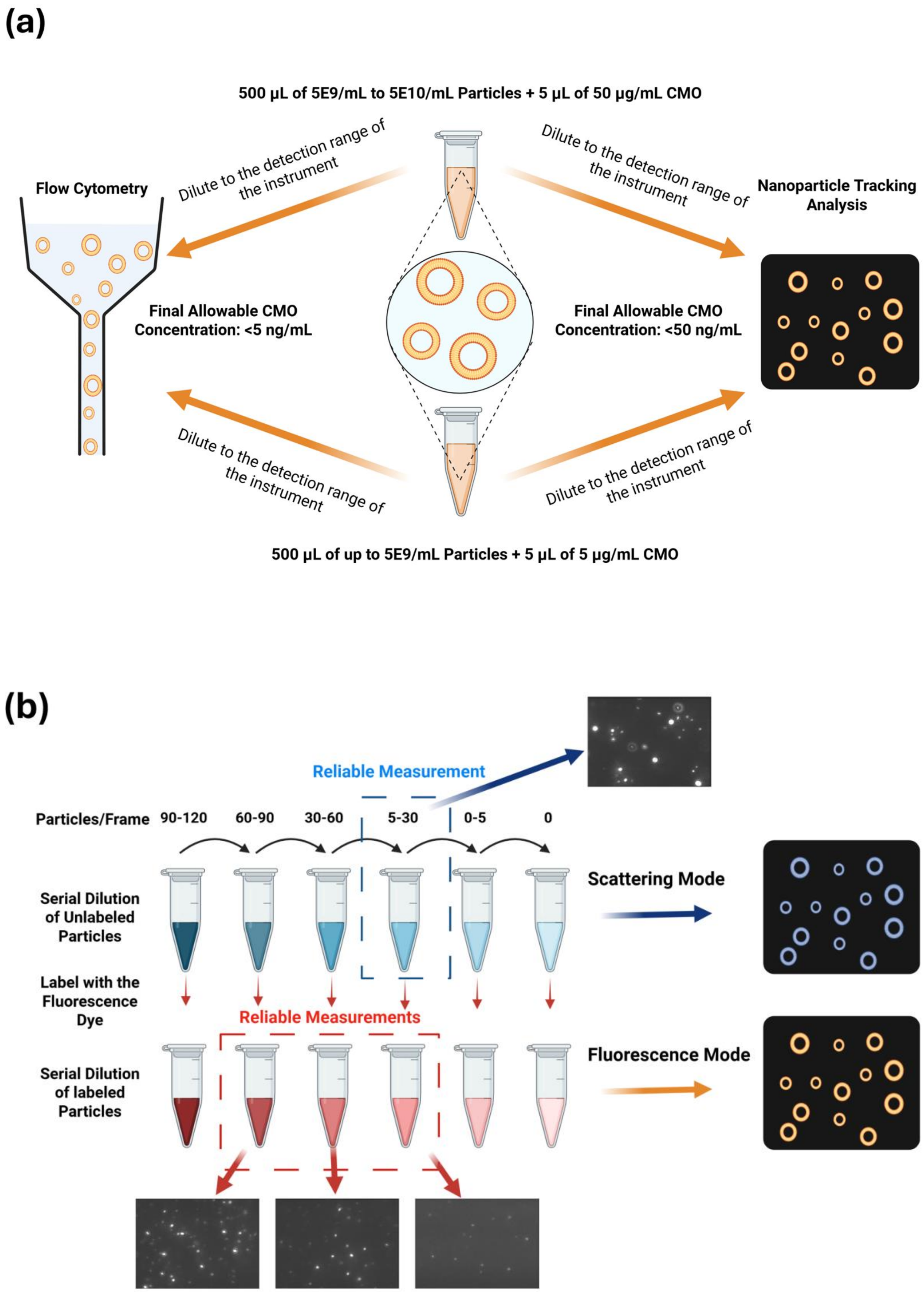
(a)
500 µL of 5E9/mL to 5E10/mL Particles + 5 µL of 50 µg/mL CMO
Dilute to the detection range of the instrument
Dilute to the detection range of the instrument
Flow Cytometry
Nanoparticle Tracking Analysis
Final Allowable CMO Concentration: <5 ng/mL
Final Allowable CMO Concentration: <50 ng/mL
Dilute to the detection range of the instrument
Dilute to the detection range of the instrument
500 µL of up to 5E9/mL Particles + 5 µL of 5 µg/mL CMO
(b)
Reliable Measurement
Particles/Frame 90-120 60-90 30-60 5-30 0-5 0
Serial Dilution of Unlabeled Particles
Scattering Mode
Label with the Fluorescence Dye
Reliable Measurements
Serial Dilution of labeled Particles
Fluorescence Mode

**Supplementary Figure 4. Analytical Optimization;** Schematic workflow of a no-wash labeling assay and quantification using flow cytometry and nanoparticle tracking analysis.

**(a)** Illustration of the optimized labeling protocol using CMO dye. Particles (up to 5E9/mL) are labeled with CMO at a final concentration of 5 µg/mL (500 µL of sample + 5 µL of 5 µg/mL CMO). Particles (up to 5E10/mL) are labeled with CMO at a final concentration of 5 µg/mL (500 µL of sample + 5 µL of 50 µg/mL CMO). Following incubation and prior to measurement, samples are diluted to fall within the detection range of each single particle analytical technology. Maximum allowable CMO concentrations for analysis are <50 ng/mL for fluorescence mode nanoparticle tracking analysis (FI-NTA) and <5 ng/mL for flow cytometry (FC), based on background signal thresholds determined from dye only controls. **(b)** NTA Measurement Strategy for reliable quantification with minimal masking effect. Unlabeled particles are serially diluted and the optimal particle per frame is 5-30 in scattering mode, while fluorescence mode has a wider optimal range, 5–60 particles/frame. Schematics were created in Biorender.com.

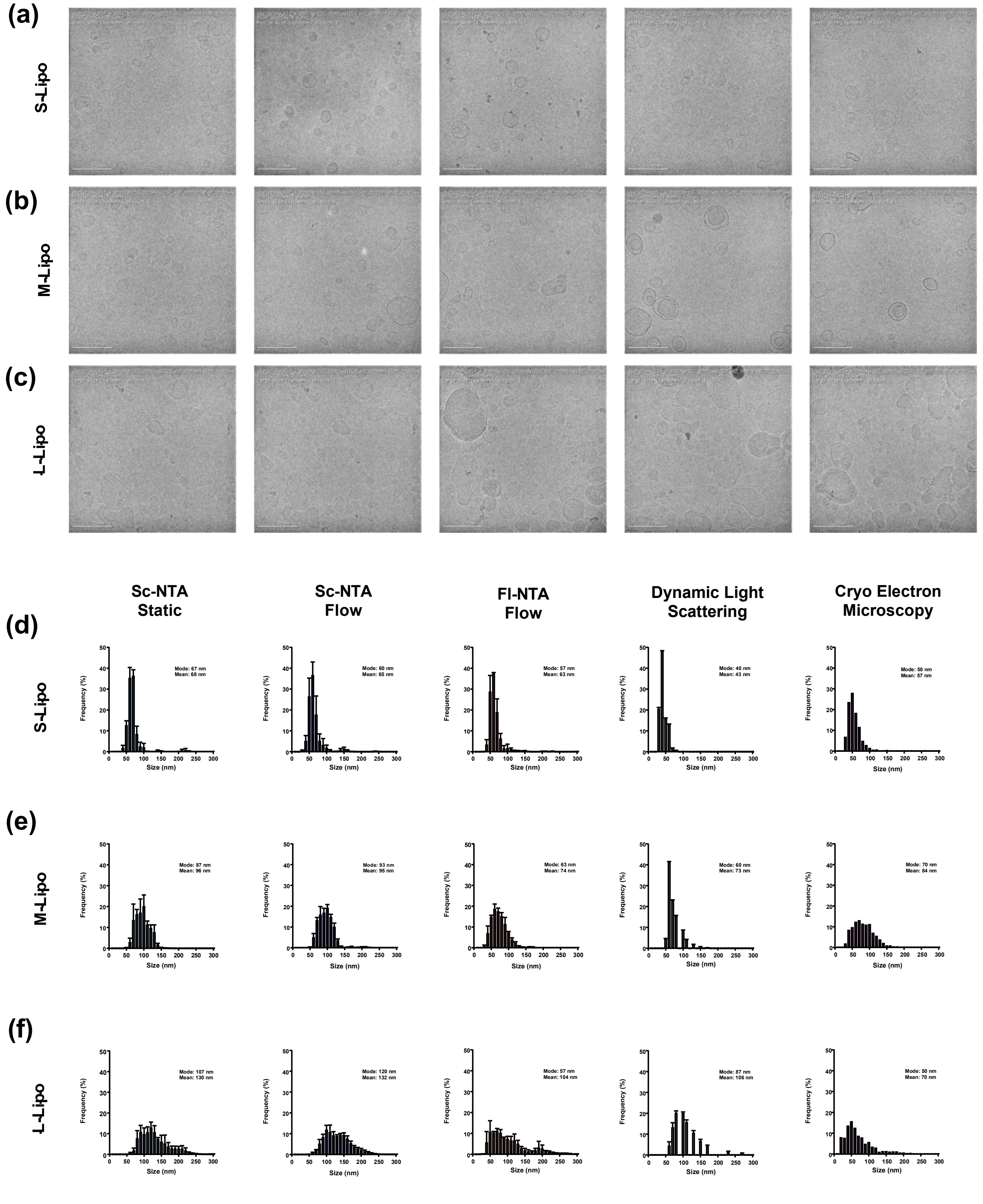


Supplementary Figure 5. Analytical Optimization; Comparative size distribution analysis of liposome samples using orthogonal techniques.

**(a–c)** Representative cryo-electron microscopy (cryo-EM) images of small (S-Lipo), medium (M-Lipo), and large (L-Lipo) liposome samples. Scale bars: 200 nm. Over 1,000 individual particles were analyzed for each liposome sample to generate size distribution profiles. **(d–f)** Size distribution histograms for **(d)** S-Lipo, **(e)** M-Lipo, and **(f)** L-Lipo obtained using scattering-mode nanoparticle tracking analysis in static and flow modes (Sc-NTA Static, Sc-NTA Flow), fluorescence mode NTA in flow mode (Fl-NTA Flow), dynamic light scattering (DLS), and cryo-EM. Among the tested techniques, Fl-NTA Flow showed similar size distribution profile as cryo-EM, accurately capturing both the peak diameter and distribution width across all liposome sizes. All data are presented as mean ± standard deviation with n=3 independent measurements.

### Size Distribution Analysis by Cryo-Electron Microscopy:

The 28,000X magnification images were used in the size distribution analysis. These images have a field of view of 2.1 µm x 2.1 µm and a pixel size of 0.512 nm. The images in the experiment were first pre-screened for quality and then randomly ordered for the analysis. A minimum of 1000 particles, or all relevant particles available, were manually traced in the image set by a senior analyst. Particle selection was based on the following criteria: (1) particles were not selected if they were over the carbon film or touching the edge of the carbon film, and (2) particle boundaries had to be clearly defined. The following particle size metrics were computed from the particle contours:

Area; the area enclosed by the contour. Perimeter; the length of the contour. Maximum and Minimum Feret Diameters; the largest and smallest projected diameter of the particle contour. Circularity; a derived metric for the roundness of the contour. A perfect circle has a circularity of 1 and a square has a circularity of 0.76.

$$Circularity = 4\pi \times \frac{\text{Area}}{(Perimeter)^2}$$

Area Equivalent Diameter (AED); the diameter of a circle with equivalent area to the tracing of the contour.

$$AED = 2 \times \sqrt{\frac{Area}{\pi}}$$

In the analysis of liposome samples, 11 images were used, and 1079 particles were analyzed for small liposomes, 19 images and 1020 particles for medium liposomes, and 8 images and 1146 for large liposomes. [Cryo-Electron microscopy was performed by Nanoimaging Services]

- **Size Distribution Plotting:**

All size distribution results were plotted as frequency vs size for comparison between techniques. The frequency was done for 100 bins with bin size of 10 nm from 0 - 1000 nm.

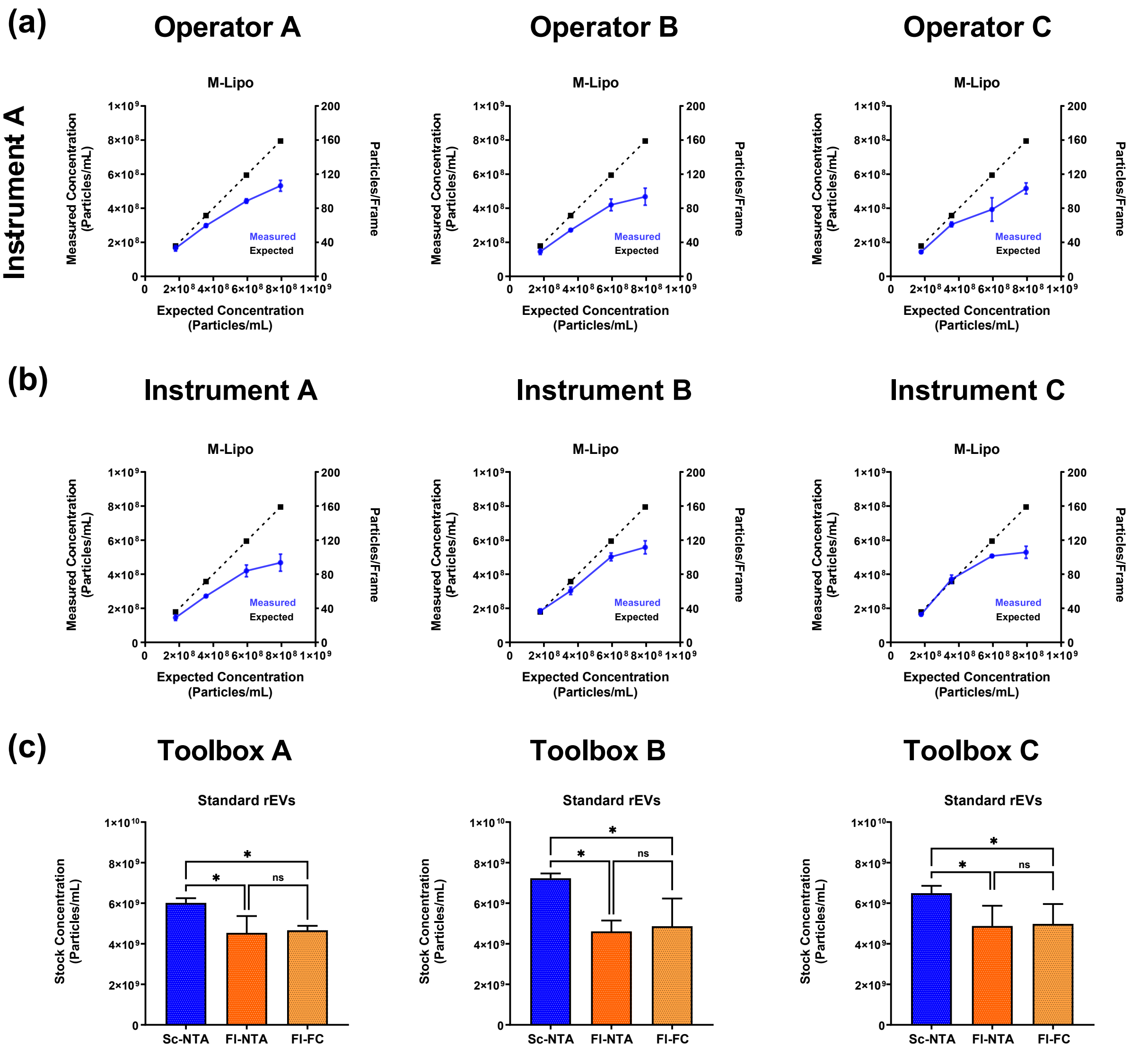


**Supplementary Figure 6. Analytical Optimization;** Variability assessment across multiple operators and instruments.

**(a)** To evaluate operator-related variability, serial dilution experiments with medium-sized liposomes (M-Lipo) were independently conducted by three operators using the same Sc-NTA. All operators demonstrated consistent deviations from the expected linear trend at higher concentrations, confirming that the masking effect is not operator dependent. **(b)** To assess instrument-specific variability, the same dilution experiments were repeated on three independent NTA instruments located in separate laboratories. The masking effect was consistently observed across all systems, indicating that this phenomenon is intrinsic to the technology under conditions of high particle concentration and/or size heterogeneity. **(e)** To evaluate reproducibility under optimal assays, stock concentrations of recombinant EVs (rEVs) were measured across three independently maintained analytical toolboxes (3 pairs of NTA and

F-FC). Similar trends in concentration measurements were observed across different setups. Multicomparison statistical analysis was performed using ordinary one-way ANOVA with Tukey's multiple comparison test (* $p<0.05$, ** $p<0.01$, *** $p<0.001$, **** $p < 0.0001$ ; ns = not significant). All data are presented as mean ± standard deviation with 3 independent measurements. Sc-NTA: Scattering mode of NTA, Fl: Fluorescence mode of NTA, Fl-FC: Fluorescence-based flow cytometry.

### Variation and reproducibility Study:

The Serial dilution experiment with medium size liposomes was repeated on 3 NTA (Malvern) instruments; two located at Sartorius Stedim North America, Marlborough, MA facility and one located at Dr. Gaborski's lab at Rochester Institute of Technology. Same serial dilution steps and same source of medium size liposomes were used. The stock concentration of standard rEVs was then measured to study the reproducibility of the established analytical toolbox with 3 pairs of NTA and Fl-FC machines. The standard rEV samples were purchased from the same vendor and reconstituted as previously described.

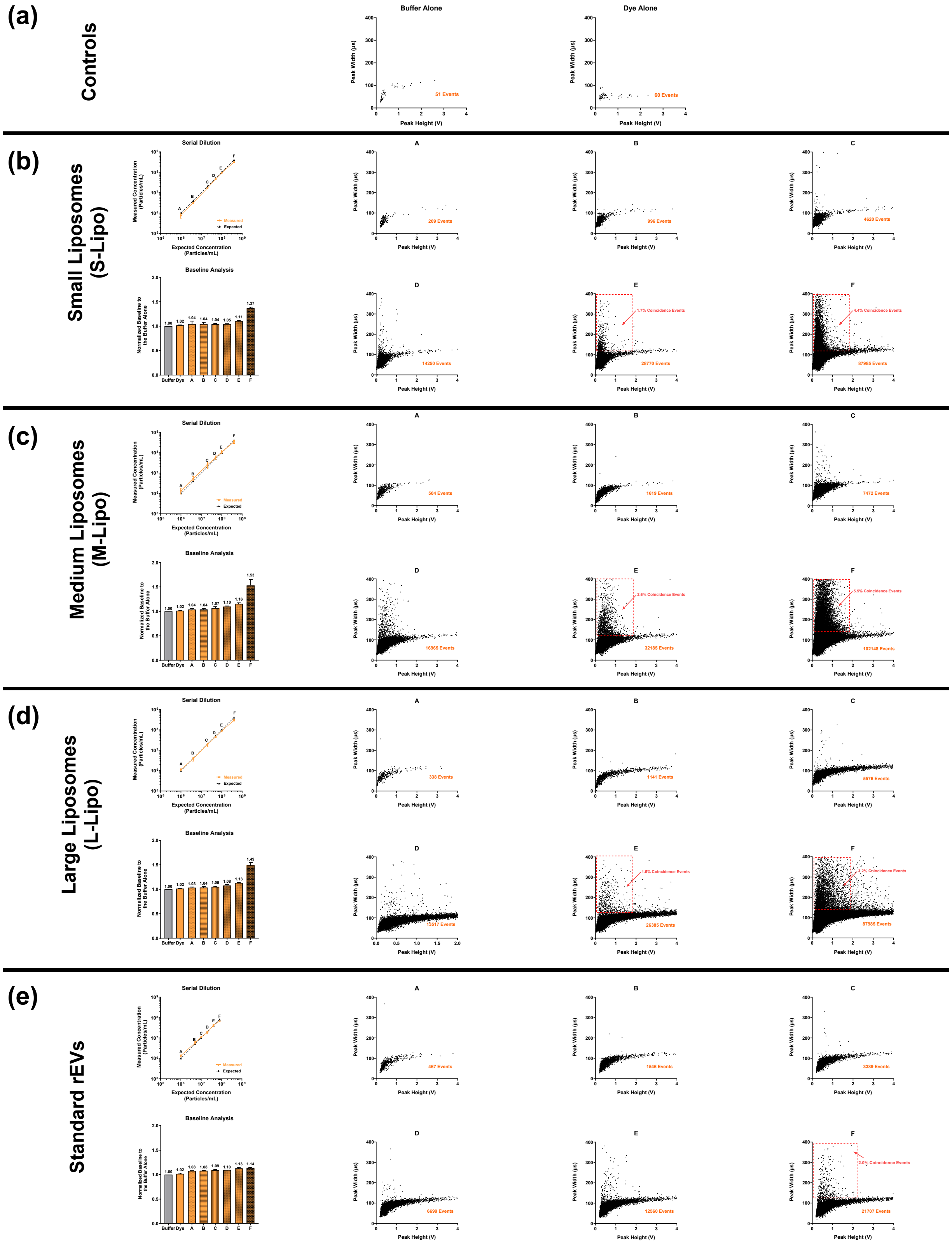
(a)
Controls
Buffer Alone
51 Events
Dye Alone
60 Events
Peak Width (µs)
Peak Height (V)
(b)
Small Liposomes
(S-Lipo)
Serial Dilution
Measured Concentration (Particles/mL)
Expected Concentration (Particles/mL)
Measured
Expected
Baseline Analysis
Normalized Baseline to the Buffer Alone
Buffer Dye A B C D E F
1.00 1.02 1.04 1.04 1.04 1.05 1.11 1.37
209 Events
996 Events
4620 Events
14250 Events
25770 Events
57985 Events
1.7% Coincidence Events
4.4% Coincidence Events
(c)
Medium Liposomes
(M-Lipo)
1.00 1.02 1.04 1.04 1.07 1.10 1.16 1.53
504 Events
1619 Events
7472 Events
16965 Events
32185 Events
102148 Events
2.6% Coincidence Events
5.5% Coincidence Events
(d)
Large Liposomes
(L-Lipo)
1.00 1.02 1.03 1.04 1.05 1.08 1.13 1.49
338 Events
1141 Events
5576 Events
13517 Events
26385 Events
57985 Events
1.5% Coincidence Events
2.2% Coincidence Events
(e)
Standard rEVs
1.00 1.02 1.08 1.08 1.09 1.10 1.13 1.14
467 Events
1546 Events
3389 Events
6699 Events
12560 Events
21707 Events
2.0% Coincidence Events

**Supplementary Figure 7. Analytical Optimization;** Detailed analysis of serial dilution experiments on fluorescence-based flow cytometry (Fl-FC).

**(a–c)** Buffer alone and dye alone as negative controls showed minimal background signal Fl-FC, confirming the optimized no-wash labeling protocol. **(b–e)** Validation data for small (S-Lipo), medium (M-Lipo), and large (L-Lipo) liposomes, as well as recombinant extracellular vesicles (rEVs), analyzed using Fl-FC:

- Left panels: Serial dilution plots demonstrate strong linear correlation between expected and measured concentrations across a broad dynamic range.
- Middle panels (A–F): 2D plots of peak height vs. peak width illustrate particle distribution at various dilutions. At high concentrations, a distinct population with broadened peak widths emerges (regions D–F), indicative of coincidence events.
- Baseline fluorescence analysis confirms consistent fluorescence intensity across dilutions, with elevation at the highest concentrations.
- Right panels: Detergent treatment with SDS resulted in a substantial decrease in particle counts (80–92% reduction), verifying that detected particles are membrane-enclosed vesicles rather than dye aggregates or non-vesicular particles. All data are presented as mean ± standard deviation with 3 independent measurements.

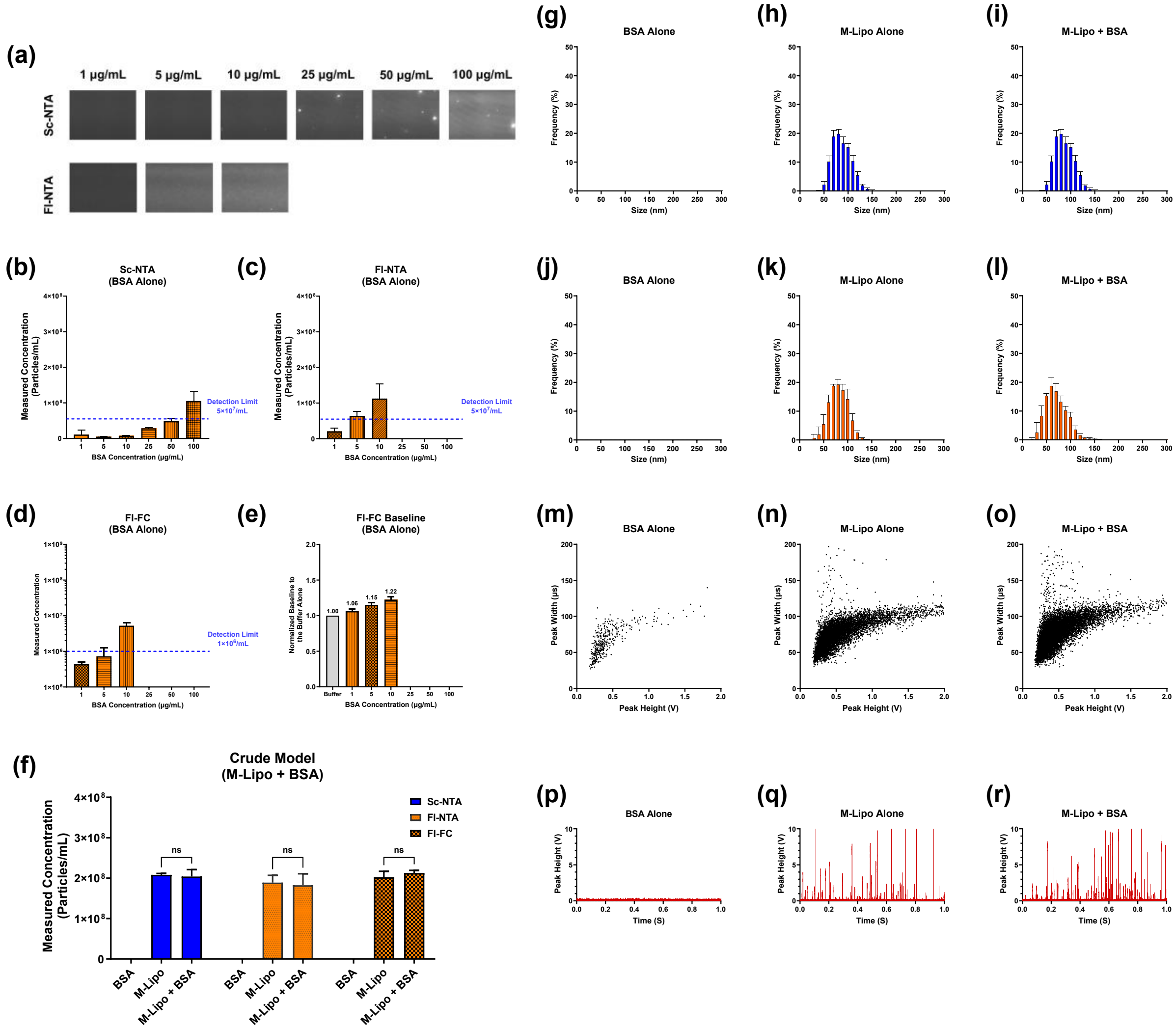


**Supplementary Figure 8. Analytical Optimization;** Evaluation of BSA interference on single particle analytics using a crude sample model.

**(a)** Representative NTA frames of BSA alone samples at increasing concentrations (1–100 μg/mL) analyzed using Sc-NTA (top row) and Fl-NTA (bottom row). Visual cloudiness and particle like signals were observed at ≥50 μg/mL in Sc-NTA and ≥10 μg/mL in Fl-NTA. **(b–d)** Quantification of BSA-alone samples on **(b)** Sc-NTA, **(c)** Fl-NTA, and **(d)** fluorescence-based flow cytometry (Fl-FC). The dashed blue lines indicate the detection thresholds for each technique. **(e)** Baseline fluorescence analysis of BSA alone samples in Fl-FC. Concentrations ≥10 μg/mL resulted in elevated baseline values. **(f)** Measured concentration of M-Lipo alone ($2 \times 10^8$ Particles/mL), BSA alone (5 μg/mL), and M-Lipo+BSA (1:1 ratio, $4 \times 10^8$ Particles/mL + 10 μg/mL of BSA) across Sc-NTA, Fl-NTA, and Fl-FC. Measured concentrations were not significantly affected by BSA contamination (ns = not significant). **(g–l)** Size distribution profiles of BSA alone, M-Lipo alone, and M-Lipo + BSA mixtures in **(g-h)** Sc-NTA and **(i-l)** Fl-NTA, showing no changes in distribution caused by BSA contamination. **(m-o)** 2D peak height vs. width plots and **(p-r)** one-second fluorescence histograms from Fl-FC analysis. No shifts or abnormal signal profiles were observed in the presence of BSA, confirming measurement accuracy. Multicomparison statistical analysis was performed using ordinary two-way ANOVA with Tukey's multiple comparison test (*p<0.05, **p<0.01, ***p<0.001, ****p < 0.0001; ns = not significant). All data are presented as mean ± standard deviation with 3 independent measurements. Sc-NTA: Scattering mode of NTA, Fl: Fluorescence mode of NTA, Fl-FC: Fluorescence-based flow cytometry.

### Crude Sample Model (BSA + Liposome) Study:

Pierce™ bovine serum albumin standards (2 mg/mL) was purchased from ThermoFisher Scientific. The stock was then serially diluted to 100, 50, 25, 10, 5, 1 μg/mL. The BSA alone control samples were then tested first on scattering mode of NTA. The samples were then labeled as previously described (with 5 μg/mL of CMO at 1:100 ratio) and ran on fluorescence mode of NTA. The same labeled samples were further diluted 10 times and measured on the Fl-FC instrument.

Medium size liposome samples were then diluted to $2 \times 10^8$ particles/mL and tested on Sc-NTA, labeled and measured in Fl-NTA and Fl-FC first. The liposome samples ($4 \times 10^8$ particles/mL) were then mixed with BSA (5 μg/mL) at a 1:1 ratio. The samples were again measured with Sc-NTA and labelled and measured on Fl-NTA and Fl-FC.

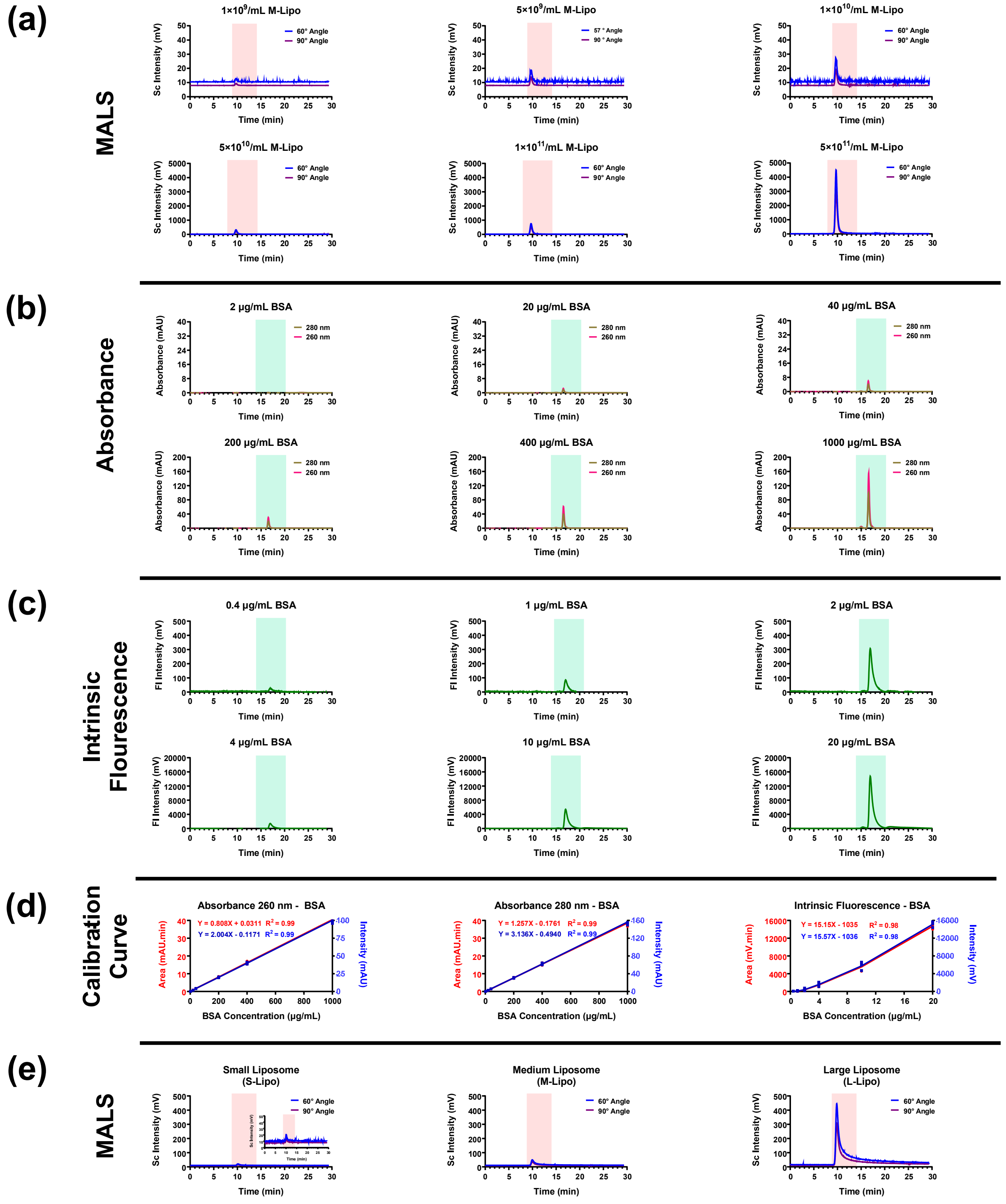


**Supplementary Figure 9. Analytical Optimization;** Establishing analytical assays using multi-detector analytical chromatography.

**(a)** Multi-angle light scattering (MALS) chromatograms for serial dilutions of medium-sized liposomes (M-Lipo) from $1{\times}10^9$ to $5{\times}10^{11}$ particles/mL. Liposomes eluted between 9–14 minutes (highlighted in red). **(b)** UV absorbance chromatograms for bovine serum albumin (BSA) standards at concentrations ranging from 2 to 1000 µg/mL, showing a consistent elution between 14–20 minutes (highlighted in green). **(c)** Intrinsic fluorescence chromatograms (excitation 280 nm, emission 350 nm) for BSA samples from 0.4 to 20 µg/mL. Fluorescence detection exhibited a lower LOD of 0.4 µg/mL and retained BSA in the expected elution window (14–20 min, highlighted in green). **(d)** Calibration curves for BSA across three detection modes: UV absorbance at 260 nm and 280 nm, and intrinsic fluorescence. Peak Intensity and area under the curve (AUC) were plotted against BSA concentration, confirming linear response and quantifiability across the tested range. **(e)** MALS profiles for small (S-Lipo), medium (M-Lipo), and large (L-Lipo) liposomes at the same particle concentration showing the increase in scattering intensity liposome size. Inset in MALS plot is showing the zoomed in version of the plot.

### Analytical Chromatography:

- **Serial Dilution Experiments:**

Medium sized liposomes were serially diluted to $\sim 1 \times 10^9$, ~ $5 \times 10^9$ , ~ $1 \times 10^{10}$, ~ $5 \times 10^{10}$, ~ $1 \times 10^{11}$, ~ $5 \times 10^{11}$ particles/mL and tested on the column to identify the lower limit of detection (LOD) on in-line MALS detector. To assess the lower LOD of UV and fluorescence detectors, BSA standards were also tested by serially diluting in the range of 2, 20, 40, 200, 400 and 1000 µg/mL for UV detector and 0.4, 1, 2, 4, 10 and 20 µg/mL for fluorescence detector.

- **Different Size Liposomes:**

Different sized liposomes (large, medium and small) were tested at the same concentration ($1 \times 10^{11}$ particles/mL) to study the impact of liposome size on in-line MALS signal.

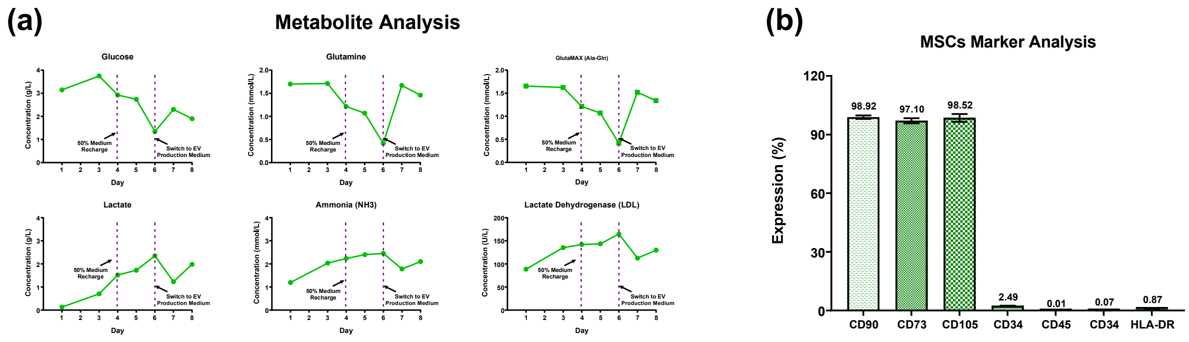


**Supplementary Figure 10. Upstream Processing (UPS);** Metabolite profiling and immunophenotypic characterization of hBM-MSCs during 3D bioreactor culture for EV production.

**(a)** Metabolite analysis of human bone marrow-derived mesenchymal stromal cells (hBM-MSCs) cultured on microcarriers in a 10 L single-use Univessel® bioreactor. Glucose, glutamine, GlutaMAX (Ala-Gln), lactate, ammonia ($NH_3$), and lactate dehydrogenase (LDL) levels were measured daily. A 50% medium exchange on day 4 replenished nutrients, and a full medium exchange to platelet lysate (PLT)-free EV production medium was performed on day 6. Metabolites measurements were done as only 1 replicate. **(b)** Flow cytometry analysis of MSC surface markers at the end of the culture phase. Over 97% of the cell population was positive for CD73, CD90, and CD105, while low expression was detected for CD34 (2.49%), and negligible expression was observed for negative markers CD45 (0.01%), CD3 (0.07%), and HLA-DR (0.87%). All data are presented as mean ± standard deviation with n=3 independent measurements.

### Upstream Processing:

- **Metabolite Analysis:**

Daily medium samples were analyzed via Cedex BioHT (Roche) to determine key metabolite concentration over time.

- **Flow Cytometry Analysis of MSCs Surface Marker:**

MSCs were stained with typical MSC surface markers: CD45 (BD Biosciences, Cat No. 560367), CD73 (BD Biosciences, Cat No. 561258), CD90 (BD Biosciences, Cat No. 328125), CD105 (BD Biosciences, Cat No. 562408), and CD34 (BD Biosciences, Cat No. 348057) and ran through iQue (Sartorius). Negative markers: HLA-DR (BD Biosciences, Cat No. 302244), CD3 (Biolegend, Cat No. 344842), and CD45 (BD Biosciences, Cat No. 560367).

Cells were quickly thawed and centrifuged at 400 x g for 5 minutes to pellet the cells. The cell pellet was then resuspended in Staining Buffer (BD) at a concentration of $2 \times 10^6$ cells/mL. Antibodies were diluted according to the manufacturer's instructions, and 75 µL of the diluted antibody solution was added to the cells. The reactions were incubated for 30 minutes in the dark. Following incubation, the cells were washed three times with Staining Buffer. The samples were then resuspended in 100 µL of Staining Buffer, transferred to a 96-well plate, and loaded onto the iQue3 cytometer. Sample analysis was conducted using the iQue3 Forecyt® Software.

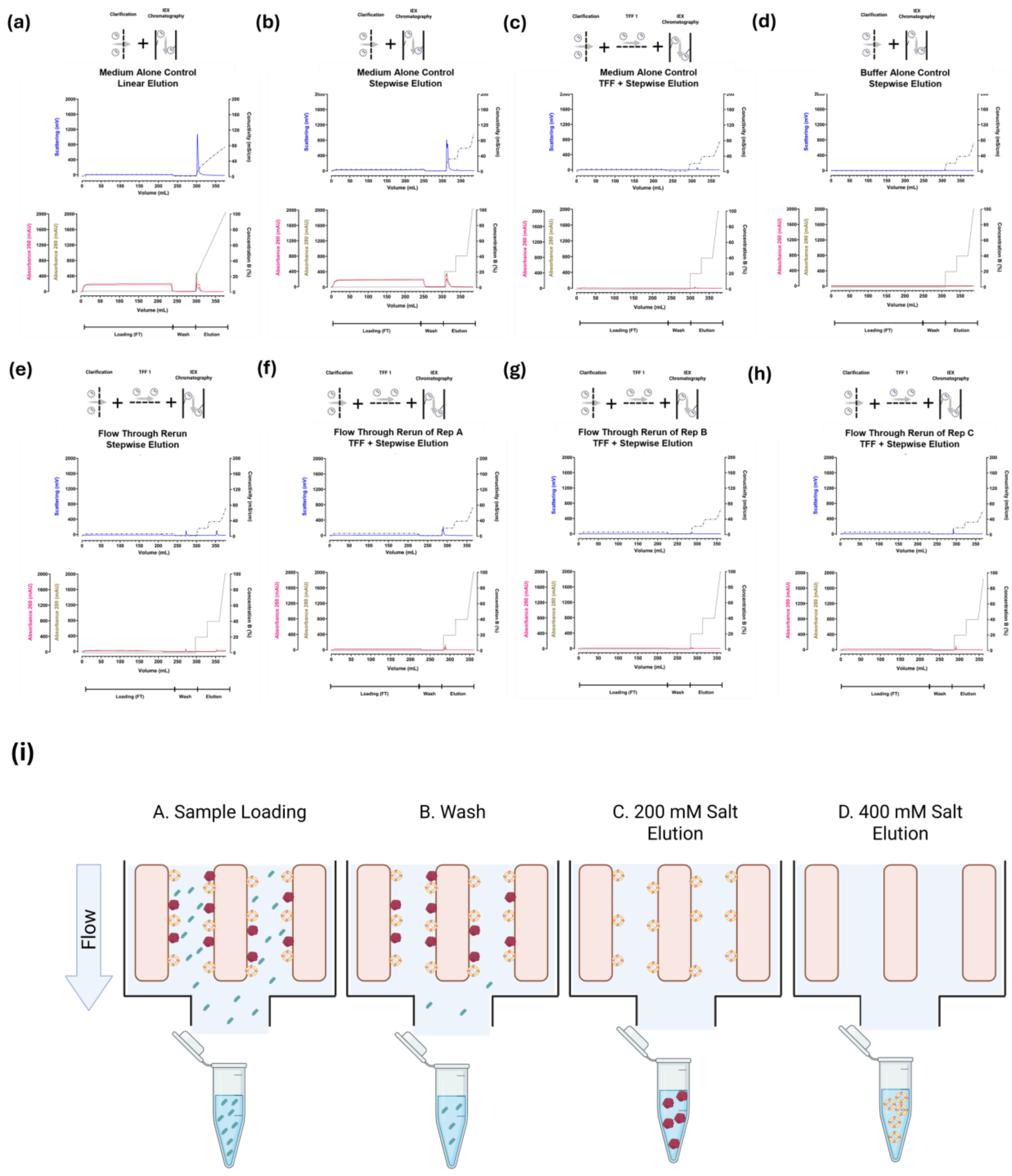


**Supplementary Figure 11. Downstream Processing (DSP);** Control experiments and flow-through re-run confirm the specificity and efficiency of the DSP workflow for MSC-EV enrichment.

**(a)** Chromatographic analysis of clarified unconditioned medium (medium alone control) subjected to linear salt elution strategy. **(b)** Medium alone control processed via stepwise elution strategy exhibited a strong RT-MALS and UV signal at 200 mM NaCl and no peak at 400 mM, indicating presence of medium-associated components. **(c)** Medium processed by clarification and 6× diafiltration prior to IEX resulted in no detectable UV or RT-MALS signals during step elution, confirming effective removal of medium-associated components by TFF. **(d)** Clarified buffer-alone control also produced no signal in either detector, further validating system cleanliness and the specificity of elution signals observed in MSCs CM runs. **(e-h)** Flow through (FT) samples from each MSCs CM run **(Fig 5-d, e and f)** were reloaded onto the IEX column and subjected to the same stepwise elution strategy. Across all reruns, no detectable peaks were observed in UV or RT-MALS detectors, indicating efficient initial capture of bindable components during the first column load and minimal loss of target analytes in the flow through (FT). **(i)** IEX schematic showing stepwise elution strategy. Schematics were created in Biorender.com.

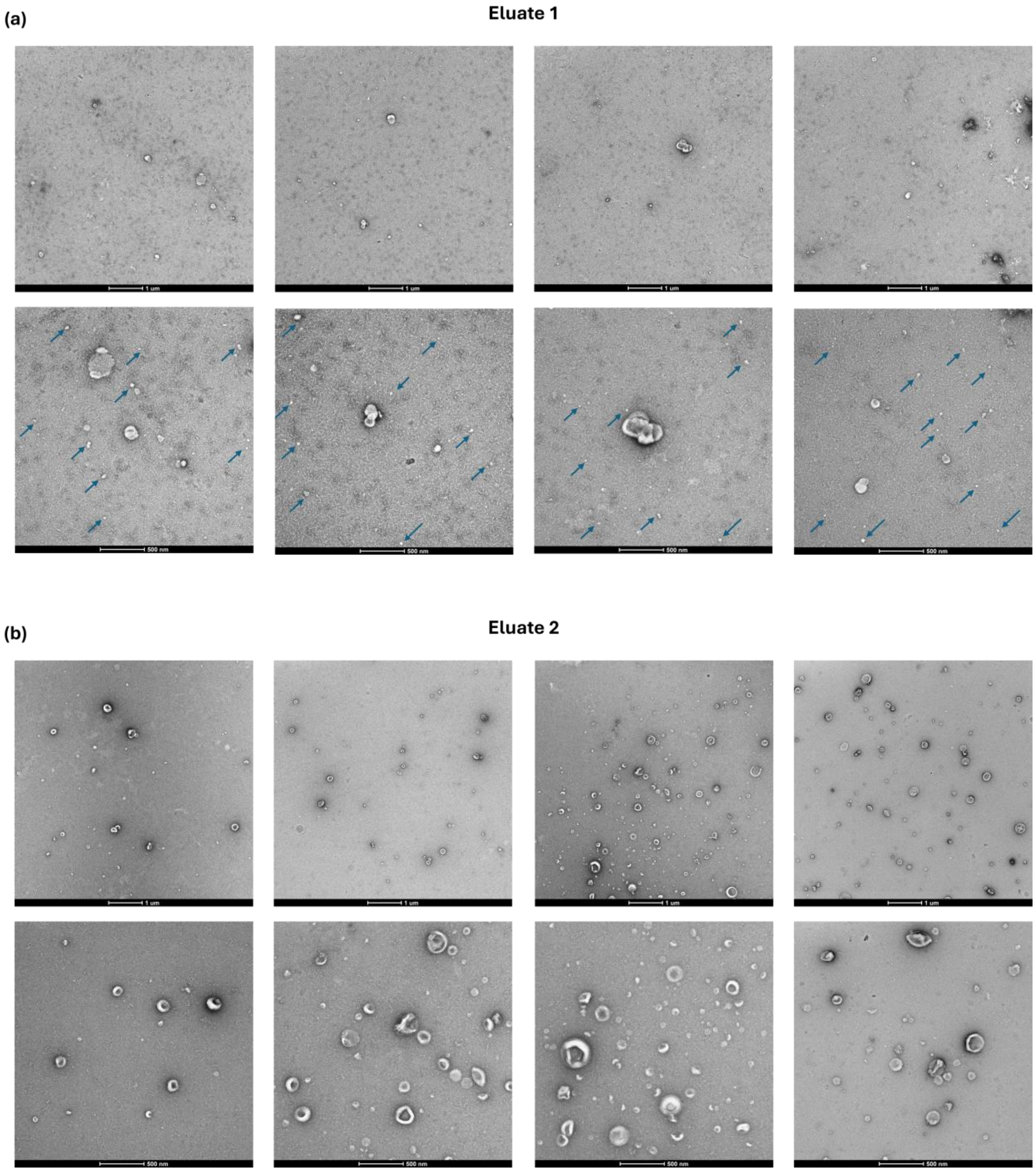


**Supplementary Figure 12. Identity Investigation;** Transmission Electron Microscopy (TEM) images of IEX eluates.

**(a)** Additional TEM images of the first eluate obtained from MSCs CM showing lack of vesicular structures and presence of contaminants. Scale bars = 500 nm and 1 µm.**(b)** Additional TEM images of the second eluate obtained from MSC-conditioned medium are shown at two magnifications, highlighting the presence of vesicle-like structures with characteristic cup-shaped morphology. Scale bars: 500 nm and 1 µm.

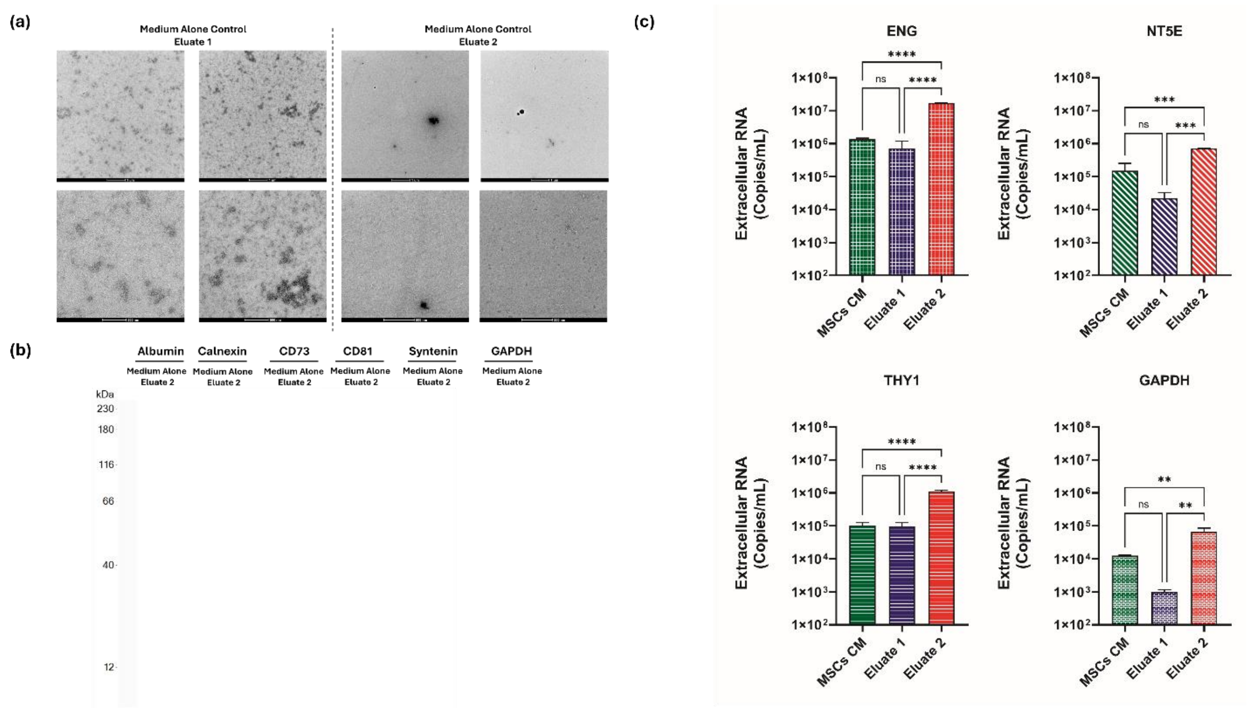


**Supplementary Figure 13. Identity Investigation;** Additional Data from Transmission Electron Microscopy, Simple Western and Reverse Transcription quantitative PCR (RT-qPCR) analysis.

**(a)** Additional TEM images of the first and second eluates obtained from medium alone control showing the absence of the observed vesicular and non-vesicular structures from MSCs CM. Scale bars: 500 nm and 1 µm. **(b)** Simple Western analysis of equivalent eluate 2 of medium alone control showing no detectable signal for any of the markers tested, including EV-associated and contaminant markers. **(c)** Expression analysis of well-established MSCs mRNA markers - *THY1* (Thy1/CD90), *ENG* (Endoglin/CD105), and *NT5E* (ecto-5′-nucleotidase/CD73) using RT-qPCR. Eluate 2 (10mL) was significantly enriched in all three mRNA species compared to eluate 1 (10 mL) and MSCs conditioned medium (MSCs CM, 100 mL). GAPDH mRNA, as a control, also showed higher expression level in eluate 1. All data are presented as mean ± standard deviation with n= 3 independent measurements. Unless otherwise specified, Multicomparison statistical analysis was performed using ordinary one-way ANOVA with Tukey's multiple comparison test (*$p<0.05$, **$p<0.01$, ***$p<0.001$, ****$p < 0.0001$; ns = not significant).

### RNA isolation and reverse transcription quantitative PCR (RT-qPCR) :

Reverse transcription quantitative real-time PCR (RT-qPCR) analysis of RNA from extracellular environment was performed on the samples of equal volume from MSCs CM, and eluates 1 and 2. The samples were used to isolate RNA by silica-column method by

PureLink™ RNA Mini Kit (Invitrogen™, cat# 11754050) according to manufacturer's guidelines. Briefly, approximately 200 µL of sample was mixed with 200 µL of the lysis buffer, containing 1% β-Mercaptoethanol and 200 µL of molecular grade ethanol and the resulting homogenate was vortexed. Next, the homogenate was passed through the silica column for RNA capture via centrifugation at 12,000 x g for 20 s and flow through was discarded. Subsequently, the column was washed with Wash Buffer I and then with Wash Buffer II, and dried by 2 minutes centrifugation at 12,000 x g. The silica spin column was transferred to a new 1.5 mL tube, and RNA was eluted by 30 µL of nuclease-free water.

Reverse transcription of total RNA was performed with the use of SuperScript™ VILO™ cDNA synthesis kit (Invitrogen™, cat# 11754050) according to the manufacturer's instructions with the addition of 1 µL of 100 µM Oligo dT (15) primer (5′-TTT TTT TTT TTT TTT-3′) per 20 µL of reaction mixture.

Next, qPCR was performed with 5 µL of 2:3 and 5x diluted cDNA using iTaq™ Universal SYBR® Green Supermix (Bio-Rad, cat#1725120) using the following pairs of primers: 1) GAPDH-specific primers GAPDH-Forward (5′-ACC ACA GTC CAT GCC ATC AC -3′, Tm = 57.7°C) and GAPDH-Reverse (5′-TCC ACC ACC CTG TTG CTG TA-3′, Tm = 58.3°C); 2) THY1-specific primers THY1-F (5′-G AAG GTC CTC TAC TTA TCC GCC-3′, Tm = 56.6°C) and THY1-R (5′-TGA TGC CCT CAC ACT TGA CCA G -3′, Tm = 59.3°C); 3) NT5E-specific primers NT5E-F (5′-AGT CCA CTG GAG AGT TCC TGC A-3′, Tm = 60.0°C) and NT5E-R (5′- TGA GAG GGT CAT AAC TGG GCA C-3′, Tm = 58.7°C); 4) ENG-specific primers ENG-F (5′- CGG TGG TCA ATA TCC TGT CGA G-3′, Tm = 57.2°C) and ENG-R (5′- AGG AAG TGT GGG CTG AGG TAG A -3′, Tm = 59.6°C).

Serially diluted human male genomic DNA (MilliporeSigma™, cat#705723) was used to generate standard curve with the following PCR conditions: 50°C for 2 min (1 cycle), 95°C for 10 min (1 cycle), 95°C for 15 s, and 60°C for 60 s (41 cycles). Quantification of RNA (copies/mL) was determined from each sample's average cycle quantification (Cq) value relative to the standard curve. The amplification was carried out in technical triplicate on CFX Opus 96 Real-Time PCR System (Bio-Rad, cat#12011319).

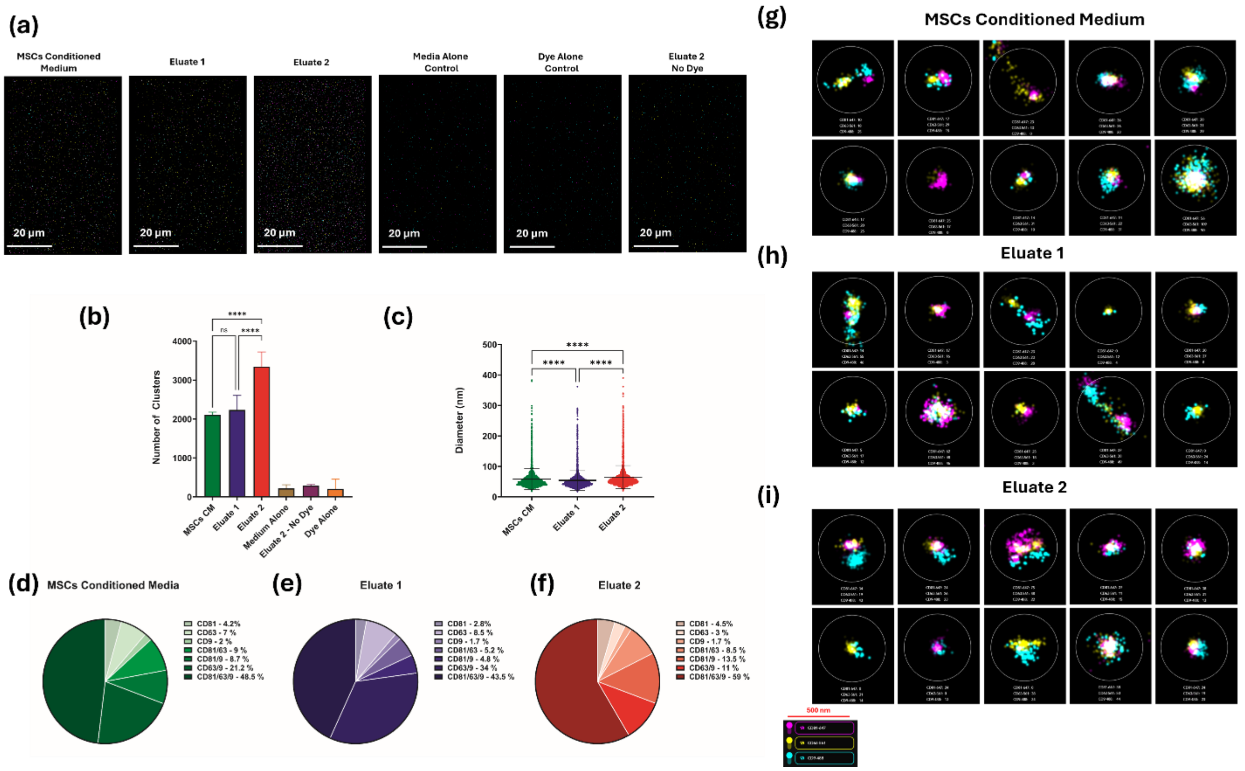


**Supplementary Figure 14. Identity Investigation;** Tetraspanins (CD9, CD63 and CD81) expression analysis of IEX eluates using super-resolution imaging.

**(a)** Representative direct stochastic optical reconstruction microscopy (dSTORM) images showing CD9-, CD63-, and CD81-labeled clusters from MSCs conditioned medium (MSCs CM), eluate 1, eluate 2, and control samples (medium alone eluate, dye alone control, and unlabeled eluate 2). Scale bars: 20 µm. All data are presented as mean ± standard deviation (n=5, independent field of views).

**(b)** Quantification of detected EV clusters, showing significantly higher cluster counts in Eluate 2 compared to eluate 1 and MSCs CM, and markedly lower counts in all controls. All data are presented as mean ± standard deviation (n=5, independent field of views).

**(c)** Violin plots of EV cluster diameters, indicating larger average particle sizes in eluate 2, followed by MSCs CM and eluate 1.

**(d–f)** Pie charts illustrating the distribution of single-, double-, and triple-positive clusters for CD9, CD63, and CD81. Eluate 2 exhibited the highest proportion of triple-positive clusters (59%), while eluate 1 displayed increased double positivity for CD63 and CD9 (34%) compared to eluate 2 (11%). All data are presented as mean (n=5, independent field of views).

**(g–i)** Representative zoom-in dSTORM reconstructions of individual clusters. Unless otherwise specified, Multicomparison statistical analysis was performed using ordinary one-way ANOVA with Tukey's multiple comparison test (*p<0.05, **p<0.01, ***p<0.001, ****p < 0.0001; ns = not significant).

### Direct Stochastic Optical Reconstruction Microscopy (dSTORM):

Both IEX eluates as well control samples (unlabeled eluate 2, dye Alone, MSCs CM and medium alone eluate 2) were immunolabeled and imaged using the EV Profiler Kit (#EV-MAN-2.0, ONI) by dSTORM. Briefly, samples were immobilized on microfluidic chips, fixed with F1 solution (provided in the kit) for 10 min, and incubated for 50 min with fluorescently labeled antibodies. The following antibodies were used, either provided in the kit: CD9-CF488 (kit, excitation (ex)/emission (em): 490/515 nm), CD63-CF568 (kit, ex/em: 562/583 nm), and CD81-CF647 (kit, ex/em: 650/665 nm). Finally, samples were again fixed with F1 for 5 min, and a freshly prepared dSTORM-imaging buffer was added prior to image acquisition. Labeled samples were imaged using the Nanoimager S Mark III microscope (ONI) with 100× oil-immersion objective. Sequential 3-color imaging was performed using recommended manufacturer laser power for the 640, 561, and 488 nm lasers, respectively, at 1000 frames per channel with the angle of illumination set to 54°. Prior to the start of the imaging session, channel mapping was calibrated using 0.1 μm TetraSpeck beads (T7279, Thermo Fisher Scientific). Data was acquired and processed by AutoEV, ONI. Subpopulation analyses of EVs that express one, two, or three markers were analyzed using ONI's online platform CODI (https://alto.codi.bio). Briefly, drift correction, filtering, clustering, and counting algorithms were performed using the predefined "EV Profiling" workflow.

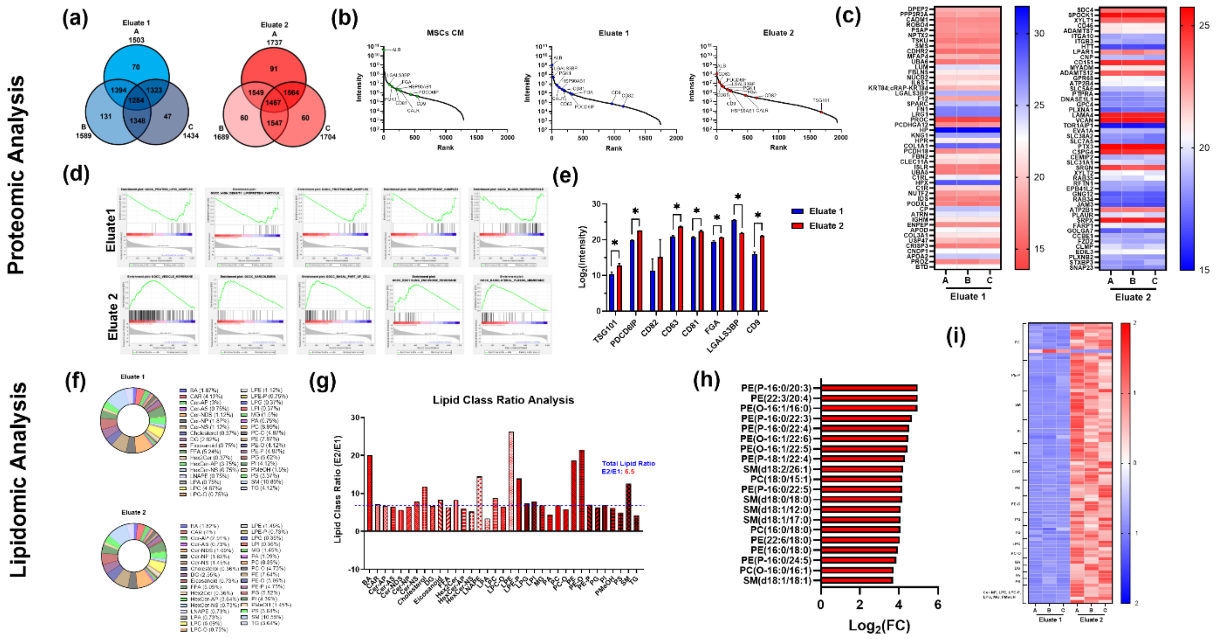


**Supplementary Figure 15. Identity Investigation;** Proteomic and lipidomic characterization of IEX eluates.

**(a-e)** Proteomic Analysis; **(a)** Venn diagrams showing overlap of protein signatures across three biological replicates for eluate1 (left) and eluate 2 (right), illustrating a shared core set of proteins with a smaller number of replicate-specific proteins. **(b)** Protein rank plots for MSCs conditioned medium (MSCs CM), eluate 1, and eluate 2, highlighting distribution of intensities for EV-associated markers (CD63, CD81, CD82, CD9, PDCD6IP/Alix, TSG101), non-vesicular particle-associated markers (LGALS3BP), and controls for cell debris (CALR) and serum protein (ALB). **(c)** Heatmaps of the top 50 differentially expressed proteins between the two eluates, with blue and red indicating lower and higher relative abundance, respectively. **(d)** Gene Set Enrichment Analysis (GSEA) revealing enrichment in lipid–protein complex components, HDL-associated proteins, and proteasome subunits in eluate 1, and vesicle membrane formation proteins, ESCRT-associated machinery, and membrane transporters in eluate 2. **(e)** Protein expression level plot for selected markers, showing differential intensity distributions across samples and indicating enrichment of common EV markers in eluate 2 after IEX step. **(f-i)** Lipidomic Analysis; **(f)** Lipid composition analysis showing the number of lipid compounds in each detected subclass. **(g)** Lipid class ratio plot (E2/E1), with a dashed line indicating the total lipid abundance ratio (6.5-fold); several lipid classes, including BA, cholesterol, and multiple glycerophospholipid subclasses, are enriched above this threshold in E2. **(h)** Bar plot showing the top 20 differentially expressed lipids between eluates, presented as $\log_2$ fold change values. **(i)** Heatmap of differentially expressed lipid species illustrating relative abundance patterns between eluates.

### Proteomic Analysis:

- Data Processing:

DIA data were analyzed using DIA-NN (v1.9.1). Database searches were launched against Swissprot Human. The digestion enzyme was set as trypsin. Carbamidomethylation of cysteine was set as fixed modification. Oxidation of methionine was specified as a variable modification. Matched peptides and protein were filtered using a 1% false discovery rate (FDR). The MS/MS spectra were searched with a precursor mass tolerance of 10 ppm and a fragment ion mass tolerance of 0.6 Da. Protein quantification was achieved by using total intensities of all precursors.

- Analysis:

Proteins with a signal in 2 of 3 replicates were used for analysis, performed in R 4.5.1. First, intensity values were transformed into log2 and proteins in medium alone eluate 1 and eluate 2 with a z-score > -3 with respect to eluate 1 and 2 were removed. Next, the limma package (Ritchie et al., 2015) was used to fill missing values by left-censored imputation. Then, a linear model was fit to the data (lmFit), empirical bayes moderation was applied (eBayes), and significant differential expression was determined using the Benjamini Hochberg procedure with a false discovery rate of 5% (topTable). The $\log_2$ intensity values were z-score normalized for the heatmap in **Fig 6**. PCA was performed in GraphPad Prism 10.4.2.633 using the correlation matrix. Gene set enrichment analysis was performed in GSEA 4.4.0 using GSEA Preranked with $\log_2$(FC) values against the gene ontology cellular component gene set (c5.go.cc.v2025.1.Hs.symbols.gmt, 1042 gene sets) in the Molecular Signatures database (https://www.gsea-msigdb.org/gsea/msigdb/human/genesets.jsp?collection=GO:CC). Enriched cellular components were determined using default settings and a false discovery rate of 5%.

### Lipidomic Analysis:

- Data Processing

Lipids were annotated based on retention time and ion-pair information from MRM mode using the MetawareBio in-house database. In MRM mode, the first quadrupole screened the precursor ions for target substance and excluded ions of other molecular weights. After ionization was induced by the impact chamber, the precursor ions were fragmented, and a characteristic fragment ion was selected through the third quadrupole to exclude the interference of other non-target ions. By selecting a particular fragment, quantification is more accurate and reproducible. Analyst 1.6.3 was used to process MS data. The MS peak of each lipid in different samples was corrected according to the retention time and peak distribution information to ensure the accuracy of the analysis. Lipids were quantified based on a calibration curve using the formula:

$$x = \frac{\frac{RcFV}{m}}{n}$$

Where *x* is the lipid content of the sample (pmol/mg), *R* is the ratio of the peak area of the substance to be measured to the peak area of the internal standard, *F* is the internal standard correction factor for different types of substances, c is the concentration of the internal standard (μmol/L), *v* is the extraction solution volume for the sample (μL), m is the weighed sample size (mL), and *n* is the sample protein concentration (mg/mL). During quantitative analysis, a method of quantitative protein correction of metabolomic data was used based on the central rule to reduce quantitative errors caused by differences in sample size. That is, the protein concentration was determined using the BCA method, and the metabolomic data of the test samples were corrected with the protein concentration of the parallel sample.

Lipidomics data analysis was performed using R 3.5.1. PCA was performed with the *prcomp* function and the parameters *scale*=FALSE and *center*=FALSE. Differentially expressed lipids were determined using orthogonal partial least squares discriminant analysis to calculate variable importance (VIP) scores in MetaboAnalystR 1.0.1. Lipid species with VIP >1 and p-value <0.05 were considered significantly differentially expressed. All heatmap values are z-score normalized.

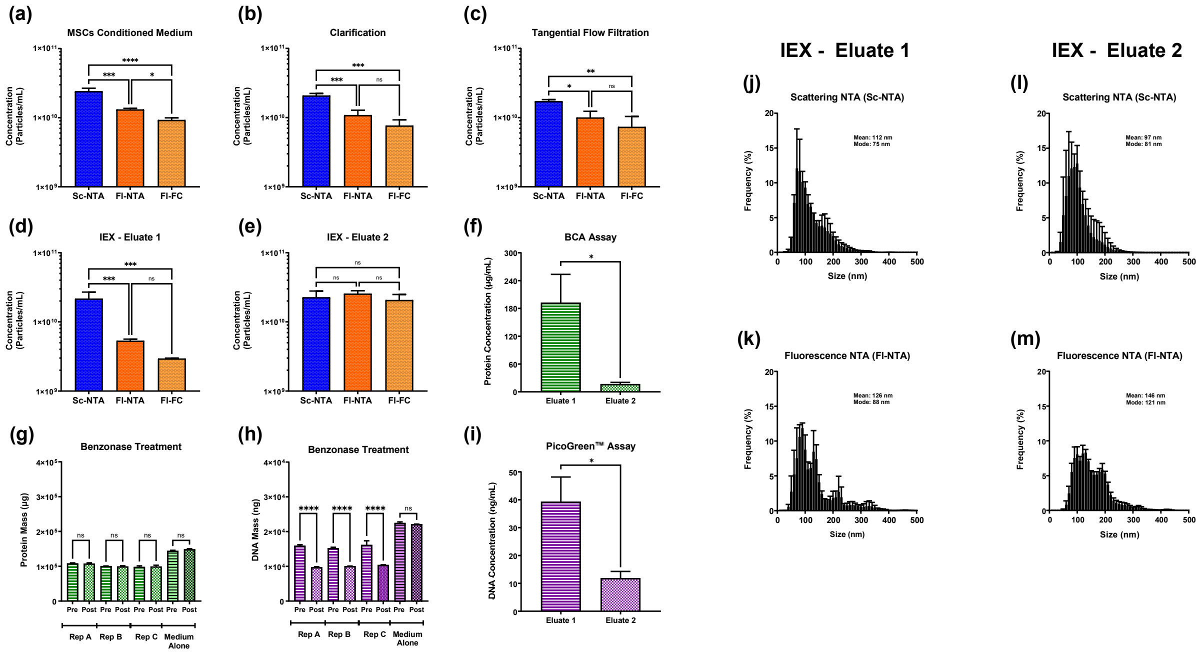


**Supplementary Figure 16. Downstream Processing (DSP) Monitoring;** Step by step DSP evaluation using orthogonal analytics to assess particle recovery and impurity removal. **(a-e)** Concentration measurements using single particle analysis techniques (SC-NTA, FI-NTA and FI-FC); **(a)** MSCs conditioned mediaum(MSCs CM), **(b)** Clarification, **(c)** Tangential Flow Filtration, **(d)** Ion Exchange Chromatography (IEX) eluate 1, **(e)** eluate 2. **(f)** Protein concentration measured by BCA assay for both eluate 1 and 2. Statistical analysis was performed using unpaired t-test with Welch's correcting (*p<0.05). **(g)** The impact of Benzonase treatment on protein mass in MSCs CM and medium alone control. **(h)** The impact of Benzonase treatment on DNA mass in MSCs CM and medium alone control measured by PicoGreen™ assay. **(f)** DNA concentration measured by PicoGreen ™ assay for eluate 1 and 2. Statistical analysis was performed using unpaired t-test with Welch's correcting (*p<0.05). **(j-m)** size distribution plots eluate 1 and eluate 2 using SC-NTA and FI-NTA. All data are presented as mean ± SD with n=3 independent DSP runs. Unless otherwise specified, Multicomparison statistical analysis was performed using ordinary one-way ANOVA with Tukey's multiple comparison test (*p<0.05, **p<0.01, ***p<0.001, ****p < 0.0001; ns = not significant). Sc-NTA: Scattering mode of NTA, FI: Fluorescence mode of NTA, FI-FC: Fluorescence-based flow cytometry.

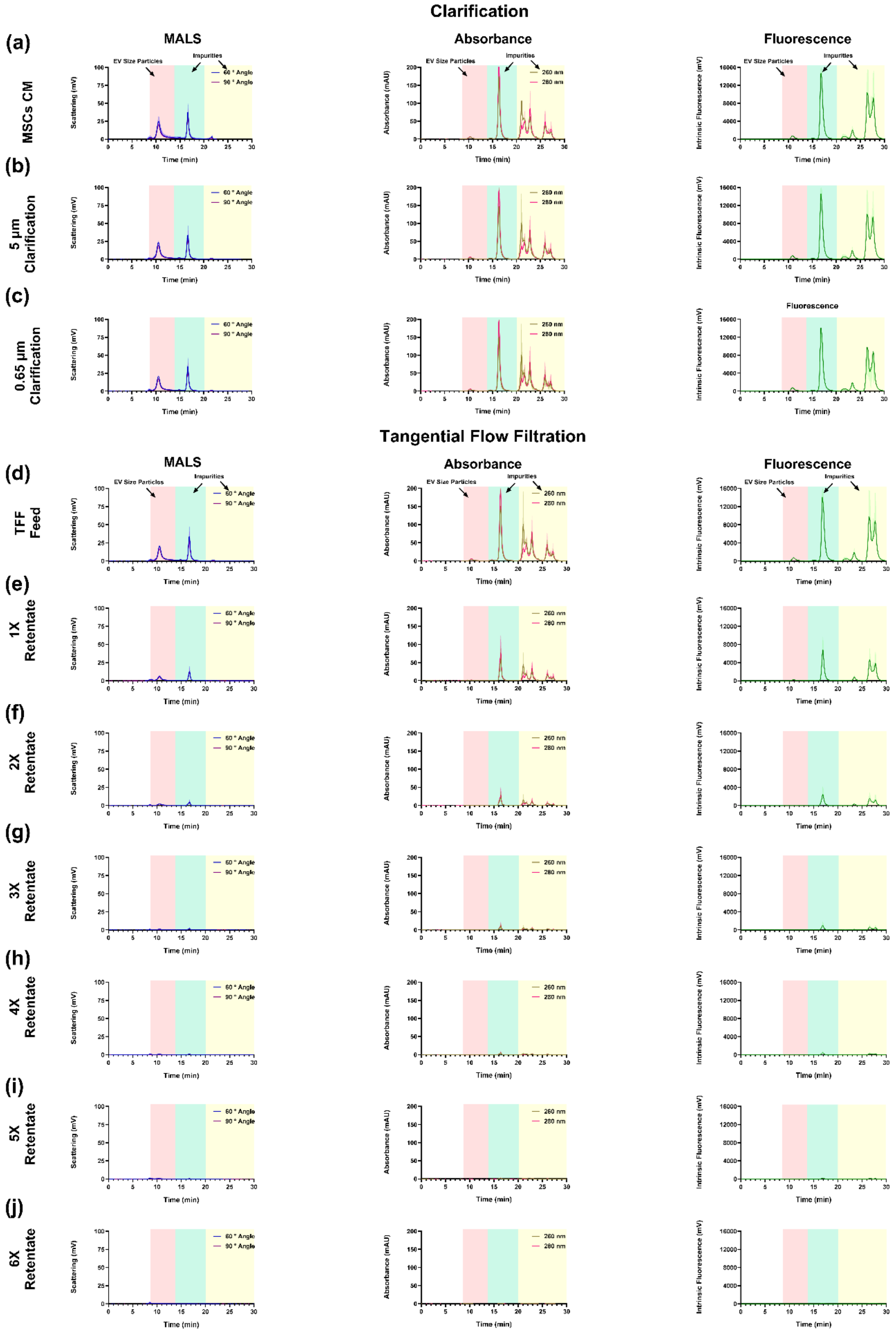

Clarification
(a) MSCs CM
(b) 5 µm Clarification
(c) 0.65 µm Clarification
MALS
Absorbance
Fluorescence
EV Size Particles
Impurities
60 ° Angle
90 ° Angle
260 nm
280 nm
Scattering (mV)
Absorbance (mAU)
Intrinsic Fluorescence (mV)
Time (min)
Tangential Flow Filtration
(d) TFF Feed
(e) 1X Retentate
(f) 2X Retentate
(g) 3X Retentate
(h) 4X Retentate
(i) 5X Retentate
(j) 6X Retentate

**Supplementary Figure 17 - Downstream Processing (DSP) Monitoring;** Step by step DSP evaluation using analytical chromatography profiles of medium alone control across clarification and tangential flow filtration (TFF) steps.

Size exclusion chromatography coupled with in-line multi-angle light scattering (MALS), UV absorbance, and intrinsic fluorescence detection was used to monitor particle and impurity distribution during processing of medium alone control.

**(a–c)** Clarification steps (5 µm and 0.65 µm filters) showing strong UV and fluorescence peaks primarily in impurity-associated regions (14–20 min, highlighted in green) and late-eluting small soluble impurities (>20 min, highlighted in yellow).

**(d–j)** Sequential TFF diafiltration (1–6X retentate) progressively reduced impurity peaks, with complete removal of 14-20 min and late-eluting small soluble impurities showing effective removal of medium-associated components. All data are presented as mean ± SD with n=3 independent DSP runs.

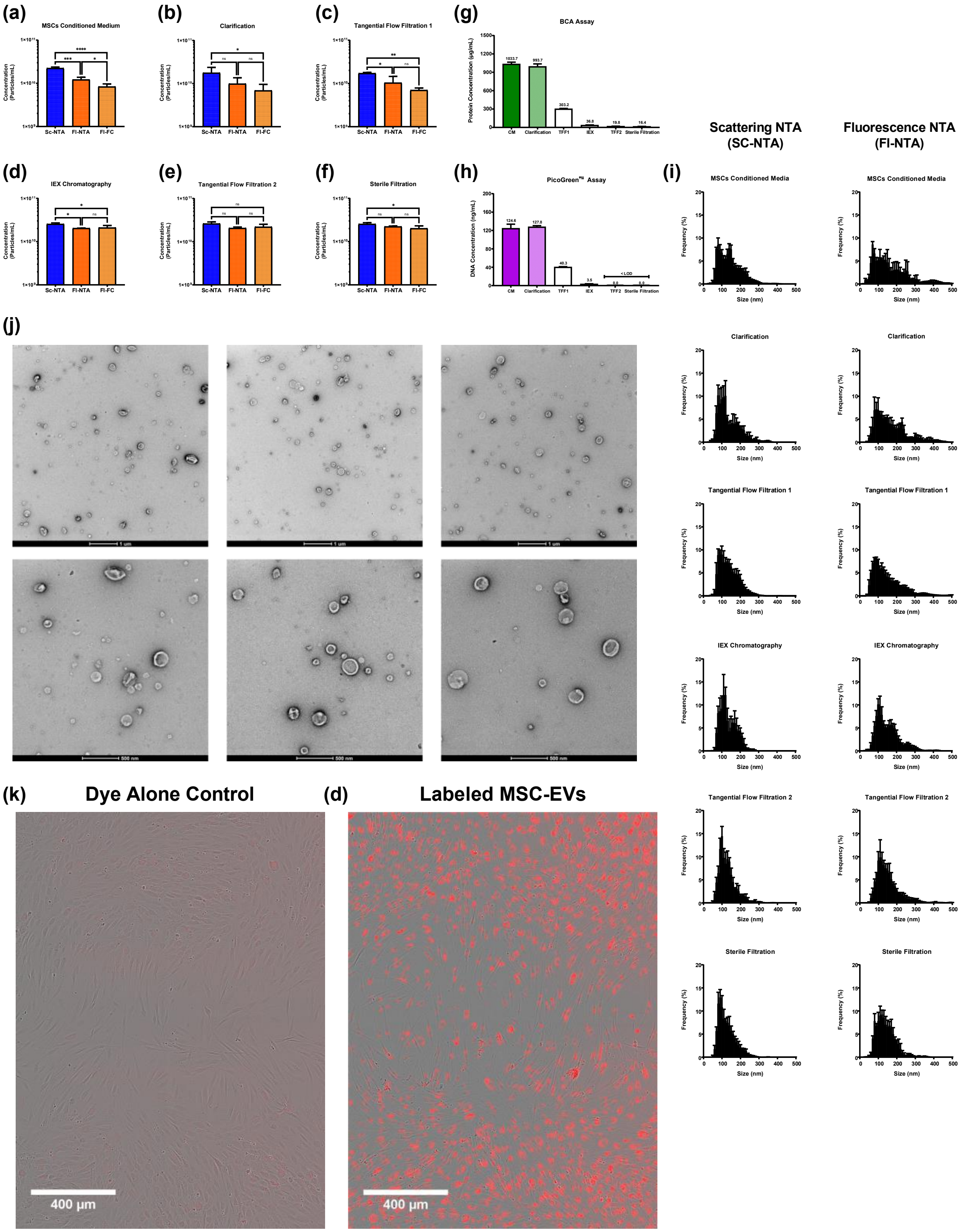


**Supplementary Figure 18. Downstream Processing (DSP) scalability;** Additional analytical assessment of particle recovery and impurity removal across the scaled DSP workflow for MSC-derived EVs, final MSC-EVs characterization and bioactivity evaluation.

Orthogonal analytical methods were applied to monitor particle, protein, and DNA content at each step of the scaled DSP process (700 mL feed volume). Particle recovery via Sc-NTA, Fl-NTA, and Fl-FC was measured for each DSP step; **(a)** MSCs conditioned medium (MSCs CM), **(b)** Clarification, **(c)** Tangential Flow Filtration 1, **(d)** Ion Exchange Chromatography (IEX), **(e)** Tangential Flow Filtration 2, and **(f)** Sterile filtration. Protein mass was measured using the BCA assay **(g)** while DNA content was determined using the PicoGreen™ assay **(f)**. **(i)** size distribution plots for each DSP step using SC-NTA and Fl-NTA. All data are presented as mean ± SD with n=3 (*p<0.05, **p<0.01, ***p<0.001, ****p < 0.0001; ns = not significant). **(j)** Additional representative TEM images at two different scale (scale bar: 1 µm and 500 nm). Uptake analysis of MSC-EVs in a fibroblast scratch would assay **(k)** dye alone control and **(l)** labeled MSC-EVs (scale bar: 400 µm). Sc-NTA: Scattering mode of NTA, Fl: Fluorescence mode of NTA, Fl-FC: Fluorescence-based flow cytometry.

### Fluorescence Labeling of MSC Derived EVs for Scratch Wound Healing Assay:

For EV uptake, the EV preparation was labeled with 80 µM ExoFluor Orange Dye (Sartorius) and incubated at 37° C for 30 mins. 100 kDA MWCO Vivaspin 2 columns (Sartorius) were used to wash the stained EV preparation and free dye control. Columns were pre-wet with PBS and centrifuged at 500 g for 5 mins. The stained EV preparation and free dye control were added to PBS in respective columns to bring the total volume to 2 mL. Columns were then centrifuged at 500 g for 5 mins to reduce the retentate volume to approximately 200 µL before washing the filter by pipetting up and down and adding PBS to increase the retentate volume back to 2 mL. This process was performed a total of 3 times, although after the final centrifugation the column was inverted and centrifuged at 200 g for 5 mins to fully collect the liquid from the retentate side of the filter. PBS was added to this retentate so that the final volume of stained EV preparation and dye only control was half of the initial volume to account for loss during washing. NTA verified labeled EV preparation particle concentration was at least as high as particle concentration prior to the labeling procedure (data not shown). 50 µL of labeled EV preparations and dye only controls + 200 µL of -CTL medium were used to treat hDFA for uptake visualization. All treatments used n=4-6 wells.

**Supplementary table 1.** Composition and pH of buffers used for EV isolation. Buffers were adjusted with 1M HCl and filtered through 0.22 µm sterile filters.

| | Wash buffer | Buffer A | Buffer B |
|---|---|---|---|
| NaCl [mM] | 0 | 0 | 1000 |
| TRIS [mM] | 20 | 20 | 20 |
| pH | 7.4 | 7.4 | 7.4 |

**Supplementary table 2.** Detailed information on all individual chromatography steps.

| Step | Volume (mL) | Flow rate [ml/min] | Solution |
|---|---|---|---|
| Flush | 15 | 15 | Buffer B |
| Equilibration | 60 | 15 | Buffer A |
| Sample Loading | 250 | 9 | MSCs CM |
| Wash | 60 | 9 | Wash Buffer |
| Elution 1 | 10 | 3 | 20% Buffer B |
| Elution 1 – Wash | 20 | 3 | 20% Buffer B |
| Elution 2 | 10 | 3 | 40% Buffer B |
| Elution 2 - Wash | 20 | 3 | 40% Buffer B |
| Linear | 10 | 3 | 40-100% Buffer B |
| Strip | 10 | 3 | 100% Buffer B |

**Supplementary table 3**. Detailed information on all individual chromatography steps – Scale up.

| Step | Volume (mL) | Flow rate [ml/min] | Solution |
|---|---|---|---|
| Flush | 100 | 100 | Buffer B |
| Equilibration | 400 | 100 | Buffer A |
| Sample Loading | 1800 | 60 | MSCs CM |
| Wash | 400 | 60 | Wash Buffer |
| Elution 1 | 80 | 20 | 20% Buffer B |
| Elution 1 – Wash | 120 | 20 | 20% Buffer B |
| Elution 2 | 80 | 20 | 40% Buffer B |
| Elution 2 - Wash | 120 | 20 | 40% Buffer B |
| Linear | 50 | 20 | 40-100% Buffer B |
| Strip | 50 | 20 | 100% Buffer B |
